# Homologous organization of cerebellar pathways to sensory, motor, and associative forebrain

**DOI:** 10.1101/2020.03.06.979153

**Authors:** Thomas J. Pisano, Zahra M. Dhanerawala, Mikhail Kislin, Dariya Bakshinskaya, Esteban A. Engel, Ethan J. Hansen, Austin T. Hoag, Junuk Lee, Nina L. de Oude, Kannan Umadevi Venkataraju, Jessica L. Verpeut, Freek E. Hoebeek, Ben D. Richardson, Henk-Jan Boele, Samuel S.-H. Wang

## Abstract

Cerebellar outputs take polysynaptic routes to reach the rest of the brain, impeding conventional tracing. Here we quantify pathways between cerebellum and forebrain using transsynaptic tracing viruses and a whole-brain quantitative analysis pipeline. Retrograde tracing found a majority of descending paths originating from somatomotor cortex. Anterograde tracing of ascending paths encompassed most thalamic nuclei, especially ventral posteromedial, lateral posterior, mediodorsal, and reticular nuclei; in neocortex, sensorimotor regions contained the most labeled neurons, but higher densities were found in associative areas, including orbital, anterior cingulate, prelimbic, and infralimbic cortex. Patterns of ascending expression correlated with c-Fos expression after optogenetic inhibition of Purkinje cells. Our results reveal homologous networks linking single areas of cerebellar cortex to diverse forebrain targets. We conclude that shared areas of cerebellum are positioned to provide sensory-motor information to regions implicated in both movement and nonmotor function.

**Graphical Abstract:** 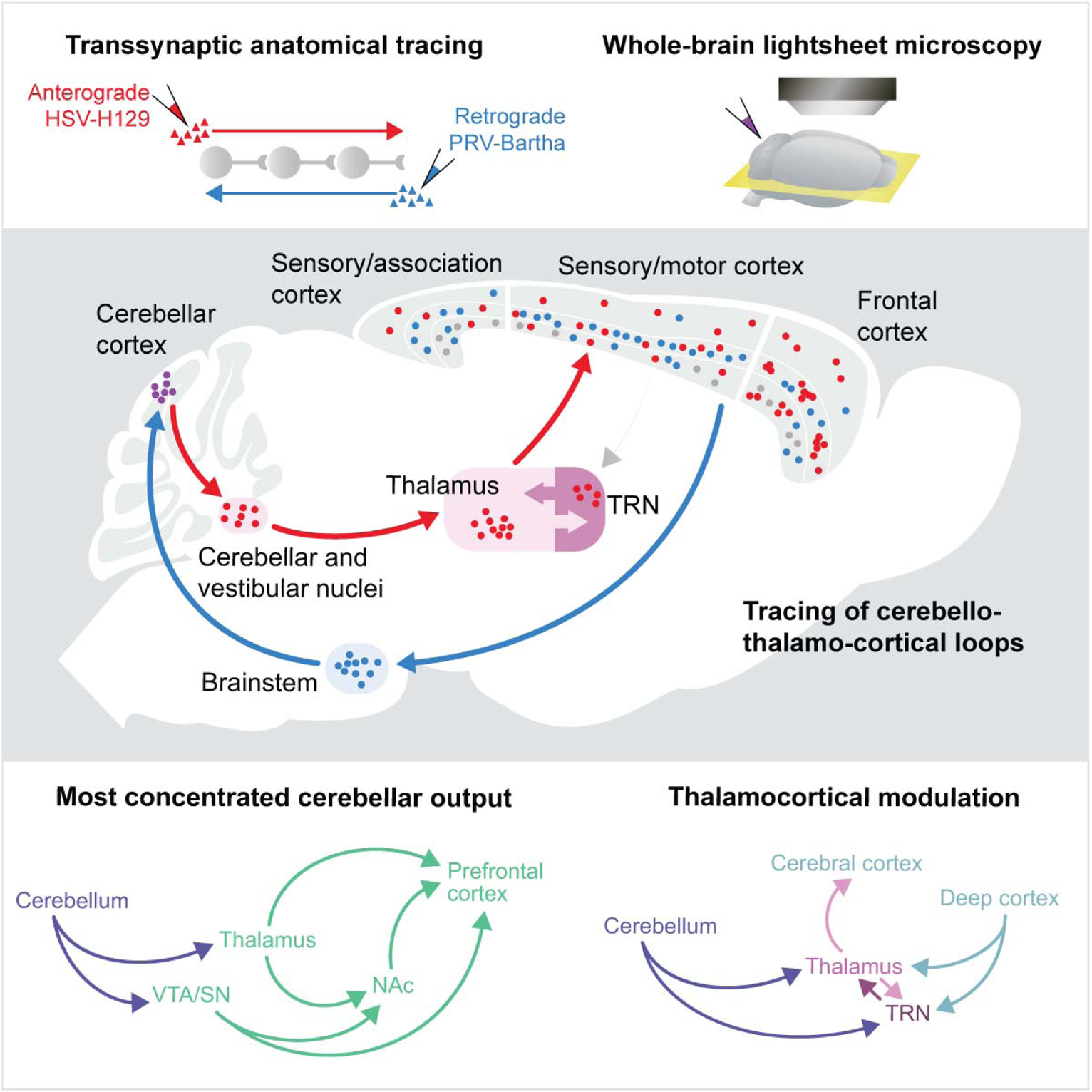

## Introduction

The cerebellum has an increasingly recognized role in nonmotor processing (Badura et al., 2018; Deverett et al., 2018; Stoodley and Schmahmann, 2009). Patients with cerebellar damage not only show motor symptoms, but also suffer from multiple cognitive and affective symptoms (Stoodley and Schmahmann, 2009). Cerebellar damage at birth leads to autism spectrum disorder (ASD) in almost half of cases (Cook et al., 2020; Courchesne et al., 2001; Limperopoulos et al., 2007; Wang et al., 2014). These observations suggest a broad role for the cerebellum in both motor and nonmotor function during development and adulthood. However, the whole-brain pathways mediating these nonmotor influences are poorly characterized.

Monosynaptic inputs and outputs of cerebellum are well-mapped (Apps and Hawkes, 2009; Sugihara and Shinoda, 2004; Suzuki et al., 2012; Voogd and Ruigrok, 2004). But classical anatomical methods cannot trace polysynaptic connections (Ugolini, 2010). Furthermore, characterization of individual projections does not characterize the overall pattern of cerebellar influence on the brain. Long-range connections between cerebellum and cerebrum have been studied using functional MRI and transcranial magnetic stimulation (Buckner et al., 2011; Choe et al., 2018; Popa et al., 2010), which do not provide cellular-resolution information.

We sought to provide whole-brain quantification of anatomical pathways between cerebellum and forebrain using cellular tracing methods. We used retrograde and anterograde transsynaptic viral tracers combined with brain clearing and whole-brain light-sheet microscopy to allow neuron-level analysis. To maximize anatomical accuracy, we generated a whole-brain atlas with an entire cerebellum, where the standard Allen Brain Atlas omits the posterior two-thirds. Because cerebellar lesions affect both motor and nonmotor function, we hypothesized that cerebellum forms long-range connections with both sensorimotor and nonmotor areas of thalamus and neocortex.

The resulting brain volumes required computationally efficient cell detection using machine learning and anatomical assignment using image registration to align brains. We generated a bidirectional cerebellum-to-forebrain map and confirmed the impact of ascending paths using optogenetic stimulation of c-Fos expression.

## Results

### HSV-H129 transsynaptic viral labeling reveals distant cerebellar targets

The canonical ascending cerebellum-neocortical circuit begins with Purkinje cells (PCs) in the cerebellar cortex that project to deep cerebellar nuclei (DCN) which project via intermediates like thalamus to neocortex. To trace anterograde transsynaptic paths from neocerebellum, we used HSV-H129-VC22 (**Figure 1a,b**), an anterograde-transported herpes simplex virus (HSV) strain that expresses nuclear-targeted enhanced green fluorescent protein (EGFP). Transsynaptic viral tracing yields weaker labeling than longer-expression-time strategies such as AAV. To achieve a high signal-to-noise ratio we used iDISCO+ (Renier et al., 2016), which allows for whole-brain immunostaining without differential depth loss, followed by tissue clearing and light-sheet microscopy (**Figure S13d**).

**Figure 1.**
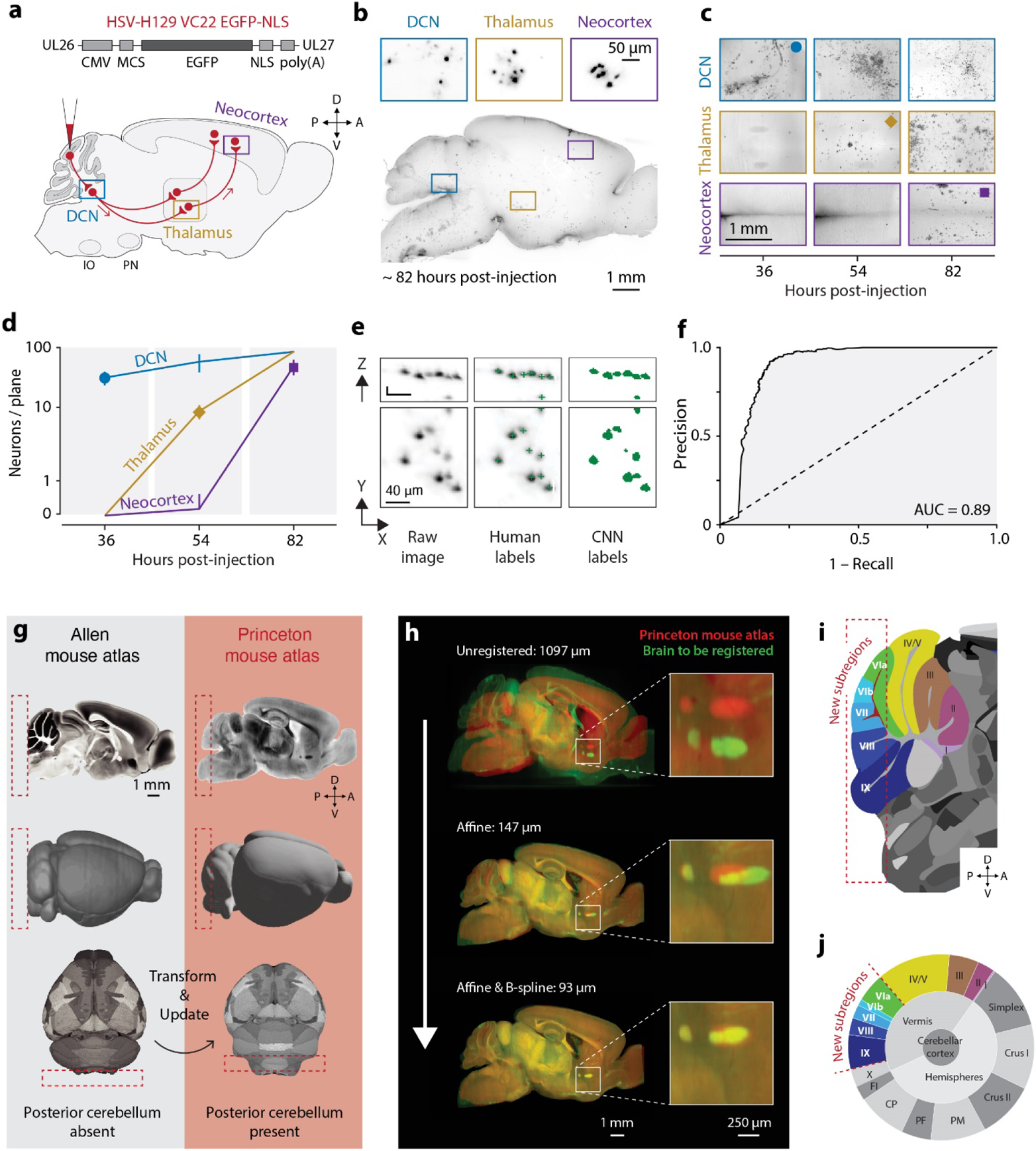
Large-scale transsynaptic tracing with tissue clearing, light-sheet microscopy and registration to the Princeton Mouse Brain Atlas. (a) *Top,* H129-VC22 expresses a nuclear location signal tagged with eGFP. *Bottom,* experimental design to trace pathways from cerebellar cortex to thalamus and neocortex. (b) Images of an iDISCO+ processed brain 82 hours post-HSV-H129 injection. 158 µm maximum intensity projections (MIP). (c) Time course of infection. Horizontal MIPs of DCN (3.0 mm dorsal of bregma), thalamus (3.0 mm dorsal), and neocortex (0.7 mm dorsal). Dorsoventral depth: 300 µm for DCN and thalamus, 150 µm for neocortex. (d) Quantification of viral spread.Cell counts from five planes at each timepoint for each brain region. Error bars show95% confidence interval. (e) Training data for convolutional neural network (CNN). *Left,* raw input data. *Middle,* human-annotated cell centers (green) for training the network. *Right,* segmented labels (green) used as training input. (f) Receiver operating characteristic curve for the trained CNN. The diagonal line indicates chance performance. (g) Differences between Allen Brain Atlas (ABA, left) and the Princeton Mouse Brain Atlas (PMA, right). The red dotted box indicates the ABA’s caudal limit. ABA annotations were transformed into PMA space. (h) Registration of whole-brain light-sheet volumes to the PMA. Individual brain (green) overlaid with PMA (red) at different stages of registration, with median discrepancy shown for each stage of alignment. (i) PMA cerebellar annotations. Red dotted box indicates updated annotated areas. (j) PMA cerebellar hierarchy showing relative substructure size contributions. Abbreviations: AUC, area under curve; PM, paramedian lobule; PF, paraflocculi; CP, copula pyramidis; Fl, flocculus. Interactive online atlas and injection site segmentations available at: https://brainmaps.princeton.edu/2021/05/pisano_viral_tracing_injections/.

We defined 54 hours post-injection (hpi) as the disynaptic (*e.g.* PC to DCN to thalamic) timepoint and 80 hours post-injection as a trisynaptic timepoint to reach neocortex (**Figure 1c,d**; see *Methods, Transsynaptic timepoint determination*). Our timepoints are consistent or shorter than other studies (**Table S61**) (Badura et al., 2018; Song et al., 2009), suggesting decreased risk for further spread.

At longer incubation times, H129 has a slow retrograde component (Wojaczynski et al., 2015) which might occur via uptake by axon terminals (Su et al., 2019) followed by transport backward across synapses. To test this, we examined dorsal column nuclei, which are two synaptic steps retrograde from the cerebellar cortex (via mossy fibers) but do not receive anterograde paths via DCN or vestibular nuclei (**Figure S1**). At 28-36 hpi, we observed minimal labeling in dorsal column nuclei. The ratio of retrograde to anterograde (DCN) cell density, which normalizes for injection size, was 0.084 ± 0.032 at 28-36 hpi (median ± estimated SEM, n=15 locations in 5 mice), and 0.096 ± 0.017 at 54 hpi (n=69 locations in 23 mice). To demonstrate the maximum possible retrograde labeling we used PRV (80 hpi) and found a median retrograde:anterograde density ratio of 2.06 ± 0.17 (n=75 locations in 25 mice). Finally, we ascertained where any retrograde H129 viral uptake would lead via subsequent anterograde spread by examining the Mouselight database (Winnubst et al., 2019). We found 36 brainstem neurons with at least one cerebellar-projecting axon. In all but one neuron, the cerebellum was the sole target (**Figure S1f,g**). Thus, to the extent that retrograde transport from injection sites occurs, it would still not lead to alternate noncerebellar pathways. We conclude that at our selected timepoints, H129 acts as an anterograde tracer (Zemanick et al., 1991).

### Generation of the Princeton Mouse Atlas

For image registration, we devised a two-step procedure to calculate an averaged light-sheet-based brain template for referral to the volumetric Allen Brain Atlas (ABA; **Figure 1g-j**). The ABA, a field standard, is based on serial two-photon microscopy and lacks a complete cerebellum (**Figure 1g**). We constructed a Princeton Mouse brain Atlas (PMA; **Figure S2a-d, Video S1**) by computing a transform between our averaged light-sheet template and ABA CCFv3 space (**Figure 1h**), then extending ABA-labels using manually-drawn contours that included complete posterior lobules (**Figure 1i,j**; **Figure S2e-h, Video S2**). The estimated accuracy of registration was 79 µm, or 4 voxels (see *Quantification and Statistical Analysis*).

### The cerebellum sends output to a wide range of thalamic targets

We used our automated analysis pipeline, which we named BrainPipe, to quantify cerebello-thalamic connectivity (**Figure 1e,f and 2a**; see *Methods, Automated detection of virally labeled cells* and *Statistical analysis of transsynaptic tracing data*). We injected 23 brains with H129 at different sites in the cerebellum (**Figure 2b and S3**) and collected brains at the disynaptic (54 hpi) timepoint. Labeled neurons per region were widely distributed among contralateral thalamus (**Figure 2c**). Neuron density in neocortical regions was 0.085 ± 0.073 (mean ± standard deviation, 17 regions) times that seen at 80 hpi, indicating sufficient transport to thalamus but not neocortex. Counts by region were not systematically related to anteroposterior position (rank correlation with anteroposterior position r=+0.05), suggesting that labeling efficiency did not depend on transport distance. Counts also did not diminish with depth along the light path (**Figure S13d**). For display, neuron counts for each region were converted to percentage of total per-brain thalamic neurons and coded as “sensory/motor” and “polymodal association” based on ABA ontology (**Figure 2**).

**Figure 2.**
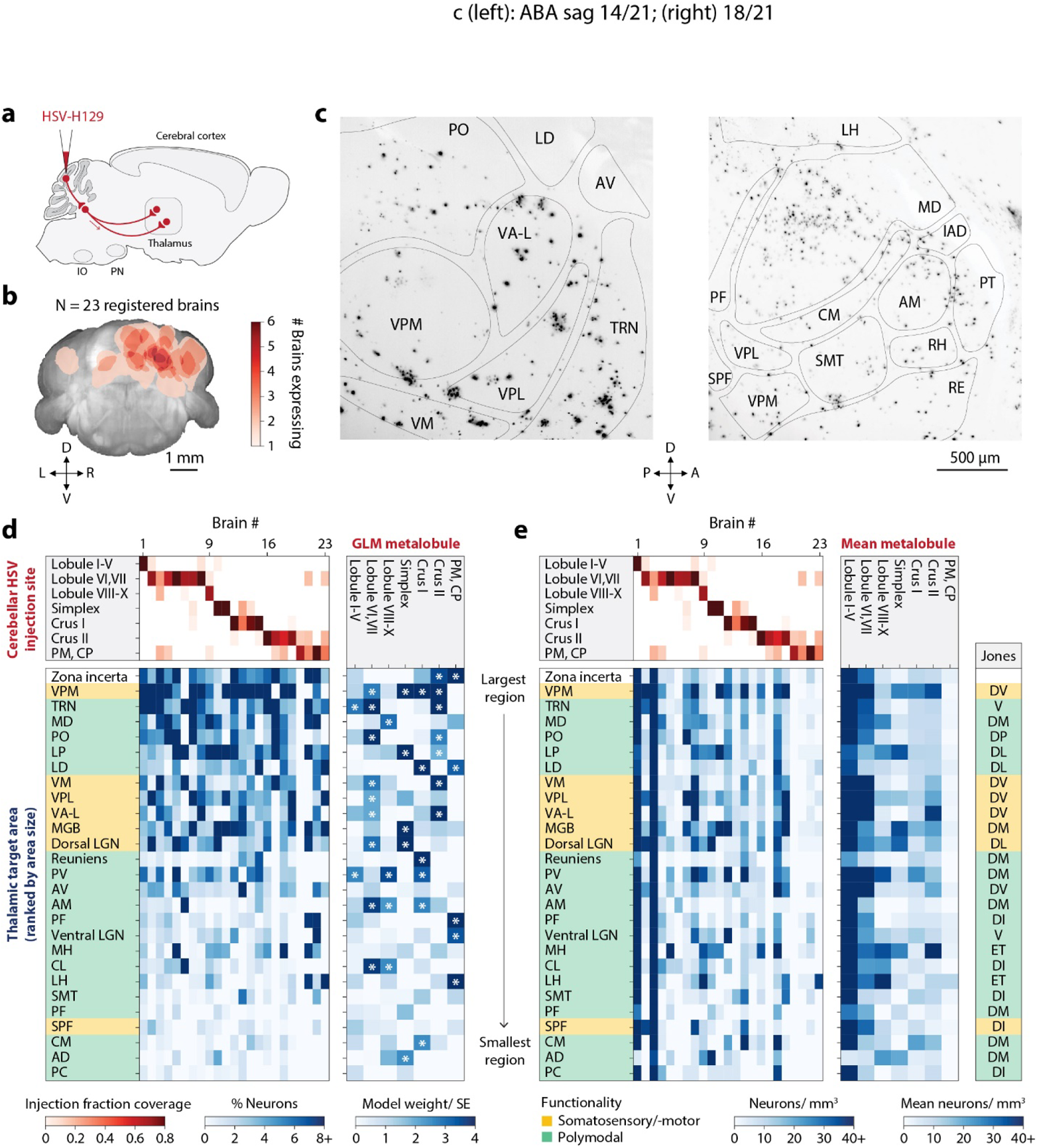
Cerebellar paths to thalamus. (a) Disynaptic H129 tracing from the cerebellar cortex to thalamus. (b) Coverage of cerebellum by thalamic timepoint injections marked by CTB-Alexafluor555. (c) Maximum intensity projections (150 µm) of anti-HSV primary and anti-rabbit Alexa Fluor 647 secondary immunolabeling. (d) *Left*, fraction of neurons across all injection sites, (area countdivided by total thalamic count). Injection coverage fractions are shown in red, fraction of neurons in blue. Each column represents one mouse. *Right*, a generalized linear model showing the influence of each cerebellar region on thalamic expression. The heatmap (blue) shows coefficient divided by standard error. Significant coefficients are marked with asterisks. (e) *Left*, neuron density in each thalamic area across all cerebellar injection sites. *Middle*, mean density across injection sites by cerebellar region. *Right*, grouping according to Ref. (Jones, 2012). Abbreviations: AD, anterodorsal; AM, anteromedial, AV, Anteroventral; IAD, interanterodorsal; CL, central lateral; CM, central medial; LD, lateral dorsal; LGN, lateral geniculate nucleus; LH, lateral habenula; MH, medial habenula; PC, paracentral; PF, parafascicular; PO, posterior complex; PT, paratenial; PV, paraventricular; RE, reuniens; RH, rhomboid; SMT, Submedial; SPF, subparafascicular; TRN, thalamic reticular nucleus; VA-L, ventral anterior-lateral; VPL, ventral posterolateral; VPM, ventral posteromedial.

The cerebellothalamic tract originates from the DCN and ascends through the superior cerebellar peduncle (also known as brachium conjunctivum), with most axons crossing the midline before reaching the thalamus. We observed labeling in vestibular nuclei, consistent with a direct projection from cerebellar cortex (**Figure S4a**). Short-incubation experiments showed that vestibular nuclei contained 22% of the total combined vestibular and DCN cell count. To assess locations of PCs that project to vestibular nuclei, we examined the MouseLight database (Winnubst et al., 2019) and found 9 PCs with direct vestibular-projecting axons, of which 6 were in non-flocculonodular regions (**Figure S4a**). Thus a substantial fraction of PCs that project to vestibular nuclei arise from non-flocculonodular lobules.

A principal target of cerebellothalamic axons is the ventral nuclear group (Gao et al., 2018; Teune et al., 2000), which includes sensorimotor nuclei (Jones, 2012). Consistent with known DCN projections, we observed strong connectivity to ventromedial (VM) and ventral anterior-lateral (VA-L), motor thalamic nuclei (Sieveritz et al., 2019), from vermal lobules (I-VII) and crus II, and moderate connectivity from crus I (**Figure 2d,e**). The ventral posteromedial nucleus (VPM), which conveys whisker and mouth information, received input from all but the most posterior cerebellum (**Figure 2c,d,e**) consistent with known interpositus and vestibular nuclear projections (Aumann et al., 1994; Wijesinghe et al., 2015). Our findings confirm that cerebellar-injected H129 labels major known pathways to neocortex via multiple distributed DCN, vestibular, and thalamic intermediates.

We also observed labeling outside the ventral nuclei, including the thalamic reticular nucleus (TRN), two association nuclei (lateral posterior, LP and mediodorsal, MD), primary relay nuclei (LGN and MGN), and zona incerta (ZI) (Ossowska, 2020). MD is engaged during reversal learning (Mitchell and Chakraborty, 2013), sends its output to frontal regions (Hunnicutt et al., 2014), and is engaged in cognitive and working memory tasks in humans (Mitchell and Chakraborty, 2013). Lobule VI, a site of structural abnormality in ASD (Courchesne et al., 1988), made dense projections to MD (**Figure 2d,e**). These results suggest a strong role for cerebellum in flexible cognition. LP sends its output to visual, sensorimotor and frontal association cortex (Hunnicutt et al., 2014). TRN, unlike other thalamic nuclei, does not project to the neocortex, but instead sends inhibitory projections to other thalamic nuclei. Thus, the cerebellum has anatomical capabilities to influence both relay nuclei and the other two major classes of nuclei, association (MD, LP) and local modulatory (TRN).

We fitted a generalized linear model (GLM; **Figure 2d**) using the fraction-by-lobule of total cerebellar injection as input, and the fraction-by-nucleus of total thalamic expression as output. In this way the GLM can identify topographical relationships shared across animals. The GLM revealed a broad mapping of lobules I-X to diverse thalamic targets, and a more focused pattern of mapping from simplex, crus I and II, paramedian lobule (PM), and copula pyramidis (CP). Mapping hotspots included lobules I-X to VPM, TRN, MD, VM, VPL, VA-L, paraventricular (PV), anteromedial, and centrolateral (CL); simplex to VPM, LP, dLGN, MG, and anterodorsal; crus I to VPM, lateral dorsal (LD), reuniens, PV and anteromedial and central lateral; crus II to ZI, VPM, MD, TRN, VA-L, posterior complex, and LP; and PM and CP to ZI, LD, parafascicular, vLGNe, and lateral habenula (**Table S2**).

### Direct projections from DCN to thalamus are largely consistent with transsynaptic tracing

As a second, non-transsynaptic approach to characterizing cerebellar projections to thalamus, we injected GFP-expressing AAV into DCN and characterized the spatial distribution of labeled terminals (**Figure 3**). Injections primarily targeted bilateral dentate and also reached interposed and fastigial nuclei (**Figure 3a; Figure S4b**).

**Figure 3.**
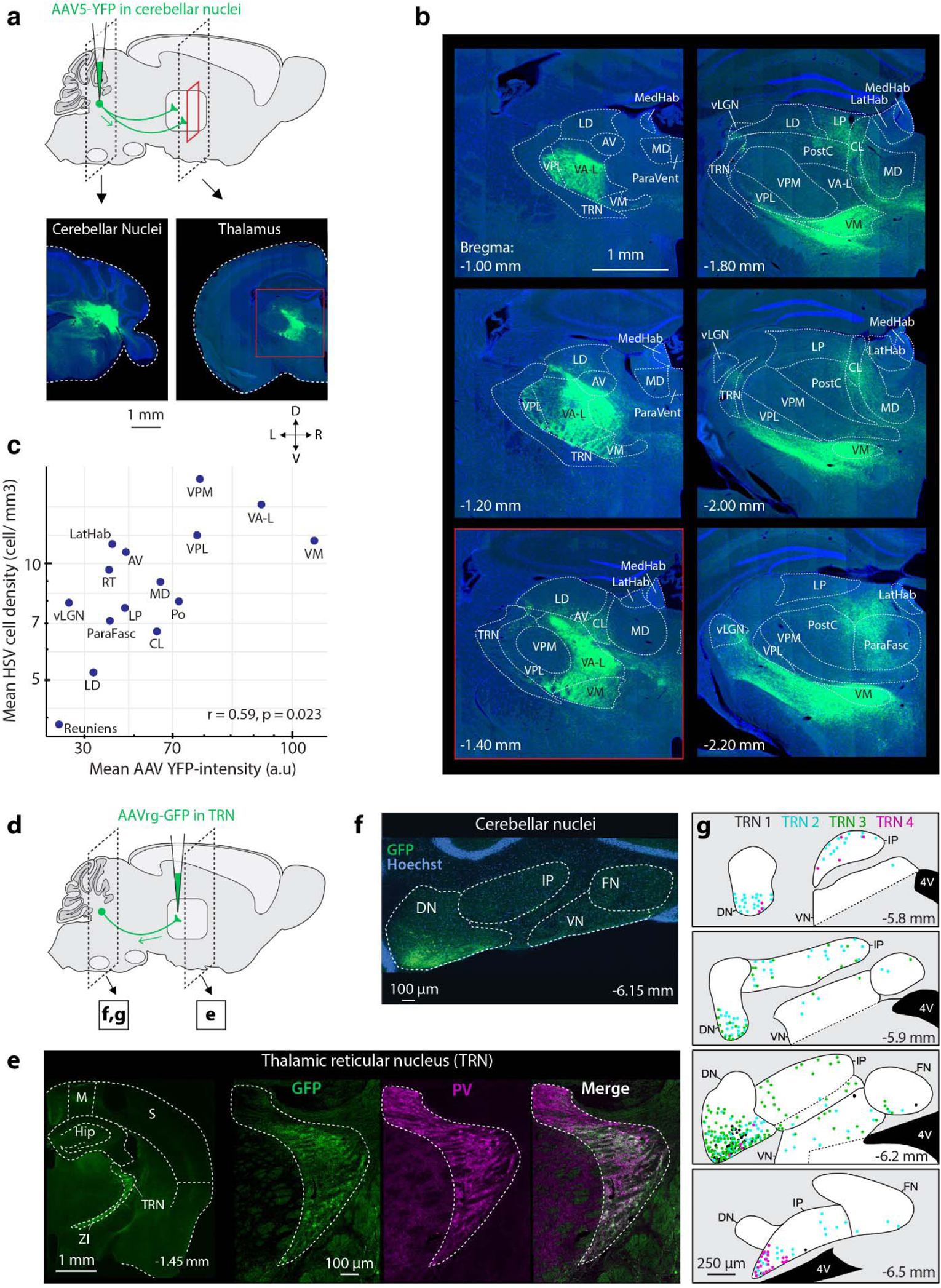
Cerebellothalamic AAV-identified axons match transsynaptic tracing, including the thalamic reticular nucleus. (a) Physical sectioning after AAV-YFP DCN injections to visualize cerebellothalamic axon projection density. (b) DCN injection primarily targeted the interpositus and dentate. Manually drawn Paxinos coronal overlays are shown. Bregma −1.40 mm corresponds to A. (c) Pearson’s correlation (r=0.59, p=0.023) of rank order density of HSV-labeled thalamic neurons after cerebellar cortical injection versus cerebellothalamic axonal projection density. (d) AAVrg-GFP injection into right TRN. (e) Confocal image of GFP (green) labeling of a major subset of neurons in TRN (white outline). (f) In left DCN, presence of GFP-expressing (green) neuron bodies primarily in ventrolateral dentate with additional expression in dorsal IP and fastigial nuclei. Nuclei are labeled with Hoechst (blue). (g) GFP+ cell body distribution (dots) in DCN quantified from 4-5 coronal cerebellar sections per animal. Dots are color-coded to individual experiments. Abbreviations: 4V, fourth ventricle; AV, anteroventral; CL, central lateral; DN, dentate nucleus; FN, fastigial nucleus; GFP, green fluorescent protein; Hip, hippocampus; IP, interpositus nucleus; LatHab, lateral habenula; LD, lateral dorsal; LP, lateral posterior; M, motor cortex; MD, mediodorsal; ParaFasc, parafascicular; Po, posterior complex; PV, parvalbumin; S, somatosensory cortex; TRN, thalamic reticular nucleus; VA-L, ventral anterior-lateral; VM, ventral medial; VN, vestibulocerebellar nucleus; VPL, ventral posterolateral; VPM, ventral posteromedial; vLGN, ventral lateral geniculate nucleus; ZI, zona incerta.

Terminals were clearly visible throughout thalamus (**Figure 3b**), largely contralateral to the injection site. Ventral thalamic nuclei, including VM, VA-L, VPM, and VPL, showed the most signal, consistent with previous reports and with our H129 cell density findings. Within-nucleus fluorescence density (summed brightness divided by the total area covered by the nucleus) was correlated with H129 neuron density averaged across injections (**Figure 3c**; log-log correlation r=+0.59, p=0.023). Taken together, these measurements indicate that H129 injections capture representative DCN-thalamic connectivity.

### DCN project directly to the reticular thalamic nucleus

To identify neurons that project directly to TRN, we injected into the TRN an AAV (AAVrg-hSyn-Chronos-GFP) that infects presynaptic terminals and moves retrogradely along axons to the parent cell body (Klapoetke et al., 2014; Tervo et al., 2016). We observed expression in the contralateral ventrolateral dentate and dorsolateral interpositus nuclei (**Figure 3d-g and S5**), consistent with prior reports (Angaut et al., 1968; Cavdar et al., 2002; Chan-Palay, 2013; Nakamura, 2018) (**Figure S6**). One injection missed TRN, and instead infected the nearby internal capsule (**Figure S5**), and did not label DCN. These findings are consistent with a disynaptic projection from cerebellar cortex to TRN as found by H129 injection.

### Cerebellar paths to neocortex are proportionally greatest to somatomotor regions and densest in frontal regions

To characterize cerebellar paths to neocortex, we examined 33 H129-injected brains at 80 hpi (**Figure 4a,b,c**). As expected, the majority of contralateral neocortical neurons were found in somatosensory and somatomotor cortex, with additional neurons at more anterior and posterior locations (**Figure 4d**). No clear differences among subregions of somatosensory and somatomotor areas were identified (**Figure S7a,b**).

**Figure 4.**
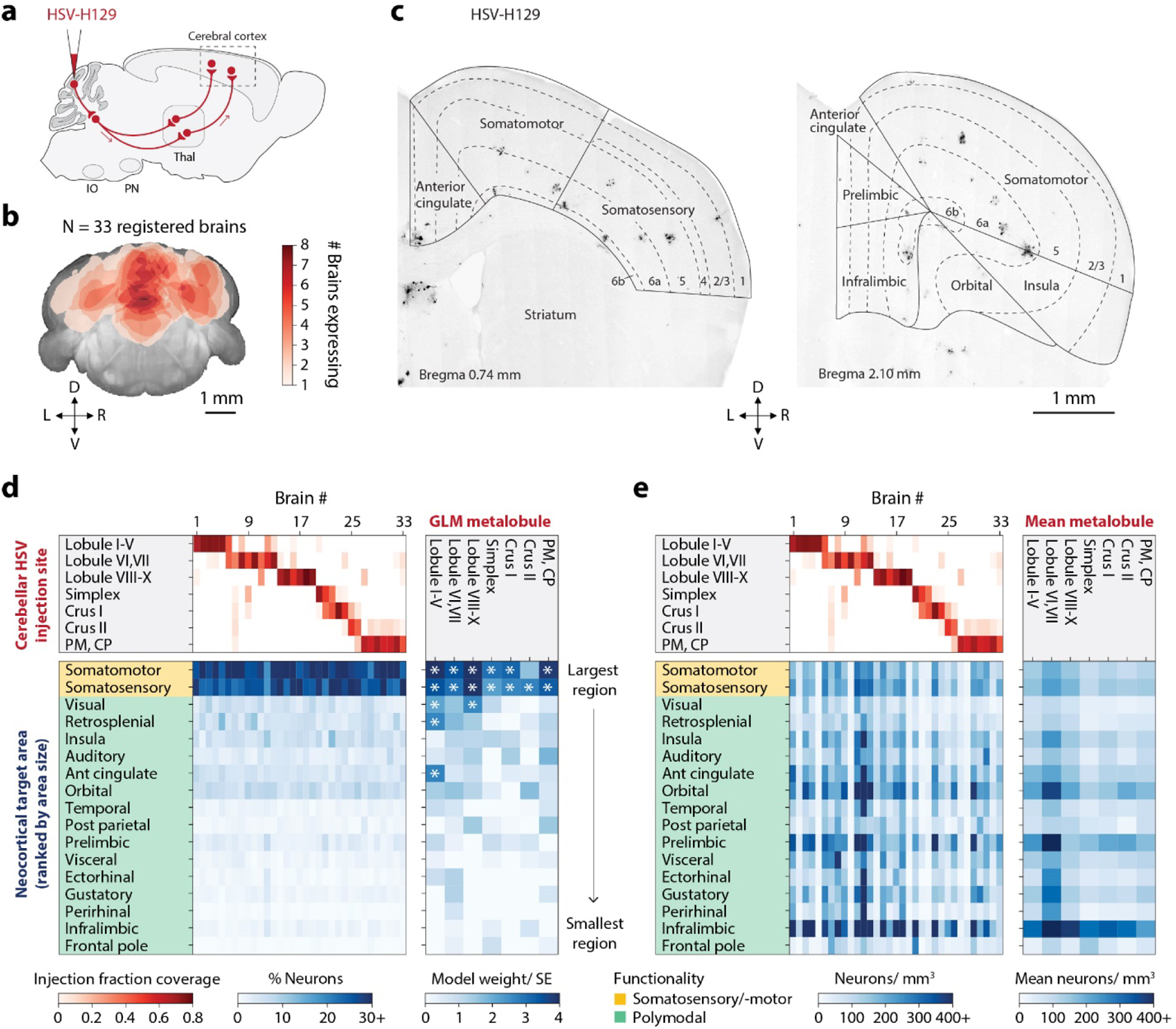
Cerebellar paths to neocortex. (a) H129 injections traced trisynaptic paths from cerebellar cortex to neocortex. (b) Coverage of cerebellum by neocortical timepoint injections marked by CTB-Alexafluor555. The number of injections covering each cerebellar location is shown. See also https://brainmaps.princeton.edu/2021/05/pisano_viral_tracing_injections/. (c) Maximum intensity projections (100 µm) with outlines defining neocortical structures. (d) *Left*, fraction of neurons in each neocortical area across injection sites. Injection coverage fractions (red) and neuron fraction (blue) for each brain, ordered by primary injection site. *Right*, generalized linear model showing cerebellar area influence on neocortical expression. The heatmap (blue) shows the coefficient divided by standard error. Significant coefficients are marked with asterisks. (e) *Left*, neuron density in each neocortical area across injections. *Right*, mean neuron density. Abbreviations: Ant, anterior; CP, Copula pyramidis; PM, Paramedian; Post, posterior.

When counts were converted to projection density by region, a different pattern emerged (**Figure 4e**). Neuron densities were highest in contralateral anterior and medial neocortical regions, with peak regions exceeding 400 neurons per mm^3^, more than twice the highest density found in somatosensory and somatomotor regions. Labeling was dense in infralimbic, orbital, and prelimbic areas (**Figure 4e**).

To build a single map from many injections, we fitted a GLM to the data in the same way as for thalamic labeling (**Figure 4d**). All injected cerebellar sites showed high weights in somatomotor and somatosensory cortex. Lobules I-V also showed significant weights in the anterior cingulate cortex. The visual and retrosplenial cortex showed weak clusters of connectivity. Mean density by primary injection site (**Figure 4e**) revealed that all sites sent dense projections to the infralimbic cortex. Vermal lobules VI-X and crus I sent denser projections than other sites to infralimbic, prelimbic, and orbital cortex (**Figure 4e**). A similar pattern was observed by taking the maximum of the fraction of neurons across each cerebellar region, where the majority of neurons were found in somatosensory and somatomotor cortex and a smaller number in retrosplenial, agranular insular, anterior cingulate, and orbital cortex (**Figure S7c,d**).

### Cerebellar paths reach reward-based structures in striatum and hypothalamus and project modestly to ventral tegmental area

Among monosynaptic targets of the DCN, renewed focus has fallen on the ventral tegmental area (VTA) (Phillipson, 1979; Watabe-Uchida et al., 2012), including cerebellar influence on reward processing (Carta et al., 2019). Using our anterograde data, we compared the relative projection strengths of contralateral cerebellar paths to thalamus and two midbrain dopaminergic areas, VTA and the substantia nigra (**Figure S8a**). Contralateral VTA (Snider and Maiti, 1976) counts were considerably lower than in thalamic regions, consistent with literature (Aumann et al., 1994; Carta et al., 2019; Phillipson, 1979). Normalized to density per unit volume of the target region, VTA projections were less than one-third as strong as projections to VPM, MD, and TRN. Densities in substantia nigra were even lower than in VTA. In summary, cerebellar projections to VTA constituted a moderate-strength projection, weaker than thalamic targets but greater than other dopaminergic targets.

Striatal regions are also involved in reward learning. The cerebellar cortex is known to project to basal ganglia trisynaptically via the DCN and thalamus (Bostan and Strick, 2018; Fujita et al., 2020). Among striatal regions, at our trisynaptic timepoint, we observed the most labeling in the caudate, nucleus accumbens (NAc), and cortical amygdala. Labeling was dense in NAc, septohippocampal and septofimbrial nuclei as well as central/medial amygdala (**Figure S8b**). At the di- and tri-synaptic timepoints we also quantified hypothalamic connectivity observing relatively strong expression in the lateral area and the periventricular nucleus. Projection density was highly variable, likely related to the small volumes of hypothalamic nuclei (**Figure S9**). At both timepoints, we observed strong labeling in the lateral hypothalamic area, which has been shown to regulate feeding and reward (Stamatakis et al., 2016) and the zona incerta, a well-established recipient of DCN output (Fujita et al., 2020).

### Cerebellum-neocortical paths strongly innervate deep neocortical layer neurons

To investigate the layer-specific contributions of cerebellar paths to neocortex, we examined trisynaptic-timepoint laminar expression (**Figure 5**). To minimize near-surface false positives, 60 µm was eroded from layer 1. In most neocortical areas, we found the most and densest anterogradely labeled neurons in layers 5, layers 6a and 6b (**Figure 5b,c**). No differences in layer-specific patterns were apparent from injections to anterior vermis, posterior vermis, and posterior hemisphere (p>0.95, ANOVA, two-tailed, 3 injection groups).

**Figure 5.**
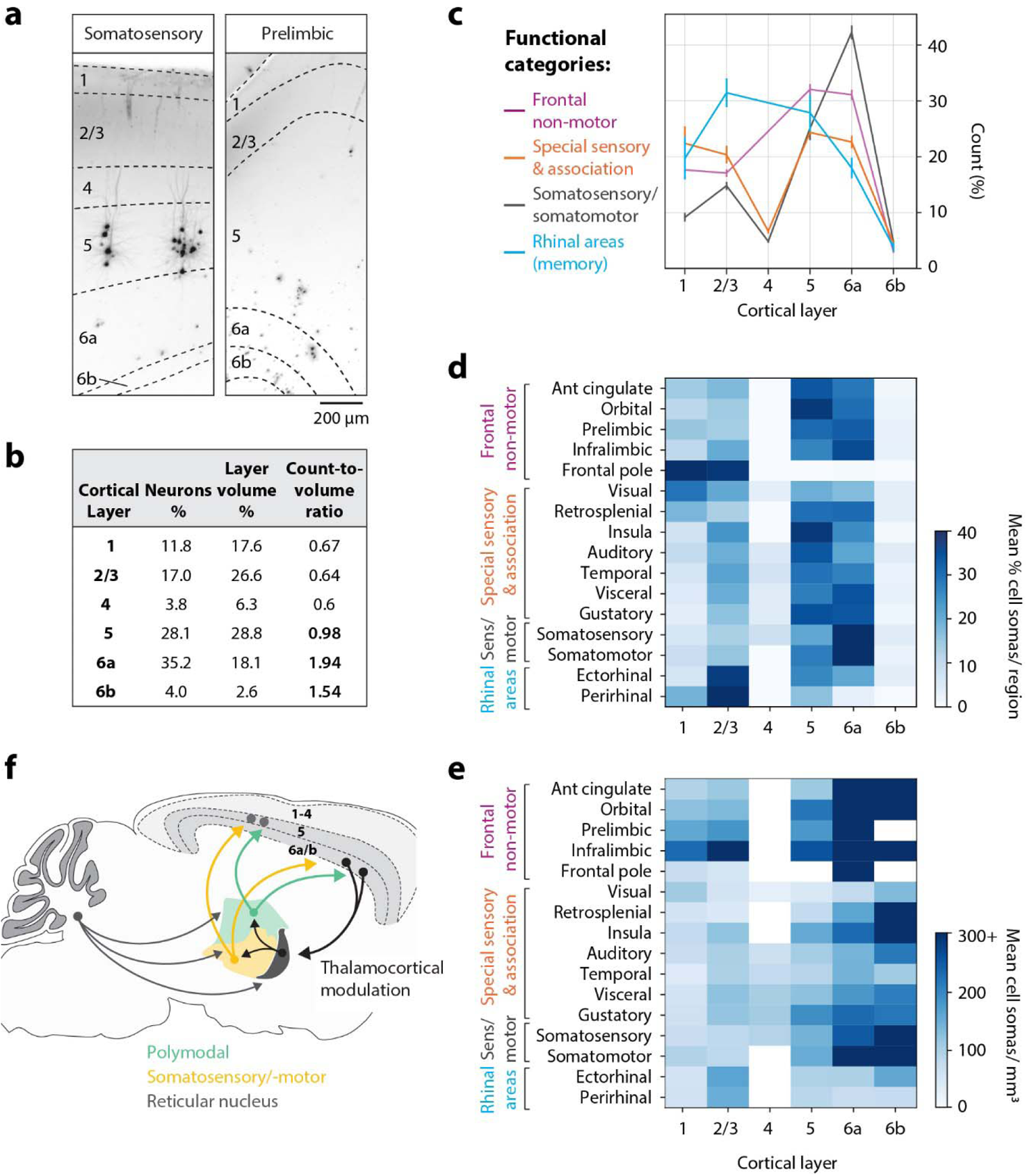
Cerebellar projections to thalamocortical and deep-layer modulatory systems. (a) Neocortical labeling with outlines depicting layers. 75 µm maximum intensity projections. (b) Distribution of neocortical neurons by layer. (c) Projection differences by layer after grouping cortical regions by function. Percent of cortical neurons shown. Error bars are 95% confidence intervals n=33. (d) Average fraction of neurons per region and (e) average density for neocortical regions separated by layer. Neocortical regions are functionally grouped as in (c). (f) Summary of cerebellar output connectivity to thalamus and neocortex. Abbreviations: Ant, anterior. Functional categories: Rhinal: perirhinal and ectorhinal areas; Somatosensory/-motor: somatosensory and somatomotor areas; Special sensory & association: retrosplenial, visual, gustatory, visceral, auditory, and temporal association areas; Frontal nonmotor: infralimbic, anterior cingulate, orbital, prelimbic, agranular insular areas.

Layer-specificity of thalamocortical connections varies by neocortical region (Jones, 1975; Jones and Burton, 1976). A common motif of thalamocortical projections is strong innervation of layer 6 neurons, especially in sensory regions (Constantinople and Bruno, 2013; Herkenham, 1980; Thomson, 2010). In sensorimotor regions (somatomotor and somatosensory), over 40% of labeled cells were in layer 6, a higher fraction than in other categories of neocortex (**Figure 5b**). To validate these findings, we injected H129 in Thy1-YFP mice, which express YFP primarily in layer 5. These injections revealed viral labeling in neocortex subjacent to YFP (**Figure S10a-c**).

Layer 4 of sensory regions receives thalamic innervation (Herkenham, 1980). However, classical tracing typically does not identify the cellular target, only the cortical layer where synapses occur (Hooks et al., 2013). We found that labeled layer 4 neurons comprised only 10% of cells in somatosensory cortex and even less in other sensory regions (gustatory, visceral, temporal, visual). Our results are consistent with the fact that although thalamocortical synapses often occur in a more superficial layer (layer 4), the recipient postsynaptic cell body resides in deeper layers (layer 5 or 6) (Llinás et al., 2002).

A different pattern was seen in rhinal cortex, part of the medial temporal system for declarative memory. Rhinal regions (perirhinal, ectorhinal, and entorhinal) had the highest fraction of layer 2/3 neurons (**Figure 5c,d,e**). This finding recalls the observation that in associative neocortical regions, thalamocortical axons send substantial projections to superficial layers (Thomson, 2010). Frontal and other association regions showed patterns that were intermediate between sensorimotor and rhinal regions, while infralimbic, prelimbic, orbital, and anterior cingulate cortex also received more and denser projections to layer 1 (**Figure 5c,d,e**). The share of labeling found in layer 5 and 6 neurons was higher for frontal nonmotor regions than for other cortical areas. Taken together, our analysis reflects past findings that thalamic influences on neocortex arrive directly through superficial and deep layer pathways (Llinás et al., 2002) (**Figure 5f**).

### Pseudorabies virus reveals strong descending input from somatomotor and somatosensory areas

PCs receive principal input from two extracerebellar sources, the inferior olive and the pons, which receive input from ascending (spinal cord and brainstem; **Figure S14a**) and descending (neocortical, mesodiencephalic junction, and other) sources (Mihailoff et al., 1989). Ascending and descending cerebellar input converge on individual microzones (Apps and Hawkes, 2009; Kubo et al., 2018). To characterize disynaptic paths from neocortex to cerebellum, we performed a series of cerebellar injections of pseudorabies virus (PRV) Bartha strain, which travels only retrogradely (**Figure 6a,b,c**). We observed that 78 and 81 hpi of PRV-Bartha gave expression in the spinocerebellar tract and neocortex, representing disynaptic transport. To isolate neocortical layer 5 neurons, whose axons comprise the descending corticopontine pathway, we analyzed neurons registered to deep layers, which comprised 64% of all contralaterally labeled neocortical neurons (**Figure S10d,e**).

**Figure 6.**
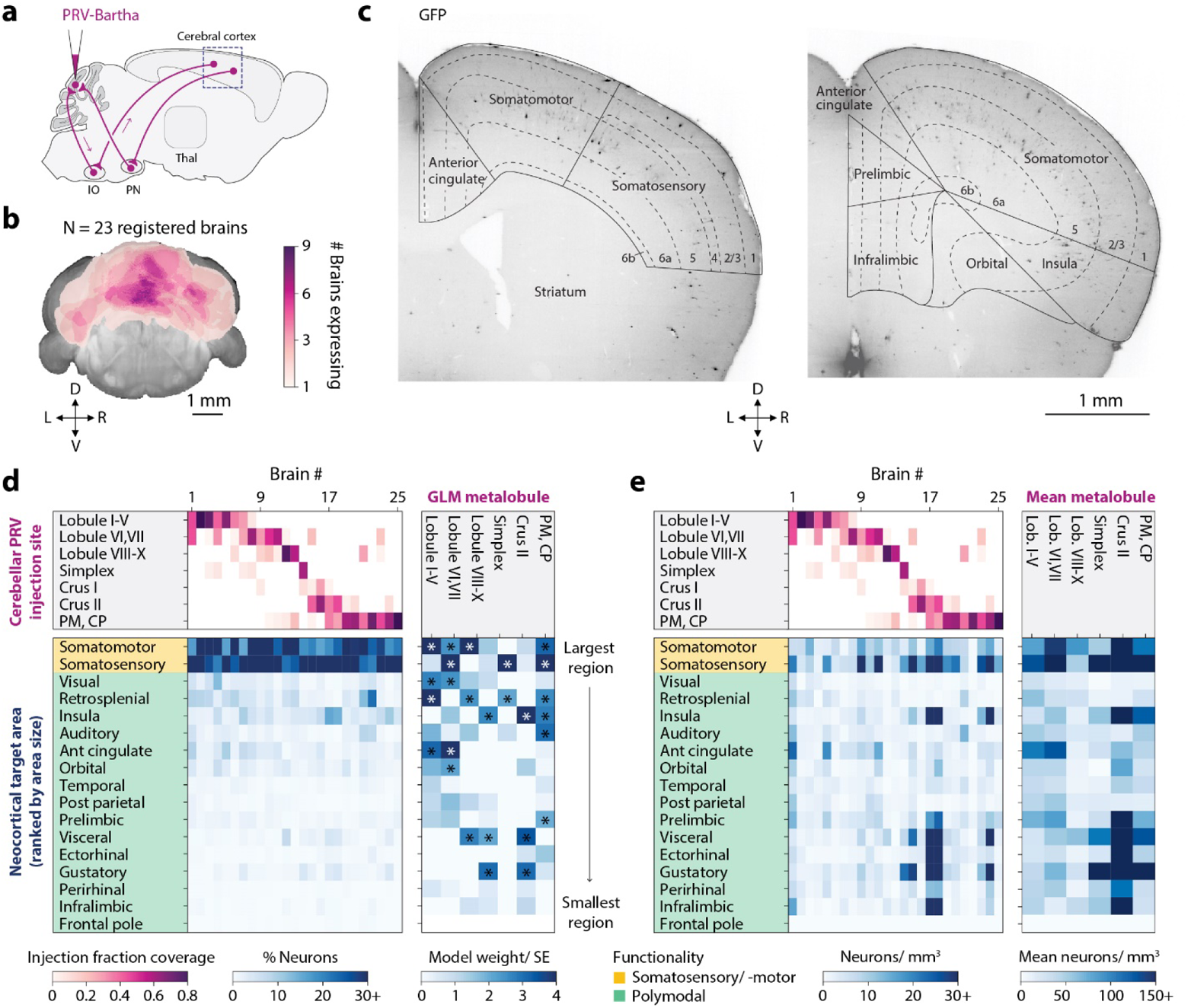
Descending projections to cerebellar cortex labeled using PRV-Bartha. (a) Retrograde disynaptic path from the cerebellar cortex to the neocortex traced using PRV-Bartha. (b) Coverage of cerebellum by neocortical timepoint injections marked by CTB-Alexafluor555. Projections show the number of injections covering each cerebellar location. (c) Maximum intensity projections (375 µm) with outlines defining neocortical structures (d) *Left*, fraction of neurons in each neocortical area across all injection sites. Injection coverage fractions (pink) and fraction of neurons (blue) for each injection. *Right*, generalized linear model showing the influence of each cerebellar region on neocortical expression. Heatmap (blue) shows the coefficient divided by standard error. Significant coefficients are marked with asterisks. (e) *Left*, density of neurons in neocortical areas across all injection sites. *Right*, mean neuron density in areas grouped by primary injection site.

Similar to the anterograde tracing results, we found the largest proportion of neurons in somatosensory and somatomotor areas (**Figure 6d and S7a,b**). Neuron densities were highest in somatosensory, somatomotor, and frontal cortex (**Figure 6e**). Two regions identified as sources of corticopontine axons by classical tracing (Wiesendanger and Wiesendanger, 1982) were labeled: anterior cingulate areas from injection of lobule VI and VII, and agranular insular cortex from crus II. In addition, retrosplenial and auditory areas were labeled from injections of paramedian lobule and copula pyramidis.

A GLM fit showed highest weighting in somatomotor, somatosensory, and frontal regions (**Figure 6d**). Weights in retrosplenial and visual cortex were smaller for vermal injections, and weights in gustatory, agranular insula, and visceral cortex were elevated for simplex and crus II injections. Averaging neuron density by primary injection site revealed that all cerebellar sites received dense projections from somatomotor and somatosensory cortex. Lobules I-VII and crus II received denser projections from anterior cingulate and prelimbic cortex compared to other injection sites. Crus II also received dense projections from infralimbic, agranular insula, gustatory, ectorhinal, and visceral cortex.

Descending corticopontine projections are largely ipsilateral (Wiesendanger and Wiesendanger, 1982) and pontocerebellar projections contralateral (Serapide et al., 2001). To test how well descending paths remain contralateral across multiple synaptic steps, we quantified the ratio of contralateral to ipsilateral cells for PRV-Bartha injections. Contralateral cells outnumbered ipsilateral cells in all major neocortical areas, with average contralateral-to-ipsilateral ratios of 1.4 in frontal cortex, 1.7 in posterior cortex, and 3.2 in somatomotor and somatosensory cortex. Contralateral:ipsilateral ratios were higher for hemispheric injection sites than vermal injection sites (**Table S3**).

Ascending DCN projections largely decussate to reach contralateral midbrain structures (Hashimoto et al., 2010). For H129 injections, we observed bilateral labeling at both di- and tri-synaptic timepoints. At the disynaptic timepoint, the mean contralateral:ipsilateral ratio was 2.5 in sensorimotor nuclei and 1.0 in polymodal association nuclei. Contralateral:ipsilateral ratios were highest for hemispheric injection sites (**Table S3**). Taken together, our H129 and Bartha observations suggest that the organization of projections between cerebellum and neocortex is, by total proportion, to sensorimotor cortical areas, most strongly contralateral in pathways that concern movement, and more symmetrically distributed for nonmotor paths.

Both Bartha and H129 tracing can identify patterns of cerebellar sites that project to, or receive information from, distinctive groups of neocortical sites. We performed multidimensional scaling on the pattern of fraction of neurons per neocortical region in PRV experiments. Similar neocortical expression patterns tended to have injections whose strongest contribution came from the same cerebellar lobule, confirming that animals with similar PRV neocortical labeling patterns had descending projections to similar cerebellar injection sites (**Figure S15a,b)**. Using the same analysis, H129 neocortical patterns showed a weaker relationship with cerebellar injection sites, with hotspots specific to each cluster (**Figure S15c,d)**.

### c-Fos mapping reveals brainwide patterns of activation consistent with transsynaptic tracing

The reciprocal paths we have identified suggest that cerebellum incorporates descending information and influences forebrain processing through multiple disynaptic paths. To test whether ascending paths could influence forebrain target activity, we measured expression of the immediate early gene c-Fos after optogenetic perturbation of PCs in stationary, head-fixed mice (**Figure 7**). c-Fos expression reflects cumulative neural activity, providing an independent means of observing long-distance influence (Dragunow and Faull, 1989). The hyperpolarizing pump ArchT was expressed in PCs using vermal AAV injections into L7-Cre+/− mice, using L7-Cre−/− mice as controls (**Figure 7a**). Inactivation of Purkinje cells, which inhibit neurons of the DCN, would be expected to have a net excitatory effect on thalamus and therefore neocortex.

**Figure 7.**
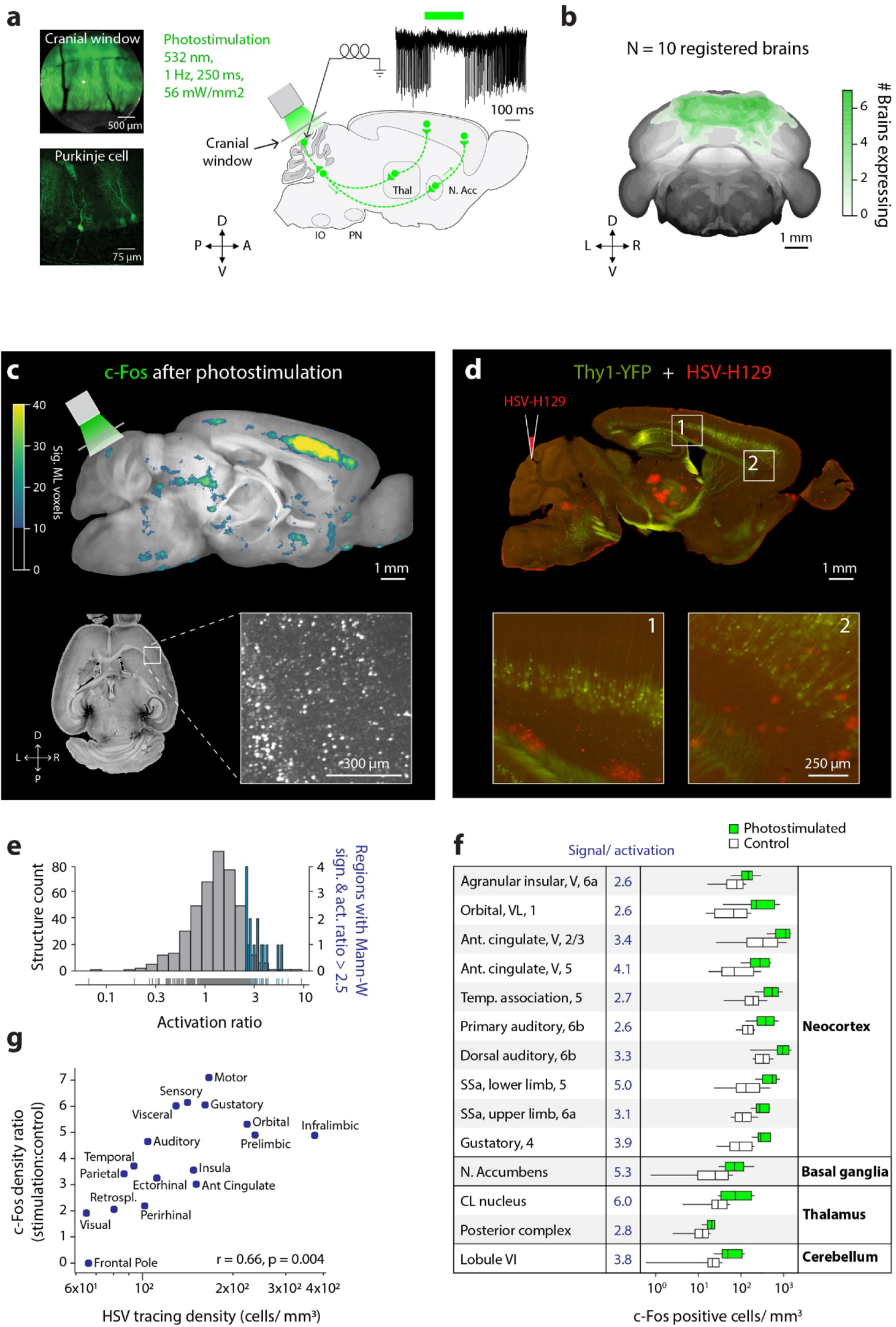
Cerebellar perturbation activates transsynaptically connected regions across the brain. (a) Experimental setup for photostimulating the inhibitory optogenetic protein ArchT-GFP through a cranial window over cerebellar lobule VI. *Top*, silencing of Purkinje cells as measured in brain-slice recordings after photostimulation. (b) Cerebellar ArchT-GFP expression. Coronal projections show the number of injections covering each location. (c) Neural activity identified by c-Fos immunostaining. *Top*, voxel-by-voxel regions of statistically significant c-Fos activation in Princeton Mouse Atlas (PMA) space (planes 320-360, 20 µm isotropic voxel size). *Bottom*, typical c-Fos labeling after perturbation. 132 µm maximum intensity projection. (d) Lobule VI transsynaptic targets labeled using H129 (red) injected into Thy1-YFP (green) mice. Standard non-clearing histological imaging, 50 µm section, 80 hpi. (e) Activation ratios, defined as c-Fos+ photostimulated mean divided by control-group mean. Regions were scored as responding (blue) if they had activation ratios greater than 2.5 and p<0.05 by two-tailed Mann-Whitney U test. (f) Distribution of c-Fos neurons for all responding regions. (g) c-Fos density ratio (mean stimulation density divided by mean control density) is positively correlated (Pearson’s r=0.66) with neocortical transsynaptic tracing density. For boxplots, center line showsmedian; box limits, upper and lower quartiles; whiskers, 1.5 times the interquartile range. Abbreviations: Ant, Anterior; C, caudal; D, dorsal; N, nucleus; Retrospl., retrosplenial area; SM, somatomotor areas; SS, somatosensory areas; V, ventral.

Photostimulation reduced PC firing during the light flash without perturbing arm speed (**Figure S11**). After 1 hour photostimulation, we used iDISCO+ to stain for c-Fos, and analyzed using ClearMap (Renier et al., 2016) (**Figure 7b,c**; **Figure S12a**) for comparison with H129 tracing (**Figure 7d**).

Fourteen structures showed both significant count differences by a Mann-Whitney U-test and an activation ratio (stimulation-group c-Fos average divided by control-group average) greater than 2.5 (**Figure 7e,f**). Strongest activation ratios occurred in the anterior cingulate cortex, CL, and the nucleus accumbens (**Figure 7f**). Lobule VI also showed elevated c-Fos counts, as expected for pulsed-light inactivation of PCs (Lee et al., 2015). As done in Renier et al. 2016, a voxel-wise t-test on cell count volumes (Clearmap stat.tTestVoxelization; **Figure S12b**) showed strong c-Fos expression in frontal neocortex, especially in deep and middle neocortical layers. These findings are consistent with transsynaptic tracing using both automated analysis of cleared tissue (**Figure S10d,e**) and standard tissue sectioning and epifluorescence microscopy (**Figure S10a-c**).

Among neocortical regions, mean c-Fos stimulation-to-control cell ratios and H129 densities were highly correlated (**Figure 7g**; c-Fos ratio vs. log HSV expression r=+0.66, p=0.004), indicating that brainwide patterns of neural activity coincide with patterns of ascending polysynaptic targets from lobule VI. C-Fos expression differed more from transsynaptic labeling under task conditions, indicating that the strength of influence also depends on underlying brain state (data not shown; J.L.V. et al., in preparation). Subcortical c-Fos measurements revealed further broad similarities with H129-VC22 labeling, including pontine nuclei, midbrain, superior colliculi, and hypothalamus (**Figure S12c-f**). Overall, these data show that c-Fos activation in awake animals coincides well with anatomical projection patterns identified by transsynaptic viral tracing.

## Discussion

We found that cerebellum projects to both motor and nonmotor thalamic and neocortical areas capable of supporting sensorimotor, flexible cognitive, and modulatory function. Homologous paths to these three systems often originated from a single injection site. Well-known sensorimotor regions contained the most connections by proportion, but nonmotor paths achieved comparable or higher local connection densities. Overall, these paths reached nearly all of neocortex after passing through a variety of thalamic, striatal, and midbrain intermediates.

### Sensory, motor, and associative pathways

In both thalamus and neocortex, the majority of neurons labeled by either anterograde or retrograde viruses were found in sensorimotor structures. In thalamus, these regions included known targets like ventral anterior-lateral (VA-L) and ventromedial (VM) thalamic nuclei (Angaut et al., 1985; Aumann and Horne, 1996; Aumann et al., 1994; Cicirata et al., 1990; Gornati et al., 2018; Teune et al., 2000). However, labeling was not restricted to classical thalamic motor regions: we also found widespread labeling in VPM, VPL, MGN, and dLGN, regions that were previously thought to receive only very limited cerebellar inputs. We also observed substantial nonmotor thalamic labeling, including the associative MD, LD and LP, suggesting routes by which cerebellar output is used for nonmotor functions.

In neocortex, by size most labeling was in somatomotor regions, but by density the strongest ascending projections went to anterior cingulate, prelimbic and infralimbic cortex, as well as agranular and orbital areas. Thus, our findings suggest that cerebellum simultaneously conveys output to functionally diverse targets.

### Modulatory pathways

The cerebellum is involved in sensory gating (Apps, 2000; Ozden et al., 2012). Our tracing suggests that such gating can contribute to a broader thalamic regulatory network. Lobules I-VII and crus II sent paths to TRN, a known, though neglected, monosynaptic target of DCN (Ando et al., 1995). Our dentate and interpositus connectivity with TRN agrees with rat (Cavdar et al., 2002; Chan-Palay, 2013) (**Figure S6**) and cat findings (Angaut et al., 1968; Nakamura, 2018). TRN is the only thalamic nucleus that is inhibitory and projects exclusively within thalamus. TRN may control sensory gain (Pinto et al., 2000) and information flow in and out of neocortex (Lam and Sherman, 2010). TRN also receives strong descending projections from neocortical layer 6 (Guillery et al., 1998; Lam and Sherman, 2010), a site of prominent expression in our work. This descending projection completes an inhibitory loop, which has been suggested to contribute to neocortical oscillations and synchrony (Bruno and Sakmann, 2006; Destexhe, 2000). Our findings add cerebellum as an input to this modulatory thalamocortical network.

### Transsynaptic labeling of cell bodies and summed paths

Our findings are largely consistent with monosynaptic tracing literature and provide new insights into the idea that cerebellum forms closed loops with motor, nonmotor, and modulatory domains in thalamus and neocortex. Transsynaptic viruses have two major advantages over traditional tracers (Fujita et al., 2020). First, they label postsynaptic somata rather than presynaptic terminals. Traditional tracers do not cross synapses, thus emphasizing large presynaptic axons up to hundreds of microns away from postsynaptic target neurons. This difference may account in part for the strength of our observed projection to TRN neurons, which have elaborate dendritic processes that extend into other thalamic nuclei (Pinault et al., 1997). Similarly, neocortical layer 5/6 pyramidal neuron dendrites extend to superficial layers where thalamocortical synapses occur. Presynaptic fiber tracing reveals connectivity to these superficial layers (Hooks et al., 2013) while our neocortical labeling is more consistent with electrophysiological recordings, which identify cortical recipient neurons in deeper layers (Llinás et al., 2002). Indeed, transsynaptic labeling can appear quite different from axonal tracing (compare H129 results in **Figure 2c** with AAV results in **Figure 3a-c**).

Second, transsynaptic tracers combine the summed contributions of multiple pathways with different intermediates. Our transsynaptic tracing results may also appear to differ from past monosynaptic cerebellothalamic tracing from DCN neurons, whereas we injected H129 one step previous, in the cerebellar cortex. Some of our connectivity to sensory thalamus may arise from non-DCN intermediates such as vestibular and parabrachial nuclei (Hashimoto et al., 2018; Shiroyama et al., 1999; Wijesinghe et al., 2015). The vestibular nuclei receive a well-known flocculonodular input, but we found even more total input from other cerebellar cortical regions (**Figure S4a**). Thus cerebellar influence over thalamus or neocortex may be transmitted by multiple paths. Vermal PCs appeared to make up most of the direct parabrachial connection (Hashimoto et al., 2018), consistent with our data (**Figure S14b**). However, apparent labeling in the parabrachial nucleus was likely to contain false positives because tissue clearing removed gray/white matter tissue differences between it and the superior cerebellar peduncle (a.k.a. brachium conjunctivum), which it wraps around and where labeling was considerable. We note that the MouseLight database contains no vermis-to-parabrachial PCs.

### Study limitations

Tracing viruses are powerful tools to identify neuroanatomical connections because they are self-amplifying and do not attenuate. However, the transsynaptic property of HSV (Nassi et al., 2015; Saleeba et al., 2019; Ugolini, 2010; Wojaczynski et al., 2015), brings different challenges from nonreplicating tracers.

Transsynaptic tracers carry uncertainty because the number of synapses traversed in a given time interval can vary. Uncertainty increases as the leading front of infected neurons advances. Our incubation times and analysis were designed to minimize these problems. Transport failures also contribute to uncertainty. To reduce nontransneural transport (Saleeba et al., 2019; Ugolini, 2010), injection size should be above but close to the infectivity threshold. For H129, the minimum dose is 10^4^ pfu (Ugolini et al., 1987), consistent with our injections. An injection close to threshold does not label all second- and third-order neurons, and can contribute to smaller subsets of DCNr and other intermediate neurons (see **Figure S3a,b**), as well as thalamic and neocortical neurons. We addressed this problem by combining data from multiple injections. Our injections spanned at least two microzones or zebrin (aldolase C) bands. Future experiments can achieve microzone specificity using conditional H129 strains with zebrin- or non-zebrin-dependent recombinase expression.

H129 is not taken up by fibers of passage, but can label tracts by infecting oligodendrocytes; this does not affect mapping applications since no further transmission occurs (Wojaczynski et al., 2015). For incubations of 96 hours or longer, H129 may follow retrograde paths (Wojaczynski et al., 2015). We used the shortest necessary incubation times, often considerably shorter than other work (**Table S1**). Control experiments using shorter timepoints (28-36 hpi) and brainstem analysis at disynaptic timepoints indicated H129 retrograde transport was minimal. Because PRV and HSV transport speed varies in different nervous system structures (Card et al., 1997; Ugolini, 1992) and spread characteristics vary across strains (Garner and LaVail, 1999), future studies will need similar control experiments to guide interpretation.

Finally, tissue-clearing methods remove gray-white matter tissue differences, making it necessary to rely on coordinates and anatomical registration. In such circumstances, registration-based counts should be supplemented by traditional tissue preparation methods (see *Methods, Automated Detection of Virally Labeled Cells*).

### Cerebello-thalamo-cortical loops

Cerebello-neocortical connectivity has been suggested to be organized into closed loops, in which each neocortical region has only a few principal cerebellar partners, and vice versa (Buckner et al., 2011; Strick et al., 2009). In this scenario, microzones, which are anatomically repeated throughout the cerebellum, would process incoming neocortical information in a similar manner and then return output to the originating neocortical structures. Our work broadens this framework and suggests that a region of cerebellar cortex, spanning several microzones, sends projections to a functionally diverse set of forebrain targets.

A cerebellar-neocortical map with multiple targets is suggested by several known stages of anatomical divergence. Notably, DCN locations project to multiple thalamic areas and each thalamic area typically projects to multiple neocortical areas (Aumann et al., 1998; Jones, 2012). The tracing patterns we observed indicate that these steps have more divergence than suggested by a strict closed-loop view. Our work is consistent with the idea (Aoki et al., 2017; Buckner et al., 2011) that the classically-proposed principal partners comprise only a small part of the total anatomical relationship, and suggests that cerebellum can simultaneously provide similar output to a functionally diverse set of structures.

The descending pathways are most strongly concentrated in somatomotor and somatosensory cortex, consistent with recent anterograde tracing from neocortex to mossy fibers in the mouse brain (Henschke and Pakan, 2020) (though also note paths from frontal areas to crus II; (Suzuki et al., 2012). Our observed connectivity patterns suggest cerebellum incorporates information from somatomotor, somatosensory, and other cortex to exert returning influence on a wide distribution of thalamic and neocortical targets. Our results provide an anatomical framework to be used by future studies to probe function and circuit mechanisms.

### Nonmotor functions of lobule VI and crus I

Nonmotor functions have been suggested for lobule VI in the posterior vermis, and crus I in the posterior parts of the hemispheres (Badura et al., 2018; Stoodley et al., 2017). We found that lobule VI sends strong projections to mediodorsal and paraventricular thalamic nuclei and to frontal neocortical regions, which serve a range of cognitive and affective functions (Marton et al., 2018; Yamamuro et al., 2020). Because postnatal refinement of neural circuitry is activity-dependent (Hubel and Wiesel, 1965), this projection may explain why lobule VI perturbations affect cognitive and social development in rodents (Badura et al., 2018) and humans (Wang et al., 2014) and the association of posterior vermal abnormalities with a high risk of ASD (Courchesne et al., 1988; Limperopoulos et al., 2007). Also, both H129 anterograde tracing (**Figure S10a-c**) and c-Fos mapping revealed an association between lobule VI and nucleus accumbens (NAc), the main component of the ventral striatum, which is implicated in reward learning and motivation (Salgado and Kaplitt, 2015).

Rodent crus I is thought to be homologous to primate crus I/II (Sugihara, 2018), which has more prefrontal and parietal connections (Balsters et al., 2014). Rodent crus II is likely to be related to human HVIIB (Luo et al., 2017), a structure with more motor connections (Balsters et al., 2014). Disruption of rodent crus I activity in adulthood or juvenile life leads to deficits in adult flexible behavior (Badura et al., 2018; Stoodley et al., 2017) and adult disruption shortens the time constant of a working memory task (Deverett et al., 2018). We found that crus I projects to lateral dorsal and paraventricular nuclei and frontal neocortical regions. Crus I also projects to paraventricular nucleus (Yamamuro et al., 2020) and septal and amygdalar regions (Heath et al., 1978), providing possible substrates for the observation that juvenile disruption of crus I leads to long-lasting deficits in social preference (Badura et al., 2018).

### BrainPipe: a pipeline for long-distance transsynaptic mapping

Many individual cerebellum-to-forebrain connections have been previously reported. Our work now presents a brainwide survey using transsynaptic tracing combined with whole-brain quantification. By comparing quantified projection targets at different injection locations of the same viral tracer, our study allows for relative comparison of output targets. Transsynaptic tracing studies have relied on time-consuming human identification for analysis (Kelly and Strick, 2003; McGovern et al., 2012; Song et al., 2009; Wojaczynski et al., 2015). BrainPipe, which combines transsynaptic tracing, whole-brain clearing and microscopy, automated neuron counting, and atlas registration, is efficient, accurate, and scalable to the whole brain. Adapting BrainPipe to other studies requires only a different annotated dataset to train a new convolutional neural network to identify objects of interest. BrainPipe is scalable to larger datasets as microscopy resolution improves, since it can run on high-performance computing clusters. Our light-sheet brain atlas overcame the problem of registration across imaging modalities by first creating a modality-specific template brain. Finally, our software (github.com/PrincetonUniversity/pytlas) is capable of generating atlases for other imaging modalities.

We took two approaches to quantifying relative projection strength: (1) subregion cell count fraction (*e.g.* VPM or infralimbic cortex) relative to parent structure (*e.g.* whole thalamus or neocortex) or (2) cell count density of a structure. Fraction of total expression allows for relative projection strength comparison within a parent structure, conveying information about the distribution of total influence. In contrast, density takes local structure into account and provides information about the concentrated influence on recipient targets. For example, although the great majority of projections to neocortex were found in somatomotor and somatosensory cortices, smaller prefrontal areas receive a higher density of projections. This suggests cerebellum’s capacity for influence on prefrontal areas, while smaller in total terms, might still be focused enough to substantially affect nonmotor behavior (Badura et al., 2018; Stoodley et al., 2017).

### From local cerebellar circuitry to global brain function

Local cerebellar circuitry is thought to make rapid predictions about future states, which then modulate the activity of other brain regions (Solari and Stoner, 2011). Contextual information comes from the mossy fiber-granule cell pathway, and teaching signals come from climbing fibers to drive learning processes (Lisberger, 2020) DCN and vestibular nuclei have their own processing and learning rules and serve as portals to other brain regions (Lisberger, 2020; Ruigrok et al., 2015). The same processing principles may be shared across homologous across cerebellar regions (Kebschull et al., 2020), with functional role determined by the particular long-distance partners. Each part of the cerebellar cortex manages a massive convergence of diverse incoming information from a distinct assortment of distant brain regions (Léna and Popa, 2016). The cerebellar cortex may thus generate predictions to fine-tune activity across motor and nonmotor functions (Deverett et al., 2018; Schmahmann and Sherman, 1998; Wang et al., 2014) as its output re-converges onto cerebellar and vestibular nuclei. Our tracing identifies the anatomical potential for output paths from one cerebellar site to affect multiple targets at once.

## STAR Methods

### Resource availability

#### LEAD CONTACT

Further information and requests for resources and reagents should be directed to and will be fulfilled by the Lead Contact: Samuel S.-H. Wang.

#### MATERIALS AVAILABILITY

All unique/stable reagents generated in this study are available from the Lead Contact without restriction.

#### DATA AND CODE AVAILABILITY

The Princeton Mouse Atlas data have been deposited at https://brainmaps.princeton.edu/2020/09/princeton-mouse-brain-atlas-links/ and are publicly available as of the date of publication. Aligned viral tracing injection data have been deposited at https://brainmaps.princeton.edu/2021/05/pisano_viral_tracing_injections/ and are publicly available as of the date of publication. Unprocessed data reported in this paper will be shared by the lead contact upon reasonable request.

All original code has been deposited at Zenodo and is publicly available as of the date of publication. DOIs are listed in the key resources table.

Any additional information required to reanalyze the data reported in this paper is available from the lead contact upon request.

### Experimental model and subject details

Experimental procedures were approved by the Institutional Animal Care and Use Committees of Princeton University (protocol number 1943-19), the University of Idaho (protocol number 2017-66), and the Dutch national experimental animal committees (DEC), and performed in accordance with the animal welfare guidelines of the National Institutes of Health (USA) or the European Communities Council Directive (Netherlands).

#### ORGANISM

Adult mice (C57BL/6J, 8-12 weeks old, The Jackson Laboratory, 000664) of both sexes were used for transsynaptic tracing viral studies. For classic sectioning-based histology transsynaptic viral injections two adult male Thy1-YFP (aged 22 weeks, B6.Cg-Tg (Thy1-YFP)HJrs/J) mice were used. For DCN and TRN AAV injections adult male mice (C57BL/6J, 1-3 month) were used. For c-Fos mapping experiments L7-Cre +/− and −/− males were used (B6; 129-Tg (Pcp2-cre)2Mpin/J, 004146, 56 days or older, bred in-house). Controls (−/−) and experimental (+/−) were littermates and housed together from birth. For electrophysiological confirmation of ArchT expression three 10 week-old male (B6.Cg-Tg (Pcp2-cre)3555Jdhu/J, 010536, The Jackson Laboratory) mice were used. All mice were group housed in shared cages with a maximum of 5 mice per cage. Mice were provided nesting and housing material for enrichment.

#### CELL LINE

The African green monkey kidney epithelial cell line Vero (ATCC cell line CCL-81, https://web.expasy.org/cellosaurus/CVCL_0059) was used to propagate and titer HSV-H129-VC22. Cells were grown at 37°C with 5% CO_2_ in DMEM supplemented with 10% FBS and 1% penicillin/streptomycin. Cells were obtained from ATCC but were not authenticated. The sex of the cell line is female.

### Methods details

#### OVERVIEW OF AUTOMATED PIPELINE FOR TRANSSYNAPTIC TRACING

In order to identify and quantify cerebellar connectivity on a long-distance scale, we developed a pipeline, BrainPipe, to enable automated detection of transsynaptically labeled neurons using the anterogradely-transported HSV-H129 (Wojaczynski et al., 2015), identifying cerebellar output targets, and retrogradely-transported PRV-Bartha (Smith et al., 2000), identifying the descending corticopontine pathway, comprised mostly of layer 5 pyramidal neurons (Legg et al., 1989). Mouse brains with cerebellar cortical injections of Bartha or H129 were cleared using iDISCO+. We then imaged the brains using light-sheet microscopy, generating brain volumes with a custom Python package. Next, to ensure accurate anatomical identification across brains, we created a local light-sheet template, the Princeton Mouse Brain Atlas (PMA) and quantified registration performance of individual volumes to the local template. We then determined the transform between the PMA and the Allen Brain Atlas, enabling standardization of our results with the current field standard. Next, to automatically and accurately detect labeled cells, we developed a convolutional neural network whose performance approached that of human classifiers.

#### ANIMAL EXPERIMENTATION

Experimental procedures were approved by the Institutional Animal Care and Use Committees of Princeton University (protocol number 1943-19), the University of Idaho (protocol number 2017-66), and the Dutch national experimental animal committees (DEC), and performed in accordance with the animal welfare guidelines of the National Institutes of Health (USA) or the European Communities Council Directive (Netherlands).

#### VIRUS SOURCES

HSV-1 strain H129 recombinant VC22 (H129-VC22) expresses EGFP-NLS, driven by the CMV immediate-early promoter and terminated with the SV40 polyA sequence. To engineer this recombinant, we used the procedure previously described to construct HSV-772, which corresponds to H129 with CMV-EGFP-SV40pA (Wojaczynski et al., 2015). We generated plasmid VC22 by inserting into plasmid HSV-772 three tandem copies of the sequence for the c-Myc nuclear localization signal (NLS) PAAKRVKLD (Ray et al., 2015), fused to the carboxy-terminus of EGFP. Plasmid VC22 contains two flanking sequences, one of 1888-bp homologous to HSV-1 UL26/26.5, and one of 2078-bp homologous to HSV-1 UL27, to allow insertion in the region between these genes. HSV-1 H129 nucleocapsid DNA was cotransfected with linearized plasmid VC22 using Lipofectamine 2000 over Vero cells, following the manufacturer’s protocol (Invitrogen). Viral plaques expressing EGFP-NLS were visualized and selected under an epifluorescence microscope. PRV-Bartha-152 (Smith et al., 2000), which drives the expression of GFP driven by the CMV immediate-early promoter and terminated with the SV40 polyA sequence, was a gift of the laboratory of Lynn W. Enquist. Adeno-associated virus was obtained from Addgene (https://www.addgene.org).

#### IN VIVO VIRUS INJECTIONS

##### Surgery for HSV and PRV injections

Mice were injected intraperitoneally with 15% mannitol in 0.9% saline (M4125, Sigma-Aldrich, St. Louis, MO) approximately 30 minutes before surgery to decrease surgical bleeding and facilitate viral uptake. Mice were then anesthetized with isoflurane (5% induction, 1-2% isoflurane/oxygen maintenance vol/vol), eyes covered with ophthalmic ointment (Puralube, Pharmaderm Florham Park, NJ), and stereotactically stabilized (Kopf Model 1900, David Kopf Instruments, Tujunga, CA). After shaving hair over the scalp, a midline incision was made to expose the posterior skull. Posterior neck muscles attaching to the skull were removed, and the brain was exposed by making a craniotomy using a 0.5 mm micro-drill burr (Fine Science Tools, Foster City, CA). External cerebellar vasculature was used to identify cerebellar lobule boundaries to determine nominal anatomical locations for injection. Injection pipettes were pulled from soda lime glass (71900-10 Kimble, Vineland, NJ) on a P-97 puller (Sutter Instruments, Novato, CA), beveled to 30 degrees with an approximate 10 μm tip width, and backfilled with injection solution.

##### AAV deep cerebellar nuclear injections

During stereotaxic surgery, mice (n=4) were anesthetized with isoflurane (PCH, induction: 5%; maintenance: 2.0-2.5%) and received a mannitol injection intraperitoneally (2.33 g/kg in Milli-Q deionized water) and a rimadyl injection subcutaneously (5 mg/kg carprofen 50 mg/ml, Pfizer, Eurovet, in 0.9% NaCl solution). Body temperature was kept constant at 37°C with a feedback measurement system (DC Temperature Control System, FHC, Bowdoin, ME, VS). Mice were placed into a stereotactic frame (Stoelting, Chicago laboratory supply), fixing the head with stub ear bars and a tooth bar. Duratears eye ointment (Alcon) was used to prevent corneal dehydration. A 2 cm sagittal scalp incision was made, after which the exposed skull was cleaned with sterile saline. Mice were given 2 small (diameter ±1 mm) craniotomies in the interparietal bone (−2 mm posterior relative to lambda; 1.8 mm lateral from midline) for virus injection. Craniotomies were performed using a hand drill (Marathon N7 Dental Micro Motor). A bilateral injection of AAV5-Syn-ChR2-eYFP (125 nl of titer 7×10¹² vg/ml per hemisphere, infusion speed ~0.05 µl/minute) in the AIN was done using a glass micropipette controlled by a syringe. This AAV was used because it gave reliable strong axon terminal labeling and because the animals were also used for another optogenetic study. After slowly lowering the micropipette to the target site (2.2 mm ventral), the micropipette remained stationary for 5 minutes before the start of the injection, and again after finishing the injection. The micropipette was then withdrawn slowly from the brain at a rate of ~1 mm/minute. Craniotomies and skin were closed and mice received post-op rimadyl. Animals were perfused transcardially 3 weeks after viral injection using 4% paraformaldehyde. Brains were collected postmortem, co-stained for DAPI (0100-20, Southern Biotech, Birmingham, AL), coronally sectioned at 40 µm/slice, and imaged with an epifluorescence microscope at 20x (Nanozoomer, Hamamatsu, Shizuoka, Japan).

To visualize YFP labeled fibers and vGluT2-positive terminals in the thalamus, 40 micron thick slices were stained for vGluT2 using anti-guinea pig Cy5 as the primary (Millipore Bioscience Research reagent 1:2000 diluted in PBS containing 2% NHS and 0.4% Triton) and anti-GP Cy5 (1:200; Jackson Immunoresearch) as the secondary antibody. Images were taken using a confocal LSM 700 microscope (Carl Zeiss). Terminals positive to VGluT2 staining were identified and morphologically studied using confocal images that were captured using excitation wavelengths of 488 nm (YFP) and 639 nm (Cy5). High-resolution image stacks were acquired using a 63X 1.4 NA oil objective with 1X digital zoom, a pinhole of 1 Airy unit and significant oversampling for deconvolution (voxel dimension is: 46 nm width × 46 nm length × 130 nm depth calculated according to Nyquist factor; 8 bits per channel; image plane 2048 × 2048 pixels). Signal-to-noise ratio was improved by 2 times line averaging.

##### AAV TRN injections

During stereotaxic surgery, mice (1-3 months of age) were anesthetized with isoflurane (VetOne, induction: 3-5%; maintenance: 1.5-2.5%). For analgesic support mice provided oral carprofen *ad libitum* from the day before and through 24 h after surgery and given slow release meloxicam (4 mg/kg; ZooPharm, Larami, WY). Body temperature was maintained by a warming blanket (Stoelting, Wood Dale, IL) under the animal throughout the surgery. Mice were placed into a stereotactic frame (Kopf, Tujunga, CA), fixing the head with non-rupture ear bars, a tooth bar and nose cone. Puralube Vet eye ointment (Dechra) was used to prevent corneal dehydration. A sagittal scalp incision was made, after which the exposed skull was cleaned with sterile saline. A single small craniotomy (diameter 0.6 mm) was made in the parietal bone (−1.3 mm posterior relative to lambda; 2.3 mm right of midline) for virus injection. Craniotomies were performed using a stereotaxic-mounted drill (Foredom K.1070 micromotor drill). A unilateral 200-300 nl injection of AAVrg-hSyn-Chronos-GFP (9.0×10^12^; Addgene, Watertown, MA) at an infusion speed of 0.01 μl/minute in the right TRN was done using a glass syringe and needle (Hamilton Company, Franklin, MA). After slowly lowering the needle to the target site (−2.9 mm ventral), the needle remained stationary for 1 minute before the start of the injection, and for 5 min after finishing the injection. The needle was then withdrawn slowly from the brain. Craniotomies and skin were closed using removable staples and mice continued to receive oral carprofen *ad libitum* for 24hrs post-surgery. Animals were euthanized 20-25 days after viral injection and brains were fixed in 4% paraformaldehyde. Brains were collected, frozen, and coronally sectioned into 40 μm slices, then co-stained with Hoechst 33324 (5 µg/ml; Invitrogen), chicken anti-GFP (1:500; Novus Biologicals; NB100-1614), and rabbit anti-parvalbumin (1:500; ZRB1218; Millipore Sigma), and imaged with a confocal fluorescence microscope at 10X and 20x (Nikon Instruments TiE inverted microscope with Yokogawa X1 spinning disk) or an epifluorescence microscope at 2.5X and 10X (Zeiss Axio Imager.M2).

##### Transsynaptic viral tracing for tissue clearing (HSV-H129 and PRV-Bartha)

Transsynaptic viral tracing studies used male and female 8-12 week-old C57BL/6J mice (The Jackson Laboratory, Bar Harbor, Maine). Injection solution was prepared by making a 9:1 dilution of either H129 or PRV virus stock to 0.5% cholera toxin B conjugated to Alexa Fluor 555 in saline (CTB-555, C22843, Sigma-Aldrich; as per Ref. (Conte et al., 2009). At the timepoints used CTB-555 persisted at the injection site. Pipettes were inserted perpendicular to tissue surface to a depth of approximately 200 µm. **Table S3** describes injection parameters and viral stock concentrations for each type of H129 and PRV experiment.

Pressure injections delivered 80 to 240 nl into the target location. Consistent with prior literature we observed that minimum injections of 10^4^ PFUs were required for successful HSV-H129 infection (Ugolini et al., 1987). Smaller injections consistently produced unsuccessful primary infections and thus no transsynaptic spread. Unfortunately, this feature also prevented consistent injections of single zones as defined by zebrin staining.

After H129 or PRV viral injection, Rimadyl (0.2 ml, 50 mg/ml, carprofen, Zoetis, Florham Park, NJ) was delivered subcutaneously. At the end of the post-injection incubation period, animals were overdosed by intraperitoneal injection of ketamine/xylazine (ketamine: 400 mg/kg, Zetamine, Vet One, ANADA #200-055; xylazine: 50 mg/kg, AnaSed Injection Xylazine, Akorn, NADA #139-236) and transcardially perfused with 10 ml of 0.1 M phosphate-buffered saline (PBS) followed by 25 ml 10% formalin (Fisher Scientific 23-245685). Tissue was fixed overnight in 10% formalin before the iDISCO+ clearing protocol began.

##### Transsynaptic timepoint determination

To determine the optimal timepoints for primarily disynaptic (*i.e.* Purkinje cell to cerebellar/vestibular nuclei to thalamus) and primarily trisynaptic (additionally to neocortex) anterograde targets, we injected H129 into the cerebellar cortex of mice and examined tissue between 12 and 89 hpi (**Figure 1b,c**; (30, 36, 41, 49, 54, 58, 67, 73, 80, 82 and 89 hours post-injection of midline lobule VI). At 54 hpi, thalamic labeling was observed with little neocortical labeling (**Figure 1c,d**), so we used this as the disynaptic timepoint (**Table S1**). Labeling was seen in other midbrain and hindbrain areas, consistent with known monosynaptic anterograde targets of the cerebellar and vestibular nuclei (Teune et al., 2000; Wijesinghe et al., 2015). Neocortical labeling was weak at 73 hpi and spanned its extent by 82 hpi; we therefore defined 80 hpi as our trisynaptic timepoint.

For retrograde transport experiments, incubation times for PRV-Bartha injections were determined by immunostaining for GFP (48, 60, 72, 78, 81, 84 and 91 hpi of midline lobule VI) targeting the canonical descending pathway: neocortex to brainstem to cerebellar cortex. We selected timepoints with the goal of achieving sufficient labeling for detection, while minimizing incubation periods, given that with increasing long distance, transport time is increasingly dominated by axon-associated transport mechanisms (Callaway, 2008; Card et al., 1999; Granstedt et al., 2013; Miranda-Saksena et al., 2018), leading to labeling of alternative paths and retrograde paths after 96 hpi (Wojaczynski et al., 2015). Our selected timepoints were shorter than published timepoints (**Table S1**), and were therefore likely to reduce the degree of supernumerary synaptic spread.

#### VIRAL TRACING WITH TISSUE SECTIONING AND SLIDE-BASED MICROSCOPY

##### Viral tracing with classical sectioning-based histology: HSV-772 cerebellar injections

Adult Thy1-YFP male mice (YFP +, n=2, B6.Cg-Tg (Thy1-YFP)HJrs/J, 003782, The Jackson Laboratory, 22 weeks), were prepared for surgery, in a similar fashion as in *Transsynaptic viral tracing for tissue clearing (H129 and Bartha)*. We used the HSV recombinant HSV-772 (CMV-EGFP, 9.02 × 10^8^ PFU/ml) (Wojaczynski et al., 2015), an H129 recombinant that produces a diffusible EGFP reporter. Again, using a 9:1 HSV:CTB-555 injection solution, 350 nl/injection was pressure-injected into two mediolateral spots at midline lobule VIa. Eighty hours post-injection, animals were overdosed using a ketamine/xylazine mixture as described previously. Brains were extracted and fixed overnight in 10% formalin and cut at 50 µm thickness in PBS using a vibratome (VT1000S, Leica). Sections were immunohistochemically blocked by incubating for 1 hour in 10% goat serum (G6767-100ML, Sigma-Aldrich, St. Louis, MO), 0.5% Triton X100 (T8787-50ML, Sigma-Aldrich) in PBS. Next sections were put in primary antibody solution (1:750 Dako Anti-HSV in 2% goat serum, 0.4% Triton X100 in PBS) for 72 hours at 4°C in the dark. Sections were washed in PBS 4 times for 10 minutes each, and then incubated with secondary antibody (1:300 goat anti-rabbit-AF647 in 2% goat serum, 0.4% Triton X100 in PBS) for two hours. Another series of PBS washes (four times, 10 minutes each) was done before mounting onto glass microscope slides with Vectashield mounting agent (H-1000, Vector Laboratories, Burlingame, CA). Sections were fluorescently imaged at 20x (Nanozoomer, Hamamatsu, Shizuoka, Japan) and at 63x with 5 μm z steps (Leica SP8 confocal laserscanning microscope).

#### TISSUE CLEARING AND LIGHT-SHEET MICROSCOPY

##### iDISCO+ tissue clearing

After extraction, brains were immersed overnight in 10% formalin. An iDISCO+ tissue clearing protocol (Renier et al., 2016) was used (*Supplemental clearing worksheet*). Brains were dehydrated step-wise in increasing concentrations of methanol (Carolina Biological Supply, 874195; 20, 40, 60, 80, 100% in doubly distilled water (ddH_2_O), 1 hr each), bleached in 5% hydrogen peroxide/methanol solution (Sigma, H1009-100ML) overnight, and serially rehydrated (methanol: ddH_2_O 100, 80, 60, 40, 20%, 1 hr each). Brains were washed in 0.2% Triton X-100 (Sigma, T8787-50ML) in PBS, then in 20% DMSO (Fisher Scientific D128-1) + 0.3 M glycine (Sigma 410225-50G) + 0.2% Triton X-100/PBS at 37°C for 2 days. Brains were then immersed in a blocking solution of 10% DMSO + 6% donkey serum (EMD Millipore S30-100ml) + 0.2% Triton X-100 + PBS at 37°C for 2-3 days to reduce non-specific antibody binding. Brains were then twice washed for 1 hr/wash in PTwH: a solution of PBS + 0.2% Tween-20 (Sigma P9416-50ML) + 10 µg/ml heparin (Sigma H3149-100KU).

For H129 and c-Fos antibody labeling, brains were incubated with primary antibody solution (see **Table S3** for antibody concentrations) consisting of 5% DMSO + 3% donkey serum + PTwH at 37°C for 7 days. Brains were then washed in PTwH at least 5 times (wash intervals: 10 min, 15, 30, 1 hr, 2 hr), immunostained with secondary antibody in 3% donkey serum/PTwH at 37°C for 7 days, and washed again in PTwH at least 5 times (wash intervals: 10 min, 15, 30, 1 hr, 2 hr). Finally, brains were serially dehydrated (methanol: ddH_2_O: 100, 80, 60, 40, 20%, 1 hr each), treated with 2:1 dichloromethane (DCM; Sigma, 270997-2L):methanol and then 100% DCM, and placed in the refractive index-matching solution dibenzyl ether (DBE; Sigma, 108014-1KG) for storage at room temperature before imaging.

##### Light-sheet microscopy for transsynaptic tracing

Cleared brain samples were glued (Loctite, 234796) ventral side down on a custom-designed 3D-printed holder and imaged in DBE using a light-sheet microscope (Ultramicroscope II, LaVision Biotec., Bielefeld, Germany). Version 5.1.347 of the ImSpector Microscope controller software was used. An autofluorescent channel for registration purposes was acquired using 488 nm excitation and 525 nm emission (FF01-525/39-25, Semrock, Rochester, New York). Injection sites, identified by CTB-555, were acquired at 561 nm excitation and 609 nm emission (FF01-609/54-25, Semrock). Cellular imaging of virally infected cells (anti-HSV Dako B011402-2) was acquired using 640 nm excitation and 680 nm emission (FF01-680/42-25, Semrock). Cellular-resolution imaging was done at 1.63 µm/pixel (1x magnification, 4x objective, 0.28 NA, 5.6-6.0 mm working distance, 3.5 mm × 4.1 mm field of view, LVMI-FLuor 4x, LaVision Biotech) with 3×3 tiling (with typically 10% overlap) per horizontal plane. Separate left- and right-sided illumination images were taken every 7.5 micrometers step size using an excitation-sheet with a numerical aperture of 0.008. A computational stitching approach (Bria and Iannello, 2012) was performed independently for left- and right-side illuminated volumes, followed by midline sigmoidal-blending of the two volumes to reduce movement and image artifacts.

The originally-reported iDISCO+ clearing methodology showed no decline in number of c-Fos+ cells detected as a function of imaging depth. To empirically confirm this in the deepest brain structure imaged, the thalamus, we quantified cell counts at the disynaptic timepoint as a function of location in all three axes (**Figure S13d**). Cell counts were not strongly correlated with position in any axis.

#### REGISTRATION AND ATLAS PREPARATION

##### Image registration

Most registration software cannot compute transformation with full-sized light-sheet volumes in the 100-200 gigabyte range due to computational limits. Using mid-range computers, reasonable processing times are obtained with file sizes of 300-750 megabytes, which for mouse brain corresponds to 20 µm/voxel. Empirically, we found that light-sheet brain volumes to be aligned (“moving”) resampled to approximately 140% the size of the reference (“fixed”) atlas volume yielded the best registration performance. Alignment was done by applying an affine transformation to generally align with the atlas, followed by b-spline transformation to account for brain-subregion variability among individual brains.

For uniformity among samples, registration was done using the autofluorescence channel, which has substantial signal at shorter wavelengths useful for registration (Renier et al., 2014). In addition to autofluorescence-to-atlas registration, the signal channel was registered using an affine transformation to the autofluorescence channel to control for minor brain movement during acquisition, wavelength-dependent aberrations, and differences in imaging parameters (Renier et al., 2016).

Affine and b-spline transformations were computed using elastix (Klein et al., 2010; Shamonin et al., 2013); see supplemental Elastix affine and b-spline parameters used for light-sheet volume registration. Briefly, the elastix affine transform allows for translation (t), rotation (R), shearing (G), and scaling (S) and is defined as:

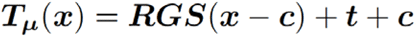

where c is a center of rotation and t is a translation. The elastix b-spline transformation allows for nonlinearities and is defined as:

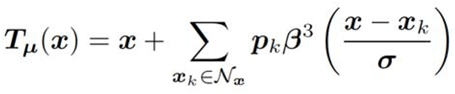

Where *x* are control points, β^3^ (*x*) the B-spline polynomial, p the b-spline coefficient vectors, N*x*, B-spline compact support control points, and Cl is the b-spline compact control point-spacing (see Ref. (Klein and Staring, 2015), pages 8-10 for reference). For the assignment of cell centers to anatomical locations, we calculated transformations from cell signal space to autofluorescence space (affine only) and autofluorescence space to atlas space (affine and b-spline; **Figure S13a**).

##### Princeton Mouse Atlas generation

To generate a light-sheet atlas with a complete posterior cerebellum, autofluorescent light-sheet volumes from 110 mice (curated to eliminate distortions related to damage, clearing, or imaging) were resampled to an isotropic 20 µm per voxel resolution (**Figure 1g-j** and **Figure S2a**). We selected a single brain volume to use as the fixed (template) volume for registration of the other 109 brains and computed the transformations between the other 109 brains and the template brain. The registration task was parallelized from ClearMap (Renier et al., 2016) adapting code for use on a Slurm-based (Yoo et al., 2003) computing cluster.

After registration, all brains were pooled into a four-dimensional volume (brain, x, y, z), and the median voxel value at each xyz location was used to generate a single median three-dimensional volume. Flocculi and paraflocculi, which can become damaged or deformed during extraction and clearing, were imaged separately from a subset of 26 brains in which these structures were intact and undeformed. Manual voxel curation sharpened brain-edges in areas where pixel intensity gradually faded. Finally, contrast-limited adaptive histogram equalization (skimage.exposure.equalize_adapthist) applied to the resulting volume increased local contrast within brain structures, generating the final PMA (**Figure S2b** and **S13b**). We then determined the transformation between the PMA and the Allen Brain CCFv3 (Allen Institute for BrainScience, 2012) space in order to maintain translatability. Our software for basic atlas creation with an accompanying Jupyter tutorial notebook is available online via github.com/PrincetonUniversity/pytlas. Volumetric projection renderings were made using ImageJ (Schmid et al., 2010); 3D project function (**Figure S2a**). The PMA interactive three-dimensional rendering of the PMA is available http://brainmaps.princeton.edu/pma_neuroglancer and can be downloaded from https://brainmaps.princeton.edu/pma_landing_page.

##### Generation of 3D printable files

To generate 3D-printable Princeton Mouse Atlas files usable for experimental and educational purposes, we loaded volumetric tiff files as surface objects using the ImageJ-based 3D viewer. After downsampling by a factor of 2 and intensity thresholding, data were then imported to Blender (Roosendaal and Selleri, 2004), where surfaces were smoothed (Smooth Vertex tool) before finally exporting as stereolithography (stl) files.

#### AUTOMATED DETECTION OF VIRALLY LABELED CELLS

##### BrainPipe, an automated transsynaptic tracing and labeling analysis pipeline

Whole-brain light-sheet volumes were analyzed using our pipeline, BrainPipe. BrainPipe consists of three steps: cell detection, registration to a common atlas, and injection site recovery. For maximum detection accuracy, cell detection was performed on unregistered image volumes, and the detected cells were then transformed to atlas coordinates.

Before analysis, datasets were manually curated by stringent quality control standards. Each brain was screened for (1) clearing quality, (2) significant tissue deformation from extraction process, (3) viral spread from injection site, (4) antibody penetration, (5) blending artifacts related to microscope misalignment, (6) injection site within target location, (7) successful registration, and (8) convolutional neural network (CNN) overlay of detected cells with brain volume in signal channel. Because of the relatively high concentration of antibody solution needed for brain-wide immunohistochemical staining, non-specific fluorescence was apparent at the edges of tissue, *i.e.* outside of the brain and ventricles, in the form of punctate labeling not of cell origin. We computationally removed a border at the brain edge at the ventricles to remove false positives, at the cost of loss of some true positives (skimage.morphology.binary_erosion, **Table S3**). For neocortical layer studies, a subregion of the primary somatosensory area: “primary somatosensory area, unassigned” in PMA did not have layer-specific mapping in Allen Atlas space and was removed from consideration.

##### Injection site recovery and segmentation

Injection sites were identified in both H129 and PRV studies by co-injecting CTB with virus (**Figure S13c**) and in c-Fos studies using ArchT-GFP expression. Post-registered light-sheet volumes of the injection channel were segmented to obtain voxel-by-voxel injection-site reconstructions. Volumes were Gaussian-blurred (3 voxel blurring parameter). All voxels less than 3 standard deviations above the mean were removed. The single largest connected component was considered the injection site (scipy.ndimage.label, SciPy 1.1.0 (Virtanen et al., 2020). CTB was selected for injection site labelling for transsynaptic tracing as it does not affect the spread of alpha-herpesviruses and its greater diffusion due to its smaller size overestimates the viral injection size by as much as two-fold (Aston-Jones and Card, 2000; Chen et al., 1999). Although typically used as a tracer itself, used during the small window of ~80 hpi, it did not have time to spread significantly. **Figure S3c-e** shows the percentage of cerebellum covered by at least one injection in each of the three datasets. Lobules I-III, flocculus, and paraflocculus were not targeted because of the primary focus on neocerebellum. and because their anatomical location and the sagittal sinus presented injection challenges. Interactive three-dimensional visualization of heatmaps and injection sites are available at https://brainmaps.princeton.edu/2021/05/pisano_viral_tracing_injections/.

##### Automated detection of transsynaptically labeled neurons

Each brain generated a dataset exceeding 100 gigabytes. To automate cell detection, we trained a three-dimensional convolutional neural network (CNN) to identify neurons. A CNN with U-Net architecture running on a GPU-based cluster was trained by supervised learning using more than 3600 human-annotated cell centers (**Figure 1e**; **Table 2**). To optimize cell detection for scalability, whole-brain light-sheet volumes (typically 100-150 GB 16-bit volumes) were chunked into approximately 80 compressed 32-bit TIF volumes per brain, with an overlap of 192 × 192 × 20 voxels in xyz between each volume, and stored on a file server.

For deploying the custom-trained cell-detection neural network, the file server streamed the volumes to a GPU cluster for segmentation. Once the segmentation was completed, the soma labels were reconstructed across the entire brain volume from the segmented image on a CPU cluster by calculating the maximum between the overlapping segments of each volume. The reconstructed brain volumes after segmentation were stored as memory-mapped arrays on a file server. Coordinates of cell centers from the reconstructed volumes were obtained by thresholding, using the established threshold from training evaluation, and connected-component analysis. Additionally, measures of detected cell perimeter, sphericity, and number of voxels it spans in the z-dimension were calculated by connected-component analysis for further cell classification if needed. The final output consisted of a comma-separated-value file that included the xyz coordinates as well as measures of perimeter, sphericity, and number of voxels in the z-dimension for each detected cell in the brain volume.

##### Convolutional neural network training

Supervised learning using CNN is useful in complex classification tasks when a sufficient amount of training data is available. Annotated training volumes were generated by selecting volumes at least 200 × 200 × 50 pixels (XYZ) from full-sized cell channel volumes. To ensure training data were representative of the animal variability across the whole-brain, training volumes were selected from different anatomical regions in different brains with various amounts of labeling (**Table S1** for dataset description). Annotations were recorded by marking cell centers using ImageJ (Schmid et al., 2010). To generate labeled volumes, Otsu’s thresholding method (skimage.filters.threshold_otsu, Scikit-Image (van der Walt et al., 2014) 0.13.1) (van der Walt et al., 2014) was applied within windows (30 × 30 × 8 voxels, XYZ) around each center to label soma. Using annotated volumes, we trained the previously mentioned CNN (Gornet et al., 2019; Lee et al., 2017) (github.com/PrincetonUniversity/BrainPipe). A 192 × 192 × 20 CNN window size with 0.75 strides was selected. The training dataset was split into a 70% training, 20% validation, and 10% testing subset. Training occurred on a SLURM-based GPU cluster. During training, the CNN was presented with data from the training dataset, and after each iteration its performance was evaluated using the validation dataset. Loss values, which measure learning by the CNN, stabilized at 295,000 training iterations, at which point training was stopped to prevent overfitting, *i.e.* the possibility that the neural network learns particular training examples rather than learning the category.

##### Evaluation of CNN

To determine CNN performance on H129 data, we calculated an F1 score (Goutte and Gaussier, 2005). First, we needed to compare CNN output with our ground truth annotations by quantifying true positives (TP), false negatives (FN), and false positives (FP). We defined human-annotation as ground truth, consistent with the machine learning field (Yao et al., 2007). Our neural network architecture produced a voxel-wise 0 (background) to 1 (cell) probability output. To determine a threshold value for binarization of the continuous 0-1 CNN-output values, F1 scores as a function of thresholds between 0 and 1 were determined (**FIgure 1f**). Connected-component analysis (scipy.ndimage.label) grouped islands of nonzero voxels to identify each island as a putative cell. Pairwise Euclidean distances (scipy.spatial.distance.euclidean) were calculated between CNN-predicted cell centers and human-annotated ground truth centers. Bipartite matching serially paired closest predicted and ground truth centers, removing each from unpaired pools. Unmatched predicted or ground truth centers were considered FPs or FNs, respectively. Prediction-ground truth center pairs with a Euclidean distance greater than 30 voxels (~49 µm) were likely inaccurate and not paired.

The F1 score was defined as the harmonic average of precision and recall. Precision is the number of correct positive results divided by the number of all positive results returned by the classifier, *i.e.* TP/ (TP+FP). Recall is the number of correct positive results divided by the number of all samples that should have been identified as positive, *i.e.* TP/ (TP+FN). The F1 score, harmonic mean of precision and recall (Chinchor, 1992), reaches its best value at 1 (perfect precision and recall) and worst at 0. Performance at different likelihood thresholds was plotted as a receiver operating characteristic curve of precision and recall (**Figure 1f**). A threshold likelihood of 0.6 was found to maximize this score. Using a 20-voxel cutoff instead of 30 gave 0.849 and 0.875 for human-CNN and human-human F1 scores, respectively. To determine CNN performance metrics, the testing dataset, which the network had yet to be exposed to, was finally run using the established threshold, producing an F1 score of 0.864. To generate the precision-recall curve, precision and recall values were calculated between thresholds of 0.002 and 0.998 with a step size of 0.002. Values of precision and 1−recall were used to plot the curve. The area-under-curve of the precision-recall curve was calculated using the composite trapezoidal rule (numpy.trapz). Querying the CNN gave an F1 score of 0.864, close to the human-vs.-human F1 score of, 0.891, indicating that the CNN had successfully generalized.

#### C-FOS MAPPING EXPERIMENT

##### c-Fos mapping after optogenetic perturbation

Neural activity has been shown to increase c-Fos, an immediate-early gene product (Martinez et al., 2002). Mapping of c-Fos expression used L7-Cre +/− (n=10) and −/− (n=8) mice (males, B6; 129-Tg (Pcp2-cre)2Mpin/J, 004146, The Jackson Laboratory, Bar Harbor, Maine, bred in-house, 56 days or older). L7-Cre mice express Cre recombinase exclusively in Purkinje neurons (Barski et al., 2000). AAV1-CAG-FLEX-ArchT-GFP (UNC Vector Core, deposited by Dr. Ed Boyden, 4×10^12^ vg/ml, AV5593B lot number, 500 nl/injection 250 µm deep perpendicular to tissue) was pressure-injected into four locations in lobule VIa/b.

Unlike transsynaptic tracing, where each individual animal can be used to test connectivity of a different cerebellar region, this experimental paradigm required targeting one cerebellar region (lobule VI) to achieve sufficient statistical power. To ensure adequate power in this experiment our sample sizes were at least double the size in the original studies developing this methodology (Renier et al., 2016). We selected lobule VI as the target given prior nonmotor findings associated with this lobule (Badura et al., 2018).

After virus injection, a cover slip (round 3 mm, #1 thickness, Warner Instruments 64–0720) was used to cover the craniotomy and a custom titanium plate for head fixation (Kloth et al., 2015) was attached using dental cement (S396, Parkell, Brentwood, NY). Mice were allowed to recover after surgery for 4 weeks and then were habituated to a head-fixed treadmill (Kloth et al., 2015) for three days, 30 minutes per day. On the last day of habituation, ArchT-GFP expression was confirmed using wide-field fluorescence microscopy. The following day, mice were again placed on the treadmill and a 200 µm fiber (M200L02S-A, Thorlabs, Newton, NJ) was placed directly over the cranial window for optogenetic stimulation with 532 nm laser (1 Hz, 250 ms pulse-width, 56 mW before entering fiber, 1 hr, GR-532-00200-CWM-SD-05-LED-0, Opto Engine, Midvale, UT). We determined the appropriate stimulation power using test animals, prepared in the same manner as previously described, but also with electrophysiological recordings. We titrated our stimulus on these test animals (not included in the manuscript cohort) to ensure it did not produce movements while producing reliable silencing of Purkinje cells (PCs) during light stimulus, but without significant silencing after termination of the light-stimulus (**Figure S11**). The experimental configuration delivered light from illumination from outside the brain, which was therefore attenuated through the air, coverslip, and brain tissue, leading to light scattering and heat dissipation. This made power requirements higher than other published studies (Choe et al., 2018).

A methodological limitation was the inability to determine whether differences were a result of a direct circuit effect, as opposed to a change in brain state that might affect animal behavior or sensory perception. As a control, we utilized a head-fixed treadmill approach, limiting types of animal movement. We quantified treadmill speed and arm movement to measure gross motor or behavioral changes.

We compared Cre +/− and Cre −/− animals, and ensured that our control animals (no Cre, no channelrhodopsin) received the same surgery, injection, coverslip placement, and head mount, and were placed on the same wheel and received the laser placement and activation. Animals were also cagemates (mixed Cre +/− and Cre −/− in each cage) since birth, providing natural blinding of condition to the experimenter.

Mice were then individually placed into a clean cage, kept in the dark for one hour, and perfused as described previously. Brains were fixed overnight in 10% formalin (4% formaldehyde) before beginning the iDISCO+ clearing protocol. Both ArchT-expressing mice and non-expressing mice received cranial windows, habituation, and photostimulation.

For behavioral quantification (**Figure S11**), videos were imported into ImageJ using QuickTime for Java library, and images converted into grayscale. Timing of optogenetic stimulation was confirmed by analysis of pixel intensity over optical fiber connection to implanted cannulae. Forelimb kinematic data and treadmill speed were analyzed by Manual Tracking plugin. For arm movement stimulation movements with forearm moving forward (initial positive slope) from forearm moving backwards (initial negative slope).

##### Electrophysiological confirmation of ArchT expression in Purkinje cells

To confirm that ArchT was optically activatable in PCs, photostimulation was done during patch-clamp recording in acutely prepared brain slices. Brain slices were prepared from three 10 week-old male Pcp2-cre mice (B6.Cg-Tg (Pcp2-cre)3555Jdhu/J, 010536, The Jackson Laboratory), two weeks after injection with AAV1-CAG-FLEX-ArchT-GFP. Mice were deeply anesthetized with Euthasol (0.06 ml/30 g), decapitated, and the brain removed. The isolated whole brains were immersed in ice-cold carbogenated NMDG ACSF solution (92 mM N-methyl D-glucamine, 2.5 mM KCl, 1.25 mM NaH2PO4, 30 mM NaHCO3, 20 mM HEPES, 25 mM glucose, 2 mM thiourea, 5 mM Na-ascorbate, 3 mM Na-pyruvate, 0.5 mM CaCl2, 10 mM MgSO4, and 12 mM N-acetyl-L-cysteine, pH adjusted to 7.3–7.4). Parasagittal cerebellar brain slices 300 μm thick were cut using a vibratome (VT1200s, Leica Microsystems, Wetzlar, Germany), incubated in NMDG ACSF at 34°C for 15 minutes, and transferred into a holding solution of HEPES ACSF (92 mM NaCl, 2.5 mM KCl, 1.25 mM NaH2PO4, 30 mM NaHCO3, 20 mM HEPES, 25 mM glucose, 2 mM thiourea, 5 mM Na-ascorbate, 3 mM Na-pyruvate, 2 mM CaCl2, 2 mM MgSO4 and 12 mM N-acetyl-L-cysteine, bubbled at room temperature with 95% O2 and 5% CO2). During recordings, slices were perfused at a flow rate of 4–5 ml/min with a recording ACSF solution (120 mM NaCl, 3.5 mM KCl, 1.25 mM NaH2PO4, 26 mM NaHCO3, 1.3 mM MgCl2, 2 mM CaCl2 and 11 mM D-glucose) and continuously bubbled with 95% O2 and 5% CO2.

Whole-cell recordings were performed using a Multiclamp 700B (Molecular Devices, Sunnyvale, CA) using pipettes with a resistance of 3–5 ΩM filled with a potassium-based mM MgCl2, 2 mM Mg-ATP and 0.3 mM Na-GTP, pH adjusted to 7.2 with KOH). Purkinje neurons expressing YFP were selected for recordings. Photostimulation parameters used were 525 nm, 0.12 mW/mm², and 250 ms pulses at 1 Hz.

##### Light-sheet microscopy for c-Fos imaging

Opaque magnets (D1005A-10 Parylene, Supermagnetman, Pelham, AL) were glued to ventral brain surfaces in the horizontal orientation and imaged using a light-sheet microscope as described previously. Version 5.1.293 of the ImSpector Microscope controller software was used. ArchT-GFP injection volumes were acquired using the 561 nm excitation filter. Cellular imaging of c-Fos expressing cells was acquired using 640 nm excitation filter at 5.0 µm/pixel (1x magnification, 1.3x objective, 0.1 numerical aperture, 9.0 mm working distance, 12.0 × 12.0 mm field of view, LVMI-Fluor 1.3x, LaVision Biotech) with a 3 µm step-size using an excitation sheet with a numerical aperture of 0.010. This resolution was selected to allow whole-brain imaging using ClearMap without tiling artifacts. To speed up acquisition, the autofluorescence channel and injection channels were acquired separately with a shorter exposure time than the cell channel. The left and right horizontal focus was shifted towards the side of the emitting sheet. Left and right images were then sigmoidally blended before analysis. In order to maximize field of view, some olfactory areas were not completely represented in images and were removed from analysis. Five brains were reimaged a second time due to ventricular imaging artifacts.

##### Automated detection of c-Fos expressing cells

Detection of c-Fos expressing cells after optogenetic stimulation was done using ClearMap software for c-Fos detection (Renier et al., 2016) modified to run on high performance computing clusters (“ClearMapCluster”). Analysis parameters used were: removeBackgroundParameter_size= (5,5), findExtendedMaximaParameter_size= (5,5), findExtendedMaximaParameter_threshold=0, findIntensityParameter_size= (3,3,3), detectCellShapeParameter_threshold=105. Cell detection parameters were optimized by two users iterating through a set of varying ClearMap detection parameters and selecting those that minimized false positives while labelling only c-Fos positive neurons with high signal-to-noise ratio.

### Quantification and Statistical Analysis

#### Statistical analysis of registration precision

To quantify atlas registration accuracy, blinded users labeled readily identifiable points in the PMA (similar to Ref. (Sergejeva et al., 2015)in four sets of unregistered, affine-only registered, and fully registered volumes (**Figure 1,h**, **Figure S2i**). This allowed for quantification of landmark distances (Sergejeva et al., 2015) between the PMA and brains at different stages of registration. Estimated standard deviations are defined as the median absolute deviation (MAD) divided by 0.6745. MADs were calculated with Statsmodels (Seabold and Perktold, 2010) 0.9.0 (statsmodels.robust.mad). Eleven blinded users annotated a total of 69 points. One measurement was considered to be user error and was dropped from the theoretical-limit measurements, as it was over 12 times the median of the other measures.

After registration, the median Euclidean distance from complementary points in the PMA was 93 ± 36 µm (median ± estimated standard deviation). Blinded users determined points in the same volume twice to establish an intrinsic minimum limit of 49 ± 40 µm. Assuming that uncertainties sum as independent variables, the estimated accuracy of registration was √ (93^2^-49^2^)=79 μm, or 4 voxels.

#### Statistical analysis of transsynaptic tracing data

For initial inspection of thalamic or neocortical neurons, each injected brain was sorted by cerebellar region with the greatest volume fraction of the injection (as in Ref. (Badura et al., 2018); this region was defined as the primary injection site.

Two primary methods of quantification were used, fraction of all labeled neurons in the thalamus or neocortex, and density within particular structures. Fraction of neurons is defined as the total labelled neuron count within a structure (*e.g.* VA-L) divided by the main parent structure (*e.g.* thalamus). This number is useful as it gives relative target projection strength relative to other projection strengths within the parent structure. However, this does not provide information in non-relative terms. Density, defined as total labelled neurons divided by volume of the structure, takes into account the relative sizes for each structure, allowing for more absolute comparisons of recipient structures. In anterograde examples, density therefore provides information on the concentration of influence a cerebellar region may have on a target structure. Unless otherwise noted in the results, density analyses utilized mean and standard deviations. Further details are provided in Results, The cerebellum sends output to a wide range of thalamic targets and Table S3. For cohort sizes see Table S1.

Results from different experimental animals were analyzed in two ways. First, the data are displayed in a column-by-column manner in **Figures 3, 5, 7, S7, S8b, S9 and S14a,b**. Second, generalized linear models (GLM) were used to identify specific topographical relationships that are shared among animals and display them as connection weights that can account for the overall pattern of results. In this way, we were able to efficiently display the results of three cohorts (**Table S1**) consisting of 23 animals for disynaptic H129, 33 animals for trisynaptic H129, and 25 animals for disynaptic PRV.

#### Generalized linear model analysis

Contribution of each cerebellar meta-lobule to viral spread in each neocortical or thalamic region was fitted to a generalized linear model (GLM) consisting of an inhomogeneous Poisson process as a function of seven targeted cerebellar regions (“meta-lobules”). The predictor variables were *x_j_*, where *x_j_* is defined as the fraction of the total injection to be found in the j-th meta-lobule, such that *∑x_j_* = 1 *y_k_* defined as the fraction of the total number of cells in the entire neocortex (or thalamus) to be found in the k-th region. For the resulting fit coefficients β*_jk_*, the change in 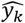 arising from a unit change in *x_j_* is e^β*jk*^. In **Figures 4f, 6f**, and **8f**, the heatmap indicates a measure of confidence, defined as the coefficient β*_jk_* divided by the coefficient’s standard error.

To determine statistically significant weights, we compared significant weights computed from the t-stats of the coefficients with those observed in a shuffle-based null model in which predictors were shuffled uniformly at random (n = 1,000). We found that the true number of positive significant weights was significantly greater than that expected under the null model with a one-sided, independent t-test of the coefficients (p < 0.05). In **Figure 6**, the neocortical region “Frontal pole, cerebral cortex” was excluded from generalized linear model analysis due to zero counts across all brains for the region.

#### AAV DCN Injection Immunofluorescence Image Analysis

Image stacks were deconvolved using Huygens software (Scientific Volume Imaging). With a custom-written Fiji-scripts (ImageJ) we identified putative synaptic contacts, *i.e.* YFP-positive varicosities that colocalized with vGluT2-staining, following the same analysis pipeline as Ref. (Gornati et al., 2018), (script available upon reasonable request). The color channels (YFP and Cy5) of the images were split to get separate stacks. The YFP and Cy5 channels were Gaussian blurred (sigma = 1) and selected by a manually set threshold. A binary open function was done on both images (iterations = 4, count = 2) and objects were removed if their size was <400 pixels (YFP). A small dilatation was done on the red image (iteration = 1, count = 1). With the image calculator an ‘and-operation’ was done using the binary red and green image. The values 255 (white) of the binary YFP image were set to 127. This image and the result of the AND-operation were combined by an OR-operation. The resulting image was measured with the 3D-object counter plugin for volumes and maximum intensities. Only objects containing pixels with an intensity of 255 (overlap) are taken in account for particle analysis. Estimation of synapse density (number of terminals/area μm^3^) was obtained for each image by dividing the number of terminals by the image area (DeKosky and Scheff, 1990). Counts of vGluT2 and YFP co-labeled varicosities in thirteen randomly picked regions in VM, VA-L, and CL (each region 100×100×5 microns) were strongly correlated with average YFP brightness for that same region (r=+0.94, t=8.76, p<0.0001; **Figure S4c-e**). Therefore we used summed brightness as a measure of total innervation. Summed brightness was defined as the total fluorescence within a nucleus, summed across all sections where the nucleus was present. Regression between AAV and HSV-H129 density was performed using two-sided Pearson’s regression was performed (R, cor.test).

#### Statistical analysis of c-Fos data

Cell and density heat maps and p-value maps were generated using ClearMap. Projected p-value maps were generated by binarizing the p-value maps and counting non-zero voxels in z; color bar thresholding displayed greater than 25% for coronal and 27% for sagittal sections of the z-distance. Injection sites were segmented and aligned in the manner described previously. Activation ratio was defined as the mean number of cells in an anatomical area across experimental brains divided by the mean number of cells in the same anatomical area in control brains. To compare the c-Fos activation data with transsynaptic tracing data across the major divisions in the neocortex, linear-least squares regression (scipy.stats.linregress, two-sided) were calculated using mean viral-labeling neocortical densities with H129-VC22 injections (80 hpi) were compared with the mean cell density ratio of c-Fos stimulation vs control groups. The Mann-Whitney U test (two-tailed; scipy.stats.mannwhitneyu, SciPy (Virtanen et al., 2020) 1.1.0), a nonparametric version of the t-test, was used to determine statistical significance (p < 0.05) between control and experimental brain regions in c-Fos studies. Pearson’s regression was performed (scipy.stats.pearsonr). Further details including cohort size can be found in Table S1 and ClearMapCluster settings can be found in Methods, Automated detection of c-Fos expressing cells.

#### SOFTWARE

Analysis pipelines were run using custom code written for Python 3+, available at github.com/PrincetonUniversity/BrainPipe and github.com/PrincetonUniversity/ClearMapCluster (see Key Resources Table). Unless otherwise noted, analyses and plotting were performed in Python 2.7+. DataFrame manipulations were done using Numpy (Oliphant, 2015) 1.14.3 and Pandas (McKinney and Others, 2010) 0.23.0. Plotting was done with Matplotlib (Hunter, 2007) 2.2.2 and Seaborn (Waskom et al., 2014) 0.9.0. Image loading, manipulation and visualization was done using Scikit-Image (van der Walt et al., 2014) 0.13.1 and SimpleITK (Lowekamp et al., 2013) 1.0.0. SciPy (Virtanen et al., 2020) 1.1.0 was used for statistical analyses. Hierarchical clustering analysis was performed using Seaborn (Waskom et al., 2014) 0.9.0 and Scikit-Learn (Pedregosa et al., 2011) 0.19.1 was used for hierarchical agglomerative clustering (average metric, Ward’s method). Multidimensional scaling was done in MATLAB 2019b. Coefficients and standard errors for the generalized linear model were obtained by fitting the model using the statsmodels 0.9.0 package in Python 3.7.1 (Badura et al., 2018).

## Supplemental Videos

**Video S1: Princeton Mouse Atlas.** Three-dimensional rendering of the Princeton Mouse Atlas, a light-sheet mouse brain atlas with complete cerebellum and compatible with the Allen Brain Mouse Atlas.

**Video S2: Princeton Mouse Atlas Annotations.** Three-dimensional rendering of the Princeton Mouse Atlas annotations. Colors represent different structures annotated in the atlas.

## Acknowledgments

We thank Aleksandra Badura for advice on experimental design, Lynn Enquist for discussion and PRV-Bartha 152 (CNNV, P40 OD010996), James Gornet for neural network implementation assistance, Nicolas Renier and Kelly Seagraves for tissue-clearing optimization, Stephan Thiberge for microscopy help, and Shruthi Deivasigamani, Joseph Gotto, Joyce Lee, Laura Lynch, Caroline Jung, Sanjeev Janarthanan, Dafina Pacuku, Federico Uquillas, and Thaddeus Weigel for technical assistance, and Pavel Osten for project advice. This work was supported by NIH R01 NS045193, R01 MH115750, and U19 NS104648 (S.W.), F31 NS089303 (T.P.), P40 OD010996 (E.E.), R21 DC018365 (B.R.), and P20GM103408 (E.H. and B.R.), Netherlands Organization for Scientific Research Veni ZonMW, 91618112 (H.-J.B), Erasmus MC Fellowship 106958 (H.-J.B), and the New Jersey Council on Brain Injury Research (J.V.).

## Author contributions

T.P., M.K., H.-J.B., and S.W. conceived and designed studies. T.P., D.B., and J.V. performed virus injections and prepared tissue. Z.D. and T.P. imaged tissue and ran the computational data analysis pipeline for light-sheet data. T.P., Z.D., and H.-J. B. performed data analysis and prepared figures. E.E. constructed HSV vectors. K.U.V. and T.P. designed image analysis algorithms. A.H. analyzed images and built visualizations. M.K., J.L., and T.P. performed c-Fos studies. E.H. and B.R. performed AAV-TRN studies, B.R. collected and analyzed images. H.-J. B., N. de O. and F.H. performed AAV-DCN studies, collected and analyzed images. T.P. and S.W. wrote the initial draft of the manuscript, which was edited by all authors.

## Inclusion and Diversity

We worked to ensure sex balance in the selection of non-human subjects. One or more of the authors of this paper self-identifies as a member of the LGBTQ+ community. One or more of the authors of this paper self-identifies as living with a disability.

## References

Allen Institute for BrainScience (2012). Technical white paper informatics data processing for the allen developing mouse brain atlas.

Ando, N., Izawa, Y., and Shinoda, Y. (1995). Relative contributions of thalamic reticular nucleus neurons and intrinsic interneurons to inhibition of thalamic neurons projecting to the motor cortex. J. Neurophysiol. 73, 2470–2485.

Angaut, P., Guilbaud, G., and Reymond, M.-C. (1968). An electrophysiological study of the cerebellar projections to the nucleus ventralis lateralis of thalamus in the cat. I. Nuclei fastigii et interpositus. The Journal of Comparative Neurology 134, 9–19.

Angaut, P., Cicirata, F., and Serapide, F. (1985). Topographic organization of the cerebellothalamic projections in the rat. An autoradiographic study. Neuroscience 15, 389–401.

Aoki, S., Coulon, P., and Ruigrok, T.J.H. (2017). Multizonal Cerebellar Influence Over Sensorimotor Areas of the Rat Cerebral Cortex. Cereb. Cortex 29, 598–614.

Apps, R. (2000). Gating of climbing fibre input to cerebellar cortical zones. Prog. Brain Res. 124, 201–211.

Apps, R., and Hawkes, R. (2009). Cerebellar cortical organization: a one-map hypothesis. Nat. Rev. Neurosci. 10, 670–681.

Aston-Jones, G., and Card, J.P. (2000). Use of pseudorabies virus to delineate multisynaptic circuits in brain: opportunities and limitations. J. Neurosci. Methods 103, 51–61.

Aumann, T.D., and Horne, M.K. (1996). Ramification and termination of single axons in the cerebellothalamic pathway of the rat. J. Comp. Neurol. 376, 420–430.

Aumann, T.D., Rawson, J.A., Finkelstein, D.I., and Horne, M.K. (1994). Projections from the lateral and interposed cerebellar nuclei to the thalamus of the rat: a light and electron microscopic study using single and double anterograde labelling. J. Comp. Neurol. 349, 165–181.

Aumann, T.D., Ivanusic, J., and Horne, M.K. (1998). Arborisation and termination of single motor thalamocortical axons in the rat. The Journal of Comparative Neurology 396, 121–130.

Badura, A., Verpeut, J.L., Metzger, J.W., Pereira, T.D., Pisano, T.J., Deverett, B., Bakshinskaya, D.E., and Wang, S.S.-H. (2018). Normal cognitive and social development require posterior cerebellar activity. Elife 7.

Balsters, J.H., Laird, A.R., Fox, P.T., and Eickhoff, S.B. (2014). Bridging the gap between functional and anatomical features of cortico-cerebellar circuits using meta-analytic connectivity modeling. Hum. Brain Mapp. 35, 3152–3169.

Barski, J.J., Dethleffsen, K., and Meyer, M. (2000). Cre recombinase expression in cerebellar Purkinje cells. Genesis 28, 93–98.

Bostan, A.C., and Strick, P.L. (2018). The basal ganglia and the cerebellum: nodes in an integrated network. Nat. Rev. Neurosci. 19, 338–350.

Bria, A., and Iannello, G. (2012). TeraStitcher - a tool for fast automatic 3D-stitching of teravoxel-sized microscopy images. BMC Bioinformatics 13, 316.

Bruno, R.M., and Sakmann, B. (2006). Cortex is driven by weak but synchronously active thalamocortical synapses. Science 312, 1622–1627.

Buckner, R.L., Krienen, F.M., Castellanos, A., Diaz, J.C., and Yeo, B.T.T. (2011). The organization of the human cerebellum estimated by intrinsic functional connectivity. J. Neurophysiol. 106, 2322–2345.

Callaway, E.M. (2008). Transneuronal circuit tracing with neurotropic viruses. Curr. Opin. Neurobiol. 18, 617–623.

Card, J.P., Enquist, L.W., Miller, A.D., and Yates, B.J. (1997). Differential tropism of pseudorabies virus for sensory neurons in the cat. J. Neurovirol. 3, 49–61.

Card, J.P., Enquist, L.W., and Moore, R.Y. (1999). Neuroinvasiveness of pseudorabies virus injected intracerebrally is dependent on viral concentration and terminal field density. J. Comp. Neurol. 407, 438–452.

Carta, I., Chen, C.H., Schott, A.L., Dorizan, S., and Khodakhah, K. (2019). Cerebellar modulation of the reward circuitry and social behavior. Science 363.

Cavdar, S., Onat, F.Y., Yananli, H.R., Sehirli, U.S., Tulay, C., Saka, E., and Gürdal, E. (2002). Cerebellar connections to the rostral reticular nucleus of the thalamus in the rat. J. Anat. 201, 485–491.

Chan-Palay, V. (2013). Cerebellar Dentate Nucleus: Organization, Cytology and Transmitters (Springer).

Chen, S., Yang, M., Miselis, R.R., and Aston-Jones, G. (1999). Characterization of transsynaptic tracing with central application of pseudorabies virus. Brain Res. 838, 171–183.

Chinchor, N. (1992). MUC-4 evaluation metrics. Proceedings of the 4th Conference on Message Understanding - MUC4 ’92.

Choe, K.Y., Sanchez, C.F., Harris, N.G., Otis, T.S., and Mathews, P.J. (2018). Optogenetic fMRI and electrophysiological identification of region-specific connectivity between the cerebellar cortex and forebrain. Neuroimage 173, 370–383.

Cicirata, F., Angaut, P., Serapide, M.F., and Panto, M.R. (1990). Functional organization of the direct and indirect projection via the reticularis thalami nuclear complex from the motor cortex to the thalamic nucleus ventralis lateralis. Exp. Brain Res. 79, 325–337.

Constantinople, C.M., and Bruno, R.M. (2013). Deep cortical layers are activated directly by thalamus. Science 340, 1591–1594.

Conte, W.L., Kamishina, H., and Reep, R.L. (2009). Multiple neuroanatomical tract-tracing using fluorescent Alexa Fluor conjugates of cholera toxin subunit B in rats. Nat. Protoc. 4, 1157–1166.

Cook, A.A., Fields, E., and Watt, A.J. (2020). Losing the Beat: Contribution of Purkinje Cell Firing Dysfunction to Disease, and Its Reversal. Neuroscience 462, 247–261.

Courchesne, E., Yeung-Courchesne, R., Press, G.A., Hesselink, J.R., and Jernigan, T.L. (1988). Hypoplasia of cerebellar vermal lobules VI and VII in autism. N. Engl. J. Med. 318, 1349–1354.

Courchesne, E., Karns, C.M., Davis, H.R., Ziccardi, R., Carper, R.A., Tigue, Z.D., Chisum, H.J., Moses, P., Pierce, K., Lord, C., et al. (2001). Unusual brain growth patterns in early life in patients with autistic disorder: an MRI study. Neurology 57, 245–254.

DeKosky, S.T., and Scheff, S.W. (1990). Synapse loss in frontal cortex biopsies in Alzheimer’s disease: correlation with cognitive severity. Ann. Neurol. 27, 457–464.

Destexhe, A. (2000). Modelling corticothalamic feedback and the gating of the thalamus by the cerebral cortex. J. Physiol. Paris 94, 391–410.

Deverett, B., Koay, S.A., Oostland, M., and Wang, S.S.-H. (2018). Cerebellar involvement in an evidence-accumulation decision-making task. Elife 7.

Dragunow, M., and Faull, R. (1989). The use of c-fos as a metabolic marker in neuronal pathway tracing. J. Neurosci. Methods 29, 261–265.

Fujita, H., Kodama, T., and du Lac, S. (2020). Modular output circuits of the fastigial nucleus for diverse motor and nonmotor functions of the cerebellar vermis. Elife 9.

Gao, Z., Davis, C., Thomas, A.M., Economo, M.N., Abrego, A.M., Svoboda, K., De Zeeuw, C.I., and Li, N. (2018). A cortico-cerebellar loop for motor planning. Nature 563, 113–116.

Garner, J.A., and LaVail, J.H. (1999). Differential anterograde transport of HSV type 1 viral strains in the murine optic pathway. J. Neurovirol. 5, 140–150.

Gornati, S.V., Schäfer, C.B., Eelkman Rooda, O.H.J., Nigg, A.L., De Zeeuw, C.I., and Hoebeek, F.E. (2018). Differentiating Cerebellar Impact on Thalamic Nuclei. Cell Rep. 23, 2690–2704.

Gornet, J., Venkataraju, K.U., Narasimhan, A., Turner, N., Lee, K., Sebastian Seung, H., Osten, P., and Sümbül, U. (2019). Reconstructing neuronal anatomy from whole-brain images. In IEEE 16th International Symposium on Biomedical Imaging, (arXiv),.

Goutte, C., and Gaussier, E. (2005). A Probabilistic Interpretation of Precision, Recall and F-Score, with Implication for Evaluation. In Advances in Information Retrieval, (Springer Berlin Heidelberg), pp. 345–359.

Granstedt, A.E., Bosse, J.B., Thiberge, S.Y., and Enquist, L.W. (2013). In vivo imaging of alphaherpesvirus infection reveals synchronized activity dependent on axonal sorting of viral proteins. Proc. Natl. Acad. Sci. U. S. A. 110, E3516–E3525.

Guillery, R.W., Feig, S.L., and Lozsádi, D.A. (1998). Paying attention to the thalamic reticular nucleus. Trends Neurosci. 21, 28–32.

Hashimoto, M., Takahara, D., Hirata, Y., Inoue, K.-I., Miyachi, S., Nambu, A., Tanji, J., Takada, M., and Hoshi, E. (2010). Motor and non-motor projections from the cerebellum to rostrocaudally distinct sectors of the dorsal premotor cortex in macaques. Eur. J. Neurosci. 31, 1402–1413.

Hashimoto, M., Yamanaka, A., Kato, S., Tanifuji, M., Kobayashi, K., and Yaginuma, H. (2018). Anatomical Evidence for a Direct Projection from Purkinje Cells in the Mouse Cerebellar Vermis to Medial Parabrachial Nucleus. Front. Neural Circuits 12.

Heath, R.G., Dempesy, C.W., Fontana, C.J., and Myers, W.A. (1978). Cerebellar stimulation: effects on septal region, hippocampus, and amygdala of cats and rats. Biol. Psychiatry 13, 501–529.

Henschke, J.U., and Pakan, J.M. (2020). Disynaptic cerebrocerebellar pathways originating from multiple functionally distinct cortical areas. Elife 9.

Herkenham, M. (1980). Laminar organization of thalamic projections to the rat neocortex. Science 207, 532–535.

Hooks, B.M., Mao, T., Gutnisky, D.A., Yamawaki, N., Svoboda, K., and Shepherd, G.M.G. (2013). Organization of cortical and thalamic input to pyramidal neurons in mouse motor cortex. J. Neurosci. 33, 748–760.

Hubel, D.H., and Wiesel, T.N. (1965). Binocular interaction in striate cortex of kittens reared with artificial squint. J. Neurophysiol. 28, 1041–1059.

Hunnicutt, B.J., Long, B.R., Kusefoglu, D., Gertz, K.J., Zhong, H., and Mao, T. (2014). A comprehensive thalamocortical projection map at the mesoscopic level. Nat. Neurosci. 17, 1276–1285.

Hunter, J.D. (2007). Matplotlib: A 2D Graphics Environment. Comput. Sci. Eng. 9, 90–95.

Jones, E.G. (1975). Lamination and differential distribution of thalamic afferents within the sensory-motor cortex of the squirrel monkey. J. Comp. Neurol. 160, 167–203.

Jones, E.G. (2012). The Thalamus (Springer Science & Business Media).

Jones, E.G., and Burton, H. (1976). Areal differences in the laminar distribution of thalamic afferents in cortical fields of the insular, parietal and temporal regions of primates. J. Comp. Neurol. 168, 197–247.

Kebschull, J.M., Richman, E.B., Ringach, N., Friedmann, D., Albarran, E., Kolluru, S.S., Jones, R.C., Allen, W.E., Wang, Y., Cho, S.W., et al. (2020). Cerebellar nuclei evolved by repeatedly duplicating a conserved cell-type set. Science 370.

Kelly, R.M., and Strick, P.L. (2003). Cerebellar loops with motor cortex and prefrontal cortex of a nonhuman primate. J. Neurosci. 23, 8432–8444.

Klapoetke, N.C., Murata, Y., Kim, S.S., Pulver, S.R., Birdsey-Benson, A., Cho, Y.K., Morimoto, T.K., Chuong, A.S., Carpenter, E.J., Tian, Z., et al. (2014). Independent optical excitation of distinct neural populations. Nat. Methods 11, 338–346.

Klein, S., and Staring, M. (2015). Image registration. In Elastix, the Manual, pp. 13–16.

Klein, S., Staring, M., Murphy, K., Viergever, M.A., and Pluim, J.P.W. (2010). elastix: a toolbox for intensity-based medical image registration. IEEE Trans. Med. Imaging 29, 196–205.

Kloth, A.D., Badura, A., Li, A., Cherskov, A., Connolly, S.G., Giovannucci, A., Bangash, M.A., Grasselli, G., Peñagarikano, O., Piochon, C., et al. (2015). Cerebellar associative sensory learning defects in five mouse autism models. Elife 4, e06085.

Kubo, R., Aiba, A., and Hashimoto, K. (2018). The anatomical pathway from the mesodiencephalic junction to the inferior olive relays perioral sensory signals to the cerebellum in the mouse. J. Physiol. 596, 3775–3791.

Lam, Y.-W., and Sherman, S.M. (2010). Functional organization of the somatosensory cortical layer 6 feedback to the thalamus. Cereb. Cortex 20, 13–24.

Lee, K., Zung, J., Li, P., Jain, V., and Sebastian Seung, H. (2017). Superhuman Accuracy on the SNEMI3D Connectomics Challenge.

Lee, K.H., Mathews, P.J., Reeves, A.M.B., Choe, K.Y., Jami, S.A., Serrano, R.E., and Otis, T.S. (2015). Circuit mechanisms underlying motor memory formation in the cerebellum. Neuron 86, 529–540.

Legg, C.R., Mercier, B., and Glickstein, M. (1989). Corticopontine projection in the rat: the distribution of labelled cortical cells after large injections of horseradish peroxidase in the pontine nuclei. J. Comp. Neurol. 286, 427–441.

Léna, C., and Popa, D. (2016). Cerebrocerebellar loops in the rodent brain. In The Neuronal Codes of the Cerebellum, (Elsevier), pp. 135–153.

Limperopoulos, C., Bassan, H., Gauvreau, K., Robertson, R.L., Jr, Sullivan, N.R., Benson, C.B., Avery, L., Stewart, J., Soul, J.S., Ringer, S.A., et al. (2007). Does cerebellar injury in premature infants contribute to the high prevalence of long-term cognitive, learning, and behavioral disability in survivors? Pediatrics 120, 584–593.

Lisberger, S.G. (2020). The Rules of Cerebellar Learning: Around the Ito Hypothesis. Neuroscience 462, 175–190.

Llinás, R.R., Leznik, E., and Urbano, F.J. (2002). Temporal binding via cortical coincidence detection of specific and nonspecific thalamocortical inputs: A voltage-dependent dye-imaging study in mouse brain slices. Proc. Natl. Acad. Sci. U. S. A. 99, 449–454.

Lowekamp, B.C., Chen, D.T., Ibáñez, L., and Blezek, D. (2013). The Design of SimpleITK. Front. Neuroinform. 7, 45.

Luo, Y., Fujita, H., Nedelescu, H., Biswas, M.S., Sato, C., Ying, S., Takahashi, M., Akita, K., Higashi, T., Aoki, I., et al. (2017). Lobular homology in cerebellar hemispheres of humans, non-human primates and rodents: a structural, axonal tracing and molecular expression analysis. Brain Struct. Funct. 222, 2449–2472.

Martinez, M., Calvo-Torrent, A., and Herbert, J. (2002). Mapping brain response to social stress in rodents with c-fos expression: a review. Stress 5, 3–13.

Marton, T.F., Seifikar, H., Luongo, F.J., Lee, A.T., and Sohal, V.S. (2018). Roles of Prefrontal Cortex and Mediodorsal Thalamus in Task Engagement and Behavioral Flexibility. J. Neurosci. 38, 2569–2578.

McGovern, A.E., Davis-Poynter, N., Farrell, M.J., and Mazzone, S.B. (2012). Transneuronal tracing of airways-related sensory circuitry using herpes simplex virus 1, strain H129. Neuroscience 207, 148–166.

McKinney, W., and Others (2010). Data structures for statistical computing in python. In Proceedings of the 9th Python in Science Conference, (Austin, TX), pp. 51–56.

Mihailoff, G.A., Kosinski, R.J., Azizi, S.A., and Border, B.G. (1989). Survey of noncortical afferent projections to the basilar pontine nuclei: a retrograde tracing study in the rat. J. Comp. Neurol. 282, 617–643.

Miranda-Saksena, M., Denes, C.E., Diefenbach, R.J., and Cunningham, A.L. (2018). Infection and Transport of Herpes Simplex Virus Type 1 in Neurons: Role of the Cytoskeleton. Viruses 10.

Mitchell, A., and Chakraborty, S. (2013). What does the mediodorsal thalamus do? Front. Syst. Neurosci. 7, 37.

Nakamura, H. (2018). Cerebellar projections to the ventral lateral geniculate nucleus and the thalamic reticular nucleus in the cat. J. Neurosci. Res. 96, 63–74.

Nassi, J.J., Cepko, C.L., Born, R.T., and Beier, K.T. (2015). Neuroanatomy goes viral! Front. Neuroanat. 9, 80.

Oliphant, T.E. (2015). Guide to NumPy, 2nd Edition (North Charleston, SC, USA: CreateSpace Independent Publishing Platform).

Ossowska, K. (2020). Zona incerta as a therapeutic target in Parkinson’s disease. J. Neurol. 267, 591–606.

Ozden, I., Dombeck, D.A., Hoogland, T.M., Tank, D.W., and Wang, S.S.-H. (2012). Widespread state-dependent shifts in cerebellar activity in locomoting mice. PLoS One 7, e42650.

Pedregosa, F., Varoquaux, G., Gramfort, A., Michel, V., Thirion, B., Grisel, O., Blondel, M., Prettenhofer, P., Weiss, R., Dubourg, V., et al. (2011). Scikit-learn: Machine Learning in Python. J. Mach. Learn. Res. 12, 2825–2830.

Phillipson, O.T. (1979). Afferent projections to the ventral tegmental area of Tsai and interfascicular nucleus: a horseradish peroxidase study in the rat. J. Comp. Neurol. 187, 117–143.

Pinault, D., Smith, Y., and Deschênes, M. (1997). Dendrodendritic and axoaxonic synapses in the thalamic reticular nucleus of the adult rat. J. Neurosci. 17, 3215–3233.

Pinto, D.J., Brumberg, J.C., and Simons, D.J. (2000). Circuit dynamics and coding strategies in rodent somatosensory cortex. J. Neurophysiol. 83, 1158–1166.

Popa, T., Russo, M., and Meunier, S. (2010). Long-lasting inhibition of cerebellar output. Brain Stimul. 3, 161–169.

Ray, M., Tang, R., Jiang, Z., and Rotello, V.M. (2015). Quantitative tracking of protein trafficking to the nucleus using cytosolic protein delivery by nanoparticle-stabilized nanocapsules. Bioconjug. Chem. 26, 1004–1007.

Renier, N., Wu, Z., Simon, D.J., Yang, J., Ariel, P., and Tessier-Lavigne, M. (2014). iDISCO: a simple, rapid method to immunolabel large tissue samples for volume imaging. Cell 159, 896–910.

Renier, N., Adams, E.L., Kirst, C., Wu, Z., Azevedo, R., Kohl, J., Autry, A.E., Kadiri, L., Umadevi Venkataraju, K., Zhou, Y., et al. (2016). Mapping of Brain Activity by Automated Volume Analysis of Immediate Early Genes. Cell 165, 1789–1802.

Roosendaal, T., and Selleri, S. (2004). The Official Blender 2.3 guide: free 3D creation suite for modeling, animation, and rendering (No Starch Press San Francisco, CA).

Ruigrok, T.J.H., Sillitoe, R.V., and Voogd, J. (2015). Chapter 9 - Cerebellum and Cerebellar Connections. In The Rat Nervous System (Fourth Edition), G. Paxinos, ed. (San Diego: Academic Press), pp. 133–205.

Saleeba, C., Dempsey, B., Le, S., Goodchild, A., and McMullan, S. (2019). A Student’s Guide to Neural Circuit Tracing. Front. Neurosci. 13, 897.

Salgado, S., and Kaplitt, M.G. (2015). The Nucleus Accumbens: A Comprehensive Review. Stereotact. Funct. Neurosurg. 93, 75–93.

Schmahmann, J.D., and Sherman, J.C. (1998). The cerebellar cognitive affective syndrome. Brain 121, 561–579.

Schmid, B., Schindelin, J., Cardona, A., Longair, M., and Heisenberg, M. (2010). A high-level 3D visualization API for Java and ImageJ. BMC Bioinformatics 11, 274.

Seabold, S., and Perktold, J. (2010). Statsmodels: Econometric and statistical modeling with python. In Proceedings of the 9th Python in Science Conference, (Scipy), p. 61.

Serapide, M.F., Pantó, M.R., Parenti, R., Zappalá, A., and Cicirata, F. (2001). Multiple zonal projections of the basilar pontine nuclei to the cerebellar cortex of the rat. J. Comp. Neurol. 430, 471–484.

Sergejeva, M., Papp, E.A., Bakker, R., Gaudnek, M.A., Okamura-Oho, Y., Boline, J., Bjaalie, J.G., and Hess, A. (2015). Anatomical landmarks for registration of experimental image data to volumetric rodent brain atlasing templates. J. Neurosci. Methods 240, 161–169.

Shamonin, D.P., Bron, E.E., Lelieveldt, B.P.F., Smits, M., Klein, S., Staring, M., and Alzheimer’s Disease Neuroimaging Initiative (2013). Fast parallel image registration on CPU and GPU for diagnostic classification of Alzheimer’s disease. Front. Neuroinform. 7, 50.

Shiroyama, T., Kayahara, T., Yasui, Y., Nomura, J., and Nakano, K. (1999). Projections of the vestibular nuclei to the thalamus in the rat: a Phaseolus vulgaris leucoagglutinin study. J. Comp. Neurol. 407, 318–332.

Sieveritz, B., García-Muñoz, M., and Arbuthnott, G.W. (2019). Thalamic afferents to prefrontal cortices from ventral motor nuclei in decision-making. Eur. J. Neurosci. 49, 646–657.

Smith, B.N., Banfield, B.W., Smeraski, C.A., Wilcox, C.L., Dudek, F.E., Enquist, L.W., and Pickard, G.E. (2000). Pseudorabies virus expressing enhanced green fluorescent protein: A tool for in vitro electrophysiological analysis of transsynaptically labeled neurons in identified central nervous system circuits. Proc. Natl. Acad. Sci. U. S. A. 97, 9264–9269.

Snider, R.S., and Maiti, A. (1976). Cerebellar contributions to the Papez circuit. J. Neurosci. Res. 2, 133–146.

Solari, S.V.H., and Stoner, R. (2011). Cognitive consilience: primate non-primary neuroanatomical circuits underlying cognition. Front. Neuroanat. 5, 65.

Song, C.K., Kay Song, C., Schwartz, G.J., and Bartness, T.J. (2009). Anterograde transneuronal viral tract tracing reveals central sensory circuits from white adipose tissue. American Journal of Physiology-Regulatory, Integrative and Comparative Physiology 296, R501–R511.

Stamatakis, A.M., Van Swieten, M., Basiri, M.L., Blair, G.A., Kantak, P., and Stuber, G.D. (2016). Lateral Hypothalamic Area Glutamatergic Neurons and Their Projections to the Lateral Habenula Regulate Feeding and Reward. J. Neurosci. 36, 302–311.

Stoodley, C.J., and Schmahmann, J.D. (2009). Functional topography in the human cerebellum: a meta-analysis of neuroimaging studies. Neuroimage 44, 489–501.

Stoodley, C.J., D’Mello, A.M., Ellegood, J., Jakkamsetti, V., Liu, P., Nebel, M.B., Gibson, J.M., Kelly, E., Meng, F., Cano, C.A., et al. (2017). Altered cerebellar connectivity in autism and cerebellar-mediated rescue of autism-related behaviors in mice. Nat. Neurosci. 20, 1744–1751.

Strick, P.L., Dum, R.P., and Fiez, J.A. (2009). Cerebellum and nonmotor function. Annu. Rev. Neurosci. 32, 413–434.

Su, P., Wang, H., Xia, J., Zhong, X., Hu, L., Li, Y., Li, Y., Ying, M., and Xu, F. (2019). Evaluation of retrograde labeling profiles of HSV1 H129 anterograde tracer. J. Chem. Neuroanat. 100, 101662.

Sugihara, I. (2018). Crus I in the Rodent Cerebellum: Its Homology to Crus I and II in the Primate Cerebellum and Its Anatomical Uniqueness Among Neighboring Lobules. Cerebellum 17, 49–55.

Sugihara, I., and Shinoda, Y. (2004). Molecular, topographic, and functional organization of the cerebellar cortex: a study with combined aldolase C and olivocerebellar labeling. J. Neurosci. 24, 8771–8785.

Suzuki, L., Coulon, P., Sabel-Goedknegt, E.H., and Ruigrok, T.J.H. (2012). Organization of cerebral projections to identified cerebellar zones in the posterior cerebellum of the rat. J. Neurosci. 32, 10854–10869.

Tervo, D.G.R., Hwang, B.-Y., Viswanathan, S., Gaj, T., Lavzin, M., Ritola, K.D., Lindo, S., Michael, S., Kuleshova, E., Ojala, D., et al. (2016). A Designer AAV Variant Permits Efficient Retrograde Access to Projection Neurons. Neuron 92, 372–382.

Teune, T.M., van der Burg, J., van der Moer, J., Voogd, J., and Ruigrok, T.J. (2000). Topography of cerebellar nuclear projections to the brain stem in the rat. Prog. Brain Res. 124, 141–172.

Thomson, A.M. (2010). Neocortical layer 6, a review. Front. Neuroanat. 4, 13.

Ugolini, G. (1992). Transneuronal transfer of herpes simplex virus type 1 (HSV 1) from mixed limb nerves to the CNS. I. Sequence of transfer from sensory, motor, and sympathetic nerve fibres to the spinal cord. J. Comp. Neurol. 326, 527–548.

Ugolini, G. (2010). Advances in viral transneuronal tracing. J. Neurosci. Methods 194, 2–20.

Ugolini, G., Kuypers, H.G., and Simmons, A. (1987). Retrograde transneuronal transfer of herpes simplex virus type 1 (HSV 1) from motoneurones. Brain Res. 422, 242–256.

Virtanen, P., Gommers, R., Oliphant, T.E., Haberland, M., Reddy, T., Cournapeau, D., Burovski, E., Peterson, P., Weckesser, W., Bright, J., et al. (2020). SciPy 1.0: fundamental algorithms for scientific computing in Python. Nat. Methods 17, 261–272.

Voogd, J., and Ruigrok, T.J.H. (2004). The organization of the corticonuclear and olivocerebellar climbing fiber projections to the rat cerebellar vermis: the congruence of projection zones and the zebrin pattern. J. Neurocytol. 33, 5–21.

van der Walt, S., Schönberger, J.L., Nunez-Iglesias, J., Boulogne, F., Warner, J.D., Yager, N., Gouillart, E., Yu, T., and scikit-image contributors (2014). scikit-image: image processing in Python. PeerJ 2, e453.

Wang, S.S.-H., Kloth, A.D., and Badura, A. (2014). The cerebellum, sensitive periods, and autism. Neuron 83, 518–532.

Waskom, M., Botvinnik, O., Hobson, P., Cole, J.B., Halchenko, Y., Hoyer, S., Miles, A., Augspurger, T., Yarkoni, T., Megies, T., et al. (2014). seaborn: v0.5.0 (November 2014).

Watabe-Uchida, M., Zhu, L., Ogawa, S.K., Vamanrao, A., and Uchida, N. (2012). Whole-brain mapping of direct inputs to midbrain dopamine neurons. Neuron 74, 858–873.

Wiesendanger, R., and Wiesendanger, M. (1982). The corticopontine system in the rat. I. Mapping of corticopontine neurons. J. Comp. Neurol. 208, 215–226.

Wijesinghe, R., Protti, D.A., and Camp, A.J. (2015). Vestibular Interactions in the Thalamus. Front. Neural Circuits 9, 79.

Winnubst, J., Bas, E., Ferreira, T.A., Wu, Z., Economo, M.N., Edson, P., Arthur, B.J., Bruns, C., Rokicki, K., Schauder, D., et al. (2019). Reconstruction of 1,000 Projection Neurons Reveals New Cell Types and Organization of Long-Range Connectivity in the Mouse Brain. Cell 179, 268–281.e13.

Wojaczynski, G.J., Engel, E.A., Steren, K.E., Enquist, L.W., and Patrick Card, J. (2015). The neuroinvasive profiles of H129 (herpes simplex virus type 1) recombinants with putative anterograde-only transneuronal spread properties. Brain Struct. Funct. 220, 1395–1420.

Yamamuro, K., Bicks, L.K., Leventhal, M.B., Kato, D., Im, S., Flanigan, M.E., Garkun, Y., Norman, K.J., Caro, K., Sadahiro, M., et al. (2020). A prefrontal-paraventricular thalamus circuit requires juvenile social experience to regulate adult sociability in mice. Nat. Neurosci. 23, 1240–1252.

Yao, B., Yang, X., and Zhu, S.-C. (2007). Introduction to a Large-Scale General Purpose Ground Truth Database: Methodology, Annotation Tool and Benchmarks. In Energy Minimization Methods in Computer Vision and Pattern Recognition, (Springer Berlin Heidelberg), pp. 169–183.

Yoo, A.B., Jette, M.A., and Grondona, M. (2003). SLURM: Simple Linux Utility for Resource Management. In Job Scheduling Strategies for Parallel Processing, (Springer Berlin Heidelberg), pp. 44–60.

Zemanick, M.C., Strick, P.L., and Dix, R.D. (1991). Direction of transneuronal transport of herpes simplex virus 1 in the primate motor system is strain-dependent. Proc. Natl. Acad. Sci. U. S. A. 88, 8048–8051.

