## Supplemental Tables and Figures for "Homologous organization of cerebellar pathways to sensory, motor, and associative forebrain"

| Structure | Timepoint | Number of brains | Edge erosion | Ventricular erosion |
|---|---|---|---|---|
| Brainstem (H129) | 28-36 hpi | 5 | 60 µm | 80 µm |
| Thalamus (H129) | 54 hpi | 23 | 60 µm | 80 µm |
| Neocortex (H129) | 80 hpi | 33 | 60 µm | 80 µm |
| Striatum (H129) | 80 hpi | 33 | 60 µm | 80 µm |
| Hypothalamus (H129) | 80 hpi | 31 | 60 µm | 160 µm |
| Neocortex (PRV) | 80 hpi | 25 | 60 µm | 80 µm |

| CNN | # of brains (# of volumes) | # of cells | Human-CNN concordance | Human-human concordance |
|---|---|---|---|---|
| H129 | 8 (44) | 3603 | F1: 0.864<br>Precision: 0.912<br>Recall: 0.821 | F1: 0.891<br>Precision: 0.947<br>Recall: 0.842<br>1091 cells annotated by both users |
| PRV | 7 (41) | 5119 | F1: 0.873<br>Precision: 0.833<br>Recall: 0.926 | F1: 0.886<br>Precision: 0.936<br>Recall: 0.841<br>1280 cells annotated by both users |

| # Synapses - Hours post injection (Publication) | Circuit investigated using HSV-H129 |
|---|---|
| 1 synapse - 48 hr<br>(McGovern et al., 2012a) | Airway related-sensory circuitry |
| 1 synapse - 48 hr<br>(Song et al., 2009) | White adipose tissue to CNS |
| 1 synapse - 50 hr<br>(Carta et al., 2019) | DCN -> midbrain |
| **2 synapses - 54 hr<br>(Our study)** | **Cerebellum -> DCN -> midbrain** |
| 2 synapses - 60 hr<br>(Badura et al., 2018) | Cerebellum -> DCN -> midbrain |
| 2 synapses - 72 hr<br>(Song et al., 2009) | White adipose tissue to CNS |
| 2 synapses - 72hr, 3 synapses at 96-144 hr<br>(McGovern et al., 2012b) | Tracheal sensory pathways<br>(peripheral nervous system -> CNS) |
| 3 synapses - 80 hr<br>(Badura et al., 2018) | Cerebellar -> DCN -> thalamus -> neocortex |
| 3 synapses - 96 hr<br>(Lo and Anderson, 2011) | Cerebellar output to neocortex |

**Table S1**. **Cohort details, cell detector training datasets, HSV-H129 used in literature.** Related to Figures 1, 2, 4, 5, 6 and 7.

| Thalamic Area | General function | Reference |
|---|---|---|
| Anteroventral | Spatial Memory | (Jankowski et al., 2013) |
| Central lateral | Emotional aspects of nociception | (Wang and Shyu, 2004) |
| Lateral dorsal | Somatosensory processing | (Bezdudnaya and Keller, 2008) |
| Lateral posterior | Visually-guided behavior | (Allen et al., 2016) |
| Lateral habenula | Reward Negative | (Matsumoto and Hikosaka, 2009) |
| Mediodorsal | Processing/integration of memory/cognition | (Mitchell and Chakraborty, 2013) |
| Medial habenula | Emotion-associated behavior | (Lee et al., 2019) |
| Parafascicular | Reversal Learning | (Brown et al., 2010) |
| Paraventricular | Emotional arousal, +/- behavioral mediation | (Kirouac, 2015; Yamamuro et al., 2020) |
| Posterior triangle | Nociception | (Gauriau and Bernard, 2004) |
| Posterior complex | Adjusting response to unexpected sensory input | (Casas-Torremocha et al., 2017) |
| Reticular | Cortical-based modulation of thalamus | (Lam and Sherman, 2010) |
| Reuniens | Hippocampal modulation | (Dolleman-van der Weel et al., 2019) |
| Submedial | Olfaction | (Tham et al., 2009) |
| VA-L | Memory/Spatial navigation & Motor | (Jankowski et al., 2013) |
| Ventral medial | Motor | (Starr and Summerhayes, 1983) |
| VPL | Sensory Body | (Vertes et al., 2015) |
| VPM | Sensory Face | (Vertes et al., 2015) |
| Ventral LGN | Visuomotor response & Circadian rhythms | (Harrington, 1997) |

**Supplementary 2. Thalamic target area function references.** Abbreviations: VA-L, ventral anterior-lateral; VPL, ventral posterolateral; VPM, ventral posteromedial; LGN, lateral geniculate nucleus. Related to Figures 2 and 3.

| Target | Injection | Primary antibody | Secondary antibody |
|---|---|---|---|
| c-Fos | rAAV1-CAG-FLEX-ArchT-GFP | 1:2000 Rabbit anti-c-Fos Synaptic Systems Cat. No. 226003 | 1:500 Donkey anti-Rabbit AlexaFluor 790 ThermoFisher A11374 |
| Anterograde thalamic timepoint (28-36 hpi) | H129-VC22 ($2.7x10^4$ to $8.0x10^4$ PFUs) | 1:1750 Rabbit anti-HSV Dako B011402-2 | 1:500 Donkey anti-Rabbit AlexaFluor 647 ThermoFisher A31573 |
| Anterograde thalamic timepoint (54 hpi) | H129-VC22 ($2.7x10^4$ to $8.0x10^4$ PFUs) | 1:1750 Rabbit anti-HSV Dako B011402-2 | 1:500 Donkey anti-Rabbit AlexaFluor 647 ThermoFisher A31573 |
| Anterograde neocortical timepoint (80 hpi) | H129-VC22 ($2.7x10^4$ to $8.0x10^4$ PFUs) | 1:350 Rabbit anti-HSV Dako B011402-2 | 1:250 Donkey anti-Rabbit AlexaFluor 647 ThermoFisher A31573 |
| Retrograde neocortical timepoint (28-50 hpi) | PRV-Bartha 152 ($6.0x10^4$ PFUs) | 1:750 Chicken anti-GFP Aves GFP-1020 | 1:400 Donkey anti-Chicken AlexaFluor 647 Jackson ImmunoResearch 703-606-155 |
| Retrograde neocortical timepoint (80 hpi) | PRV-Bartha 152 ($6.0x10^4$ PFUs) | 1:500 Chicken anti-GFP Aves GFP-1020 | 1:300 Donkey anti-Chicken AlexaFluor 647 Jackson ImmunoResearch 703-606-155 |

| Target | Cerebellar injection site | Structure | Mean ± std. dev. |
|---|---|---|---|
| Anterograde thalamic timepoint (53 hpi) | All injections | Sensory-motor | 2.5 ± 5.7 |
| | | Polymodal association | 1.0 ± 0.7 |
| | Vermis | Sensory-motor | 1.6 ± 2.4 |
| | | Polymodal association | 1.0 ± 0.6 |
| | Hemisphere | Sensory-motor | 3.5 ± 7.8 |
| | | Polymodal association | 1.2 ± 0.9 |
| Anterograde neocortical timepoint (80 hpi) | All injections | Frontal | 1.2 ± 0.5 |
| | | Medial | 1.2 ± 0.4 |
| | | Posterior | 1.0 ± 0.4 |
| | Vermis | Frontal | 1.2 ± 0.5 |
| | | Medial | 1.2 ± 0.5 |
| | | Posterior | 1.0 ± 0.5 |
| | Hemisphere | Frontal | 1.3 ± 0.4 |
| | | Medial | 1.2 ± 0.3 |
| | | Posterior | 1.2 ± 0.3 |
| Retrograde neocortical timepoint (80 hpi) | All injections | Frontal | 1.4 ± 0.6 |
| | | Medial | 3.2 ± 2.8 |
| | | Posterior | 1.7 ± 1.5 |
| | Vermis | Frontal | 1.2 ± 0.3 |
| | | Medial | 2.7 ± 3.1 |
| | | Posterior | 1.3 ± 0.8 |
| | Hemisphere | Frontal | 1.6 ± 0.7 |
| | | Medial | 3.9 ± 2.4 |
| | | Posterior | 2.2 ± 1.8 |

**Table S3. Injection and clearing details for transsynaptic and physiologic tracing from cerebellum, and projection ratios.** Contralateral-to-ipsilateral projection ratios for sub-regions in ascending and descending cerebellar pathways traced using H129-VC22 and PRV-Bartha. Front neocortical regions include infralimbic, prelimbic, anterior cingulate, orbital, frontal pole, gustatory, auditory, and visual cortex; medial regions include somatomotor and somatosensory cortex; posterior regions include retrosplenial, posterior parietal, temporal, perirhinal, and ectorhinal cortex. Ratios are shown as mean ± standard deviation across all brains in each cohort. Abbreviations: hpi, hours post-injection. Related to Figures 2, 4, 5 and 6.

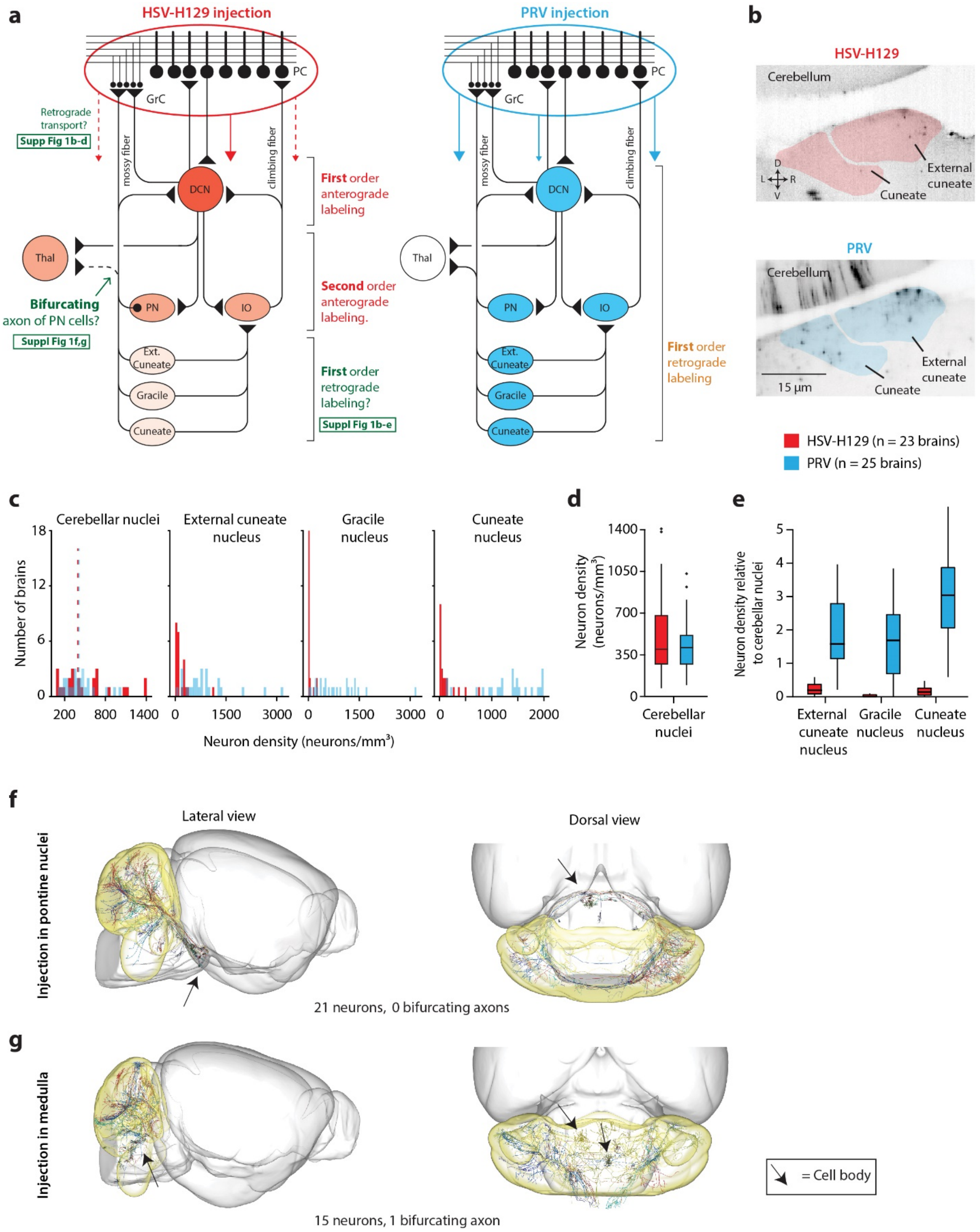


**Supplementary Figure 1**. HSV-H129 can be reliably used for anterograde tracing in the cerebellum. Related to Figures 1, 2, 6. (a) Summary circuit schematic depicting spread after cerebellar cortical injection of HSV-H129 (blue; Simplex injection) and PRV (orange; Lobule IX injection). *Left*, potential areas of retrograde spread after HSV-H129 injection are shown and control experiments performed to quantify retrograde spread are depicted. *Right*, PRV, an exclusively retrograde spreading virus is shown for comparison. Red and blue color intensity indicate intensity of expected labeling. (b) Example sections at disynaptic timepoints showing HSV-H129 (left; 54 hpi) and PRV (right; 80 hpi) in the cuneate and external cuneate nuclei. Any viral transport here is exclusively retrograde from the cerebellar cortex, as the dorsal column (cuneate, external cuneate and gracile) nuclei receive no monosynaptic anterograde projections from the deep cerebellar nuclei. (c) HSV-H129 and PRV disynaptic timepoint cell count density histograms in the cerebellar and dorsal column nuclei. Mean values shown as dotted lines for deep cerebellar nuclei. Retrograde:anterograde density ratios for HSV-H129 (n=23) and PRV (n=25) for deep cerebellar nuclei (d) and dorsal column nuclei (e). Densities in the dorsal column nuclei are divided by the deep cerebellar nuclei densities. Boxplots: center line represents median; box limits, upper and lower quartiles; whiskers, 1.5 times the interquartile range. Brainstem neurons, pontine (f) and medulla (g), that send axons into the cerebellum do not send axons to extracerebellar regions. To determine if retrogradely-transported HSV-H129 could spread extracerebellarly via axons, the Mouselight database was surveyed for brainstem somas with at least one axonal cerebellar projection. The query revealed 36 traced neurons. Of those neurons only one had projections both into the deep cerebellar nuclei, as well as extracerebellar axons. The remaining axons had exclusive projections back to the cerebellum. Somata in pons with at least one cerebellar axon: AA1003, AA1004, AA1005, AA1007, AA1008, AA1009, AA1010, AA1028, AA1029, AA1052, AA1053, AA1057, AA1060, AA1071, AA1072, AA1073, AA1074, AA1076, AA1087, AA1091, AA1092. Somata in medulla with at least one cerebellar axon: AA0503, AA0922, AA0950, AA0951, AA0953, AA1062, AA1063, AA1064, AA1068, AA1070, AA1077, AA1083, AA1084, AA1085, AA1093.

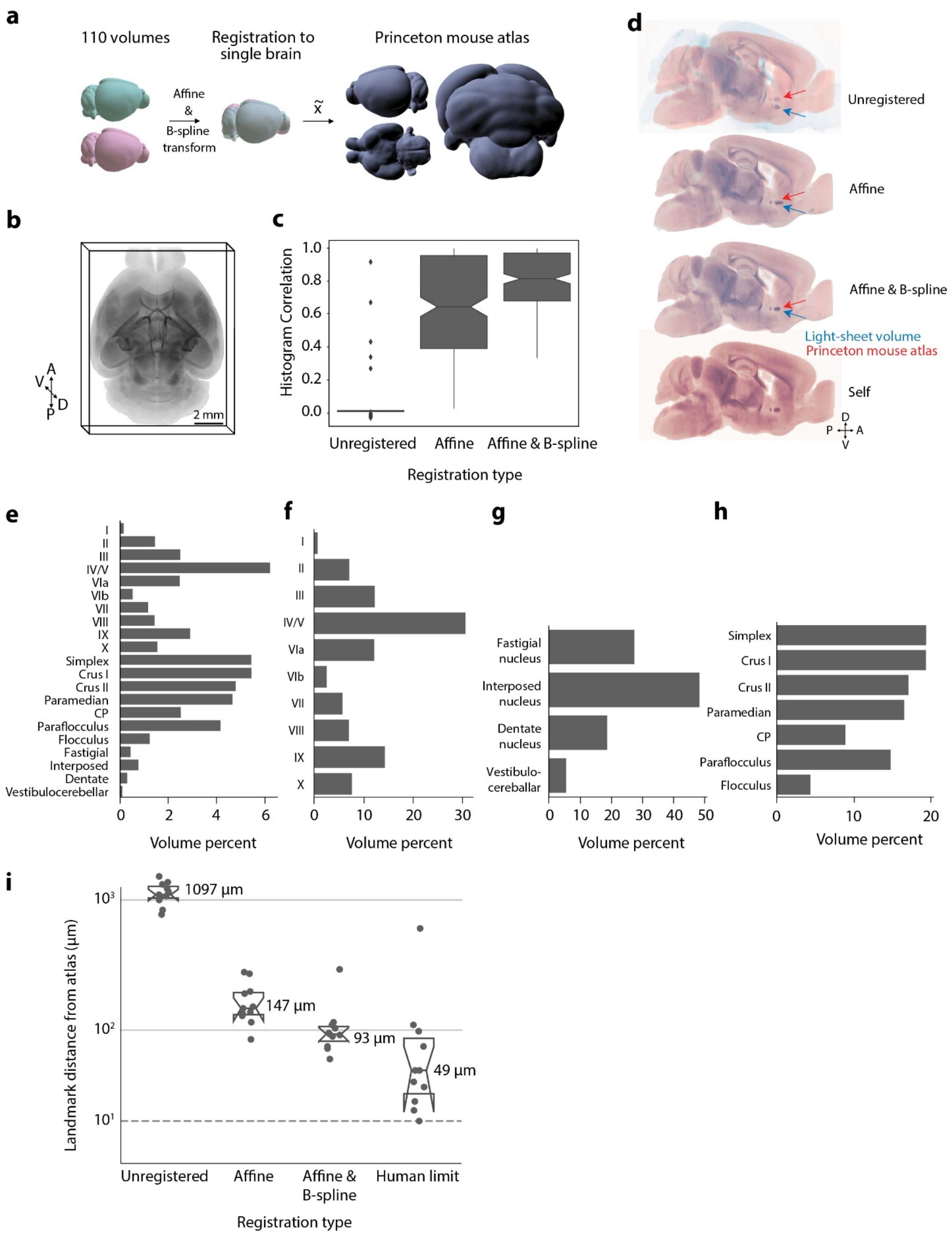


**Supplementary Figure 2**. The Princeton mouse atlas, a light-sheet volumetric atlas with a complete cerebellum. Related to Figure 1. (a) Schematic depicting atlas generation. Mouse brains cleared using iDISCO+ (n=110) were imaged using a light-sheet microscope and were downsized to 20 μm/voxel. A single volume was selected and the other brains registered to it. The median XYZ voxel was then used from the resulting metabrain. (b) Three-dimensional projection rendering ("3D project" function, ImageJ) of the light-sheet atlas. (c) Histogram correlations demonstrate human-independent improvement in volumetric alignment. Pearson's correlations (scipy.stats) were calculated using normalized histograms (bins=300) for unregistered (r=.005, p=.856, medians), affine (r=0.518, p=4.94 x 10^-22), and affine & B-spline (r=0.712, p=1.26 x 10^-47) registered volumes (n=224) with the PMA. (d) Color-blind friendly version demonstrating landmark alignment example. Percent contributions of substructures to cerebellar volume in the PMA. (e) Cerebellar substructure percent volumes. Bar plot depicts volumes as percentage of gross cerebellar volumes in the PMA. Relative volume percentages of substructures in the vermis (f), deep cerebellar nuclei (g), and hemispheres (h) are also shown. Abbreviations: CP, copula pyramidis. (i) Landmark euclidean distance quantification demonstrates registration performance. Users (n=11), blinded to each volume's condition, annotated a total of 69 complementary points, across four brains, in unregistered (two identical volumes, human precision), affine, affine & B-spline. Three-dimensional euclidean distances were determined. Points are median user performance per condition and numbers displayed are median distances across users. Dashed horizontal line depicts single voxel distance (20 μm).

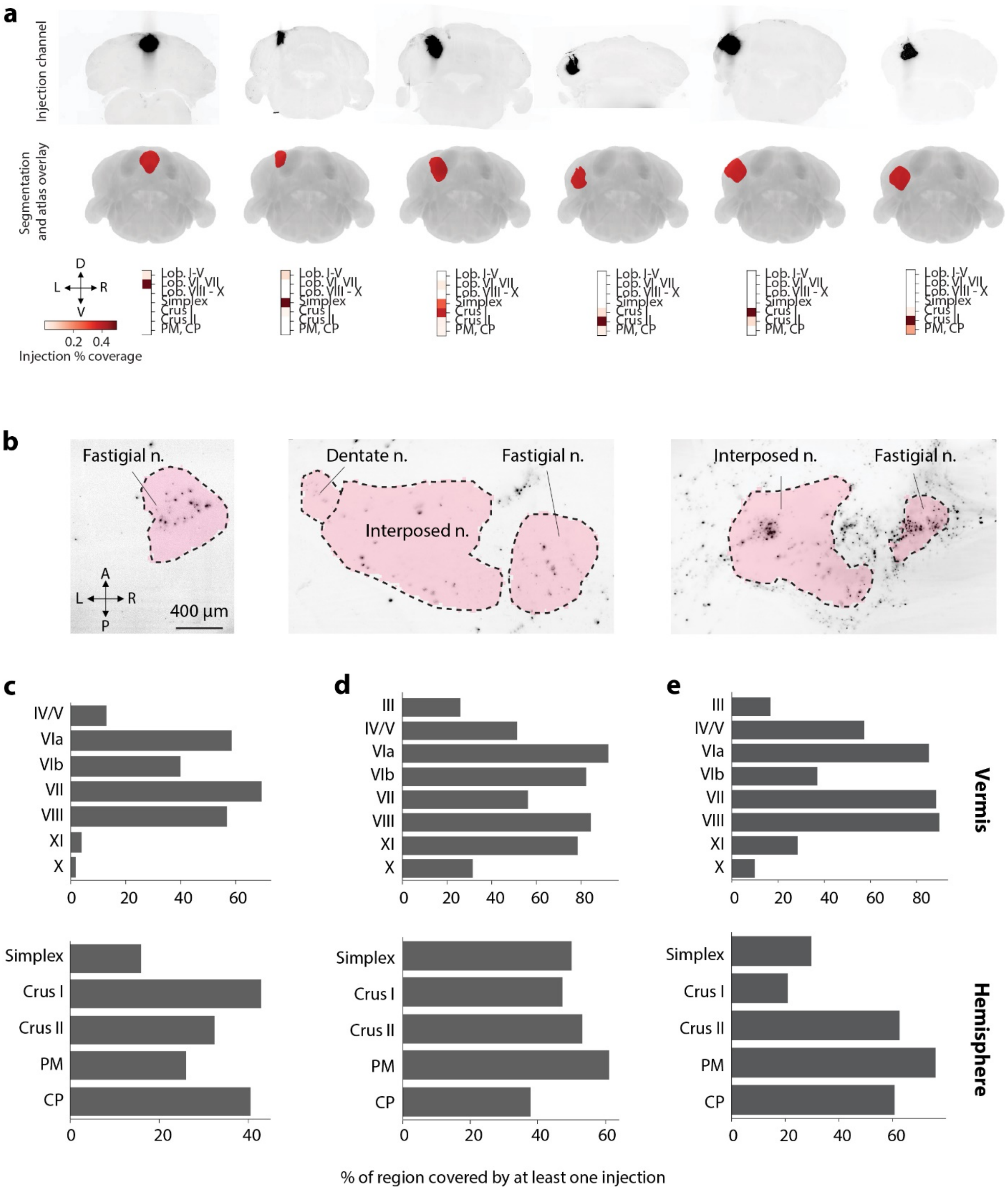


**Supplementary Figure 3**. Example injection site mapping and deep cerebellar nuclear HSV-H129 spread. Related to Figures 2, 4, 6. (a) Injection site segmentation and mapping onto the Princeton Mouse Atlas. Cholera toxin conjugated to fluorophore allows for visualization of the cerebellar injection region. Coronal maximum intensity projections of injection volumes are shown after registration to the Princeton Mouse atlas. After segmentation, the injection volume is overlaid onto the Princeton Mouse atlas. (b) Example horizontal sections from cleared mouse brains showing HSV-H129 labeling in deep cerebellar nuclei. *Left,* bilateral fastigial nuclei labeling at 36 hpi; *Middle*, left fastigial and interposed nuclei labeling at 53 hpi; *Right*, right fastigial and interposed nuclei labeling at 80 hpi. Pink shading shows boundaries of cerebellar nuclei. Abbreviations: Lob., Lobule, L, left; R, right; n., nuclei. Graphs show percent of cerebellar cortical region covered by at least 1 injection after automated injection site quantification of H129-VC22 and PRV injected brains. Brains used in the H129 thalamic cohort (n=23) (c), the H129 neocortical cohort (n=33) (d), and the PRV neocortical cohort (n=25) (e).

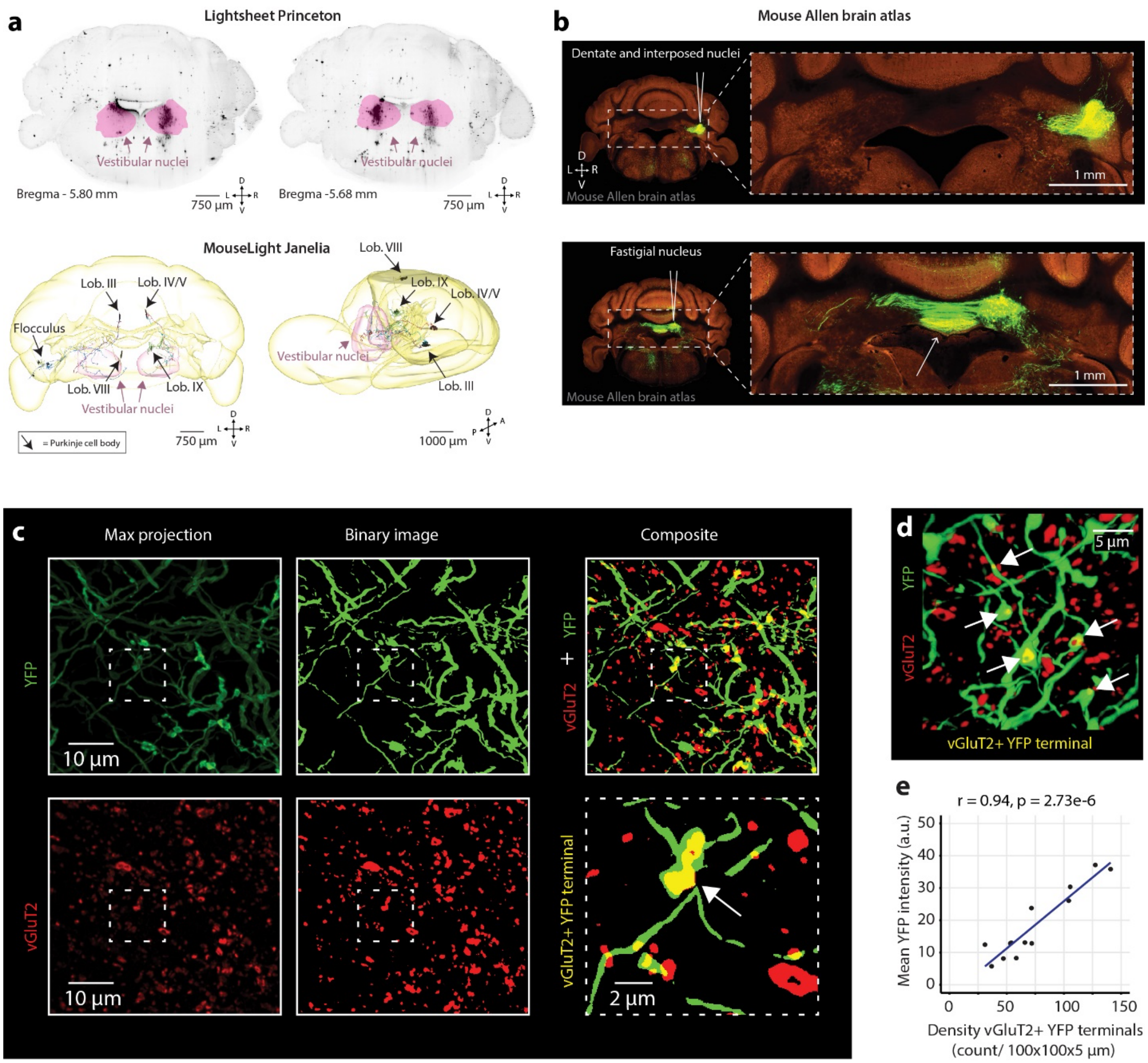


**Supplementary Figure 4**. Purkinje neurons projecting to vestibular nuclei and DCN injection quantification validation. Related to Figures 2, 3. (a) Purkinje neurons projecting to vestibular nuclei. To determine the lobular location of Purkinje neurons that directly project to the vestibular nuclei, Mouselight was queried for neurons with somata in the cerebellum and at least one axonal projection to the vestibular nuclei. Nine Purkinje neurons met this criteria. Of them 6 were nonflocculonodular neurons. The 9 mouselight neurons meeting criteria of cerebellar cortex soma with vestibular axon are: AA1022, AA0986, AA0985, AA0983, AA0975, AA0972, AA0971, AA0963, AA0962. (b) Cerebellar stereotactic AAV injection site revealed successful targeting of deep cerebellar nuclei. Coronal section after a unilateral cerebellar injection with dentate and interposed nuclear expression. Axons were visible exiting from nuclei. Coronal section after a unilateral cerebellar injection (different animal) demonstrating fastigial nuclear expression. Axons were visualized exiting bilaterally from the cerebellum. (c) Validation of YFP intensity as an estimator for the number of axon varicosities. Image processing and segmentation pipeline. Raw images were first background subtracted. Images were binarized and particle analysis was used to quantify connected pixels as individual varicosities. In total we quantified 12 image stacks. (d) Three-dimensional rendering showing colocalization (arrows) of vGluT2 (red) and YFP terminals (green). (e) Correlation of number of vGluT2+ terminals versus YFP density within individual ROIs.

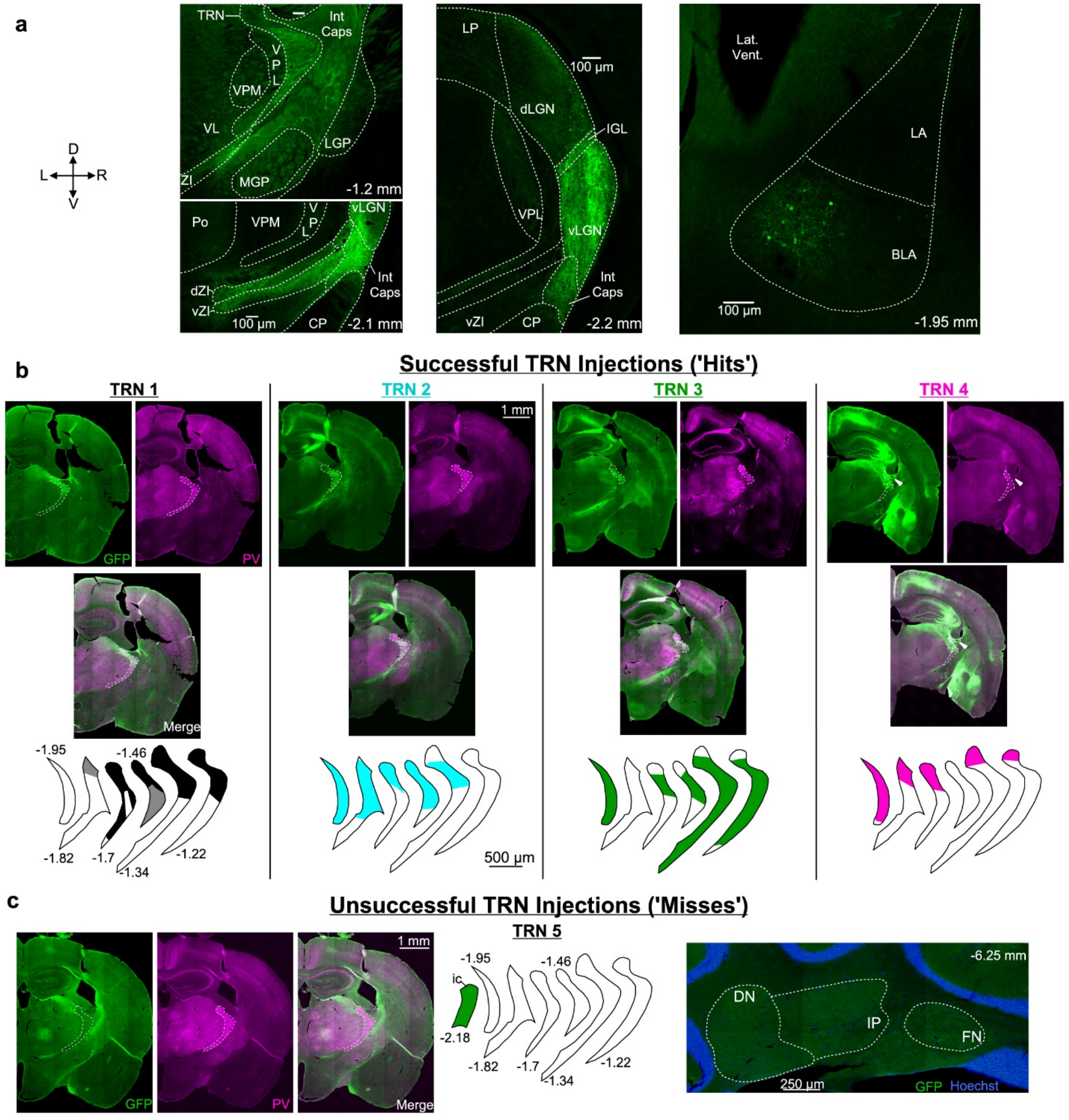


**d**

| Experiment | TRN | Cerebellar Nuclei: Dentate | Cerebellar Nuclei: Interpositus | Cerebellar Nuclei: Fastigial | Vestib. Nuclei: Medial/VCB | Thalamic Nuclei: ZI | vLGN | dLGN | VAL | VPL | VPM | LD | VL | CM | CL | Cortex: Som | Vis | Aud | Hip | BLA | Int Caps | Caud Put |
|---|---|---|---|---|---|---|---|---|---|---|---|---|---|---|---|---|---|---|---|---|---|---|
| **Successful 'Hit'** | | | | | | | | | | | | | | | | | | | | | | |
| TRN1 | ++ | ++ | + | - | + | + | +++ | ++ | - | + | - | + | - | - | - | + | + | + | - | ++ | + | +/- |
| TRN2 | +++ | ++ | + | + | + | ++ | +++ | ++ | - | + | + | + | - | - | - | + | ++ | + | ++ | + | + | +/- |
| TRN3 | +++ | ++ | + | + | + | +++ | +++ | + | - | + | + | + | - | - | - | + | ++ | ++ | ++ | ++ | + | +/- |
| TRN4* | ++ | ++ | + | + | + | ++ | +++ | + | - | + | - | + | - | - | - | + | +++ | ++ | +++ | +++ | + | + |
| **Unsuccessful 'Miss'** | | | | | | | | | | | | | | | | | | | | | | |
| TRN5 | - | - | - | - | - | + | +/- | - | - | - | - | - | - | - | - | - | ++ | + | + | + | + | +/- |

+++ Strong GFP   ++ Moderate GFP   + Minor GFP   - No GFP

**Supplementary Figure 5**. Assessment of AAVrg-GFP expression accuracy and complete results. Related to Figure 3. (a) Confocal images of additional areas expressing GFP after injection of AAVrg-GFP into TRN. In addition to cerebellar nuclei, viral injections targeting TRN labeled cells in internal capsule, ZI, dLGN, and basolateral amygdala with minor labeling in ventral posterior and laterodorsal thalamic nuclei. Labeling was also seen in LV pyramidal neurons in somatosensory, visual, and auditory cortex likely due to infection of axons passing through TRN. Terminal labeling in vLGN is consistent with infection of visual L5 corticothalamic neurons passing through TRN. Minor labeling was observed in caudate putamen which is near the TRN and may have taken up virus. Labeling in hippocampus was typically observed due to deposit of virus upon insertion/removal of the injection needle. Distance relative to bregma is provided. (b) Epifluorescence microscopy image of AAVrg-GFP (green; left) and parvalbumin (PV; magenta; right) expression in TRN (outlined in white) for all four injections that successfully targeted or 'hit' TRN. The portions of TRN expressing GFP (coronal plane) throughout the anterior-posterior axis of TRN are shaded in color with the color corresponding to the same experiments in Figure 4. Complete absence of shading/labeling at one plane indicates that section was not examined histologically. Location of each plane relative to bregma is provided for TRN1 and applies to all. Each experiment labeled separate regions of TRN: TRN1 – anteriorodorsal and middle; TRN2 – posteriodorsal, middle; TRN3 – ventral; TRN4 – dorsal. *TRN4 labeled the dorsal TRN, but also infected stria terminalis (arrow) to produce more substantial labeling in amygdala than other injections. The more infection of the more dorsal TRN corresponded to retrograde labeling of neurons in the posterior interpositus. (c) Characterization of an unsuccessful 'missed' injection that did not induce GFP expression at any location in the TRN nor in any cerebellar nuclei. This injection location was deemed to be in the internal capsule adjacent to the TRN. Distance relative to bregma is provided. (d) Summary table of major sites of GFP expression or regions known to receive direct projections from cerebellar nuclei. GFP expression was evaluated as strong (+++), moderate (++), minor (+), or none (-) and examples can be found for corresponding expeirments and brain regions in raw data shown in a-c. Abbreviations: 4V, fourth ventricle; BLA, basolateral amygdala; CM, centromedial thalamus; CL, centrolateral thalamus; CP, cerebal peduncle; dLGN, dorsal lateral geniculate nucleus, DN, dentate nucleus; dZI, dorsal zona incerta; FN, fastigial nucleus; GFP, green fluorescent protein; Hip, hippocampus; ic, internal capsule; Int Caps, internal capsule; IP, interpositus nucleus; LA, lateral amygdala; LD, laterodorsal thalamus; LP, lateral posterior thalamus; LGP; lateral globus pallidus; M, motor cortex; MGP, medial globus pallidus; Po, posterior thalamus; PV, parvalbumin; S, somatosensory cortex; TRN, thalamic reticular nucleus; VL, ventrolateral thalamus; vLGN, ventral lateral geniculate nucleus; VN, vestibulocerebellar nucleus; VPM, ventroposterior lateral thalamus; VPM, ventroposterior medial thalamus; vZI, ventral zona incerta; ZI, zona incerta.

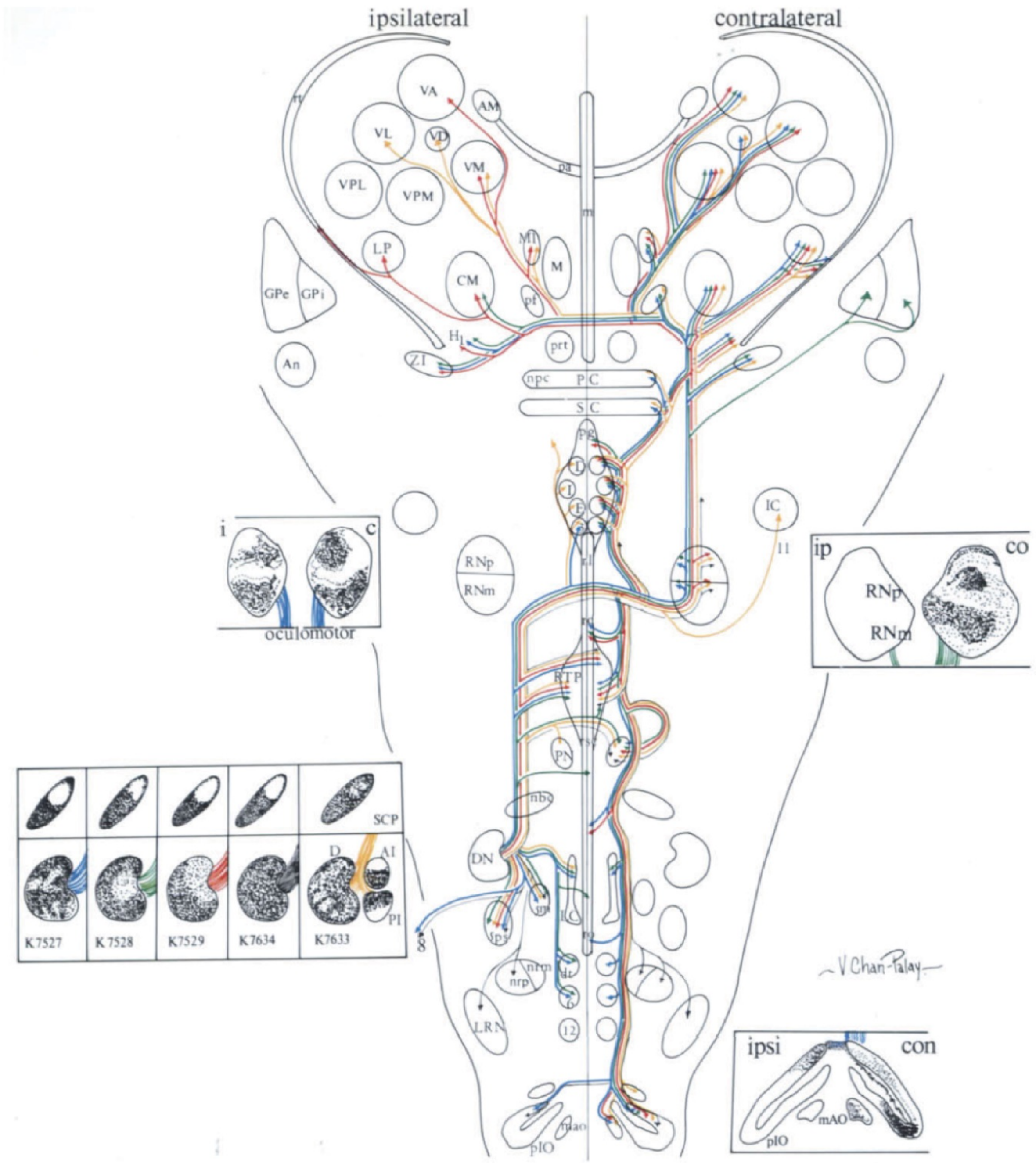


**Supplementary Figure 6**. Summary of course and terminations of dentatofugal fibers labeled after precisely localized injections of 35S-methionine into the lateral nucleus of rats in cases K 7527, K 7528, K 7529, K 7634, and into the lateral and interpositus nuclei in K 7633. Related to Figures 2 and 3. The regions of labeled neurons in the cerebellar nuclei at the injection site are shown in the insets (lower left) with their indentation number and color code. The brainstem and thalamus are shown in horizontal view with the locations of various structures labeled according to the listed abbreviations. The course and projections gathered from each case are summarized with the appropriate color code for each. The remaining insets who the topography of label concentrated in the superior cerebellar peduncle (SCP) after each experiment and in projection sites reconstruction in horizontal views of the ipsilateral and contralateral oculomotor nuclei (left), red nuclei (upper right), and inferior olivary complex (lower right). *Figure from Chan-Palay, V. (2013). Cerebellar Dentate Nucleus: Organization, Cytology and Transmitters. Springer.*

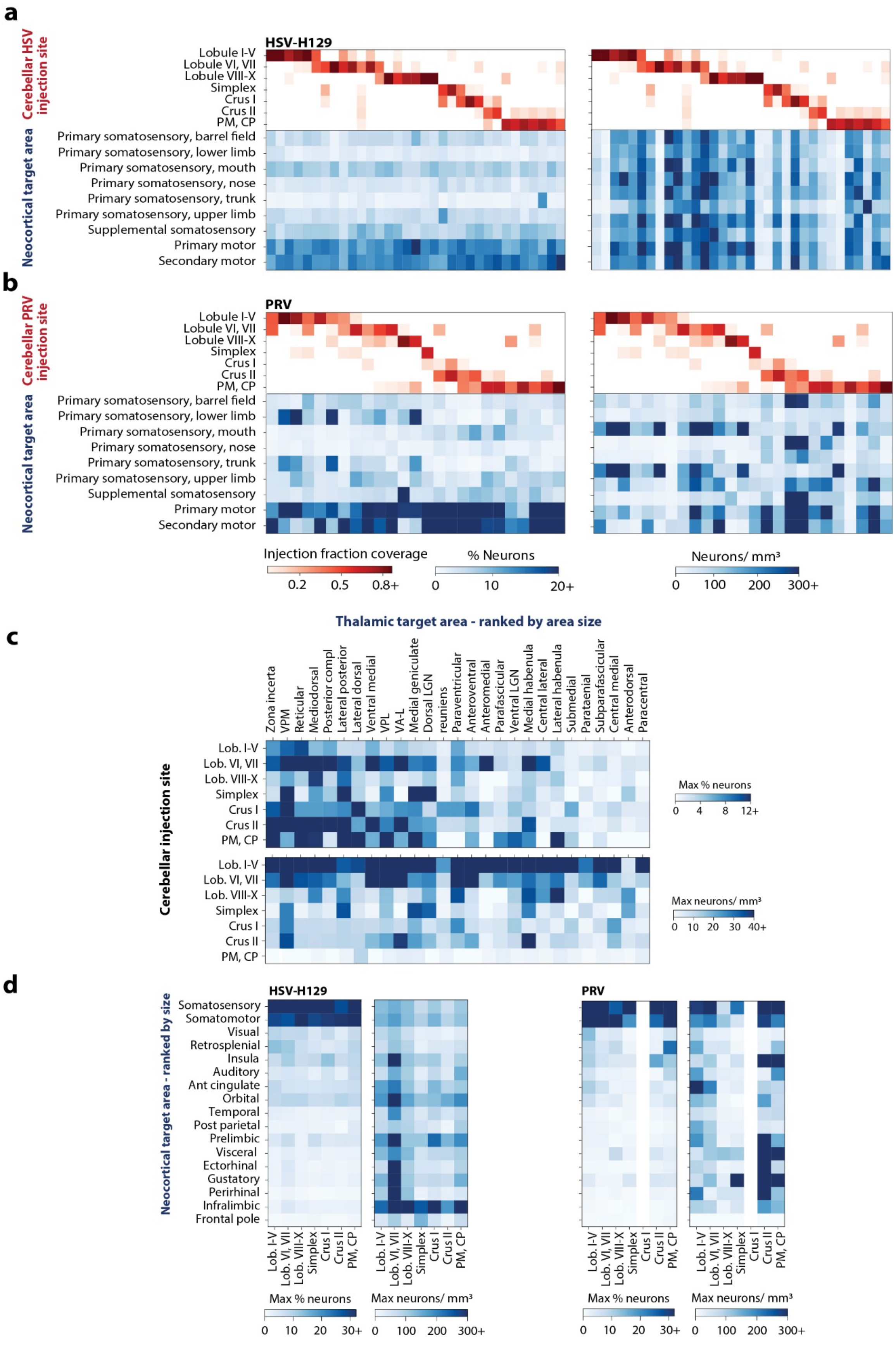
a
HSV-H129
Cerebellar HSV injection site
Lobule I-V
Lobule VI, VII
Lobule VIII-X
Simplex
Crus I
Crus II
PM, CP
Neocortical target area
Primary somatosensory, barrel field
Primary somatosensory, lower limb
Primary somatosensory, mouth
Primary somatosensory, nose
Primary somatosensory, trunk
Primary somatosensory, upper limb
Supplemental somatosensory
Primary motor
Secondary motor
b
PRV
Cerebellar PRV injection site
Lobule I-V
Lobule VI, VII
Lobule VIII-X
Simplex
Crus I
Crus II
PM, CP
Neocortical target area
Primary somatosensory, barrel field
Primary somatosensory, lower limb
Primary somatosensory, mouth
Primary somatosensory, nose
Primary somatosensory, trunk
Primary somatosensory, upper limb
Supplemental somatosensory
Primary motor
Secondary motor
Injection fraction coverage
0.2 0.5 0.8+
% Neurons
0 10 20+
Neurons/ mm³
0 100 200 300+
c
Thalamic target area - ranked by area size
Zona incerta
VPM
Reticular
Mediodorsal
Posterior compl
Lateral posterior
Lateral dorsal
Ventral medial
VPL
VA-L
Medial geniculate
Dorsal LGN
reuniens
Paraventricular
Anteroventral
Anteromedial
Parafascicular
Ventral LGN
Medial habenula
Central lateral
Lateral habenula
Submedial
Parataenial
Subparafascicular
Central medial
Anterodorsal
Paracentral
Cerebellar injection site
Lob. I-V
Lob. VI, VII
Lob. VIII-X
Simplex
Crus I
Crus II
PM, CP
Max % neurons
0 4 8 12+
Max neurons/ mm³
0 10 20 30 40+
d
HSV-H129
PRV
Neocortical target area - ranked by size
Somatosensory
Somatomotor
Visual
Retrosplenial
Insula
Auditory
Ant cingulate
Orbital
Temporal
Post parietal
Prelimbic
Visceral
Ectorhinal
Gustatory
Perirhinal
Infralimbic
Frontal pole
Lob. I-V
Lob. VI, VII
Lob. VIII-X
Simplex
Crus I
Crus II
PM, CP
Max % neurons
0 10 20 30+
Max neurons/ mm³
0 100 200 300+

**Supplementary Figure 7**. Sensorimotor subregions in HSV and PRV neocortical tracing and maximum projections for each timepoint. Related to Figures 2, 4 and 6. (a-b) Quantification of motor and sensory cortical subregions of transsynaptic tracing studies. (a) HSV subregional quantification. *Lef*t, percent of total motor/sensory HSV labeling by each structure. Right: density of HSV labeling by subregions. (b) PRV subregional quantification. *Left*, percent of total motor/sensory PRV labeling by each structure. *Right*, density of PRV labeling by subregions. (c-d) Maximum percent neurons and density projections for each timepoint. Injections were then pooled by taking the maximum subregion value across all injections of a given cerebellar location. (c) HSV-H129 tracing that the thalamic timepoint (54 hpi) (d). *Left*, HSV-H129 tracing at the neocortical timepoint (80 hpi) and *right*, PRV tracing at the neocortical timepoint (80 hpi).

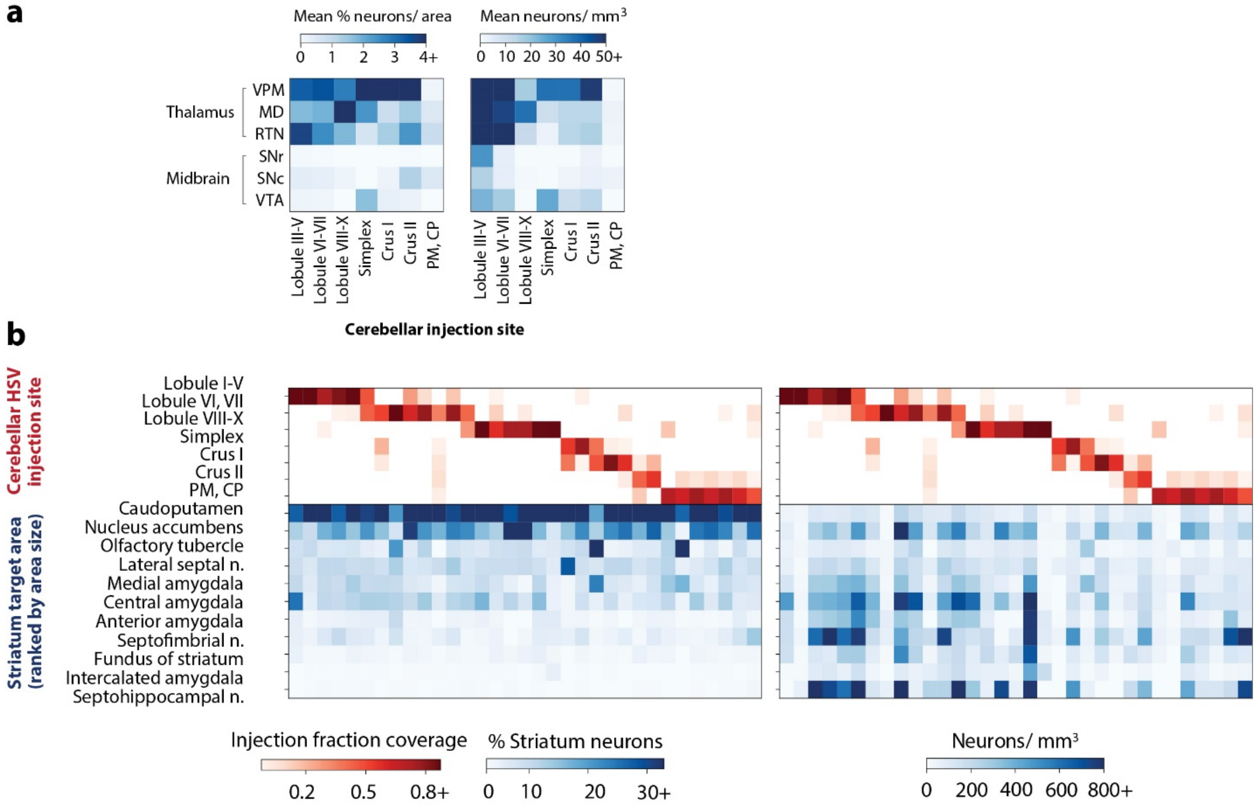


**Supplementary Figure 8**. Striatal projections of the cerebellum. Related to Figure 2 and 4. (a) Cerebellar paths to ventral tegmental area are weaker than thalamic projections at the 54 hour timepoint. *Left*, mean percentage of total thalamic and midbrain neurons in each region grouped by primary injection site. *Right,* mean density of neurons in each region grouped by primary injection site. The top 3 most labeled thalamic regions and selected midbrain regions are shown. (b) Cerebellar projections to the contralateral striatum at the neocortical timepoint. *Left,* percent of total labeled striatum neurons. *Right*, neuron density. Abbreviations: n., nucleus.

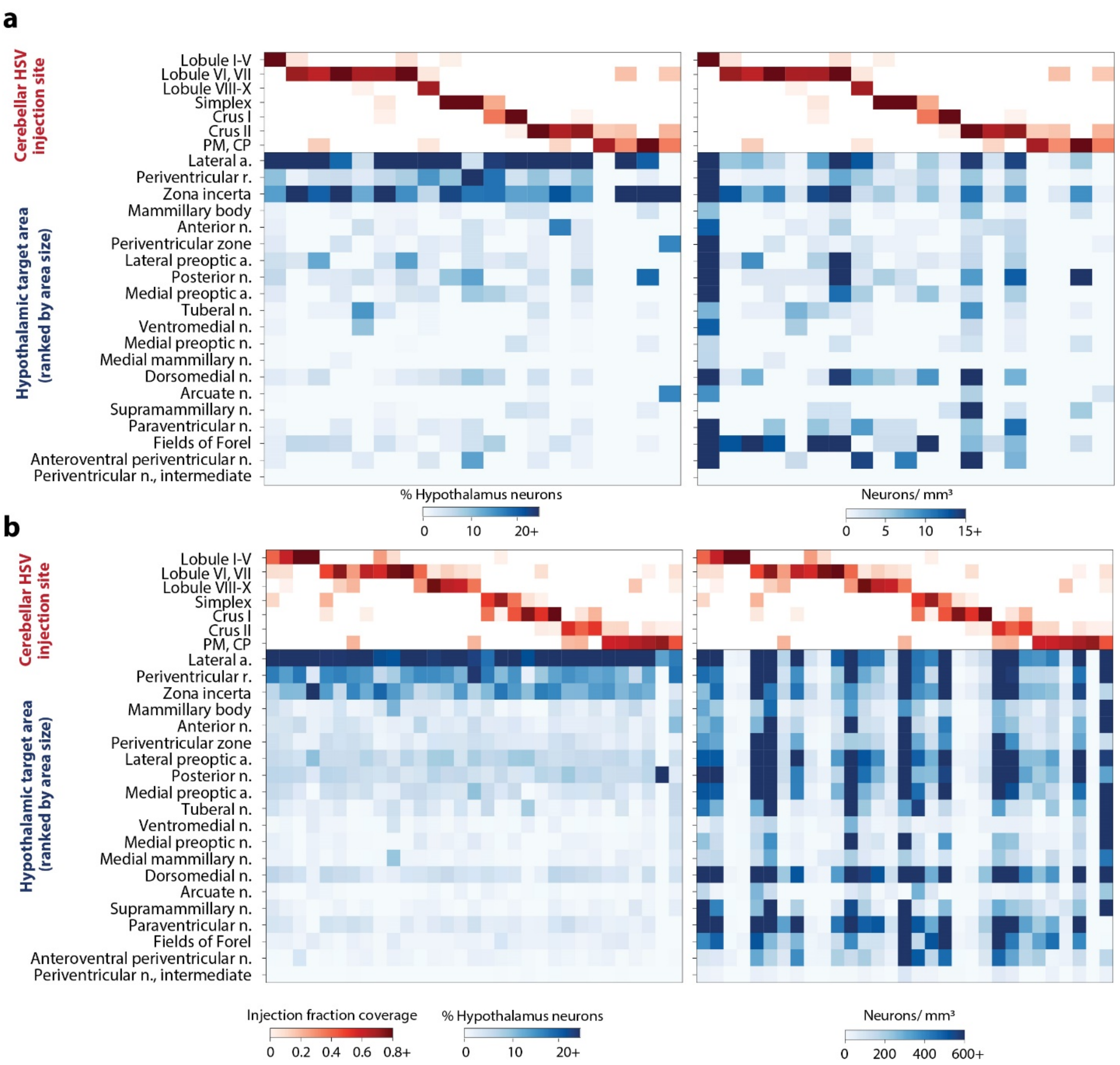


**Supplementary Figure 9.** Cerebellar output to bilateral hypothalamus at the thalamic (a) and neocortical timepoints (b). Related to Figures 2 and 4. *Left column,* percent of total labeled striatum neurons. *Right column*, neuron density. To minimize false positives, areas around ventricles were eroded by 160 µm removing some volume from the hypothalamic areas around ventral portions of the third ventricle. Hypothalamic regions are sorted from largest to smallest in descending order. Abbreviations: a., area; n., nucleus; r., region.

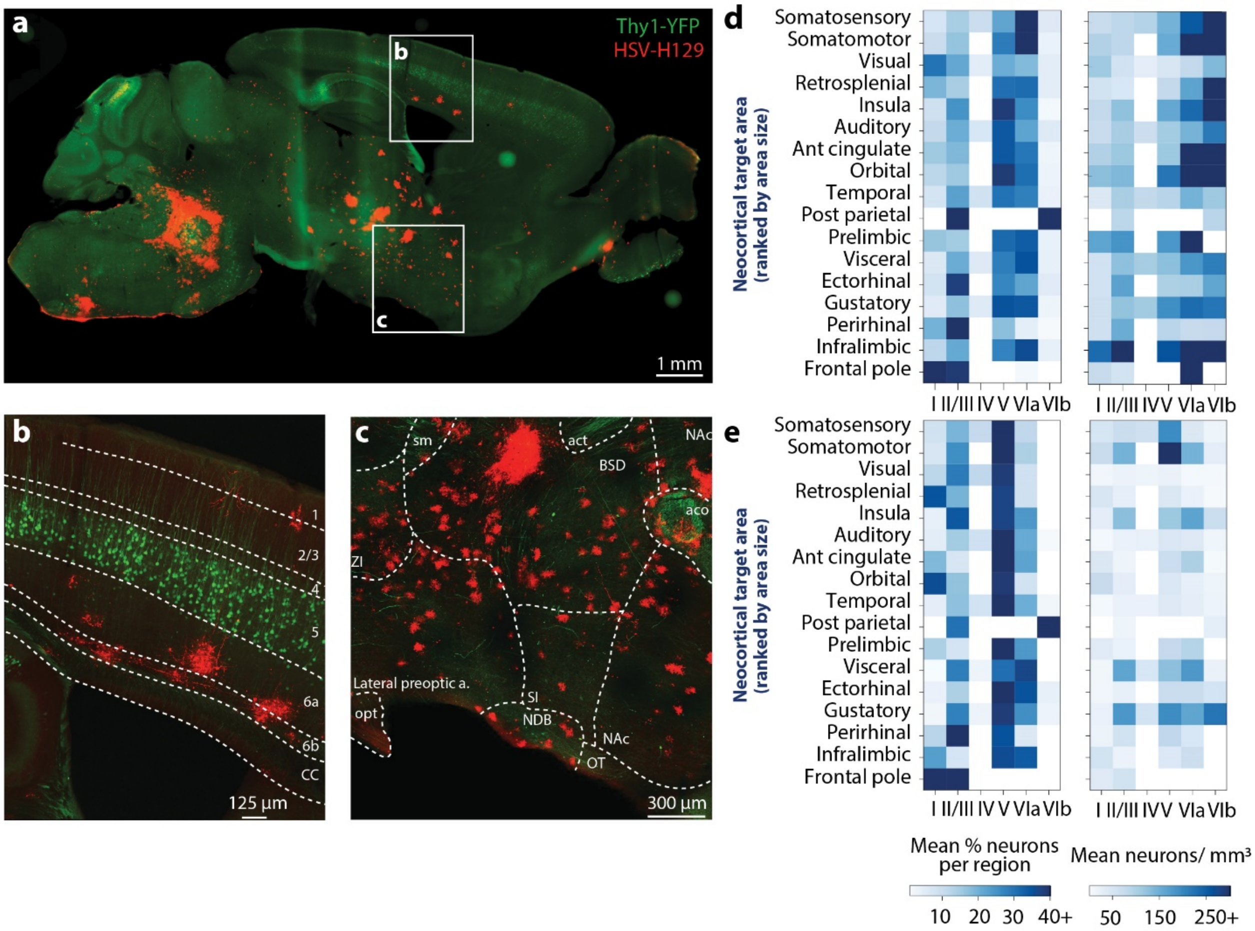


**Supplementary Figure 10**. HSV-H129 injection in cerebellar cortex reveals deep neocortical and nucleus accumbens labeling. Related to Figures 4, 5 and 7. (a) Cerebellar output connections labeled using the anterograde tracer H129 at 80 hpi. HSV-H129 (red) was injected into lobule VI of Thy1-YFP (green) mice. 50 µm section. (b) Confocal image of neocortical region shows HSV-H129 viral label in layer VI of the neocortex, separate from the layer V labelled in Thy1-YFP mouse. Cortical layers are outlined. (c) Confocal image shows viral labeling in nucleus accumbens, pallidal and hypothalamic areas. Abbreviations: a., area; aco, anterior commissure, olfactory limb; act, anterior commissure, temporal limb; BST, bed nuclei of the stria terminalis; CC, corpus callosum; n., nucleus; NAc, nucleus accumbens, NDB, diagonal band nucleus; opt, optic tract; OT, olfactory tubercle; SI substantia innominata; sm, stria medullaris; Thal., thalamus; ZI, zona incerta. (d-e) Quantification of neocortical transsynaptic layer labeling after HSV (d) and PRV (e) cerebellar injections. *Left column*, mean percent count across cerebellar injection regions by neocortical region and layer. *Right column*, mean density of labeling across cerebellar injection regions by neocortical region and layer.

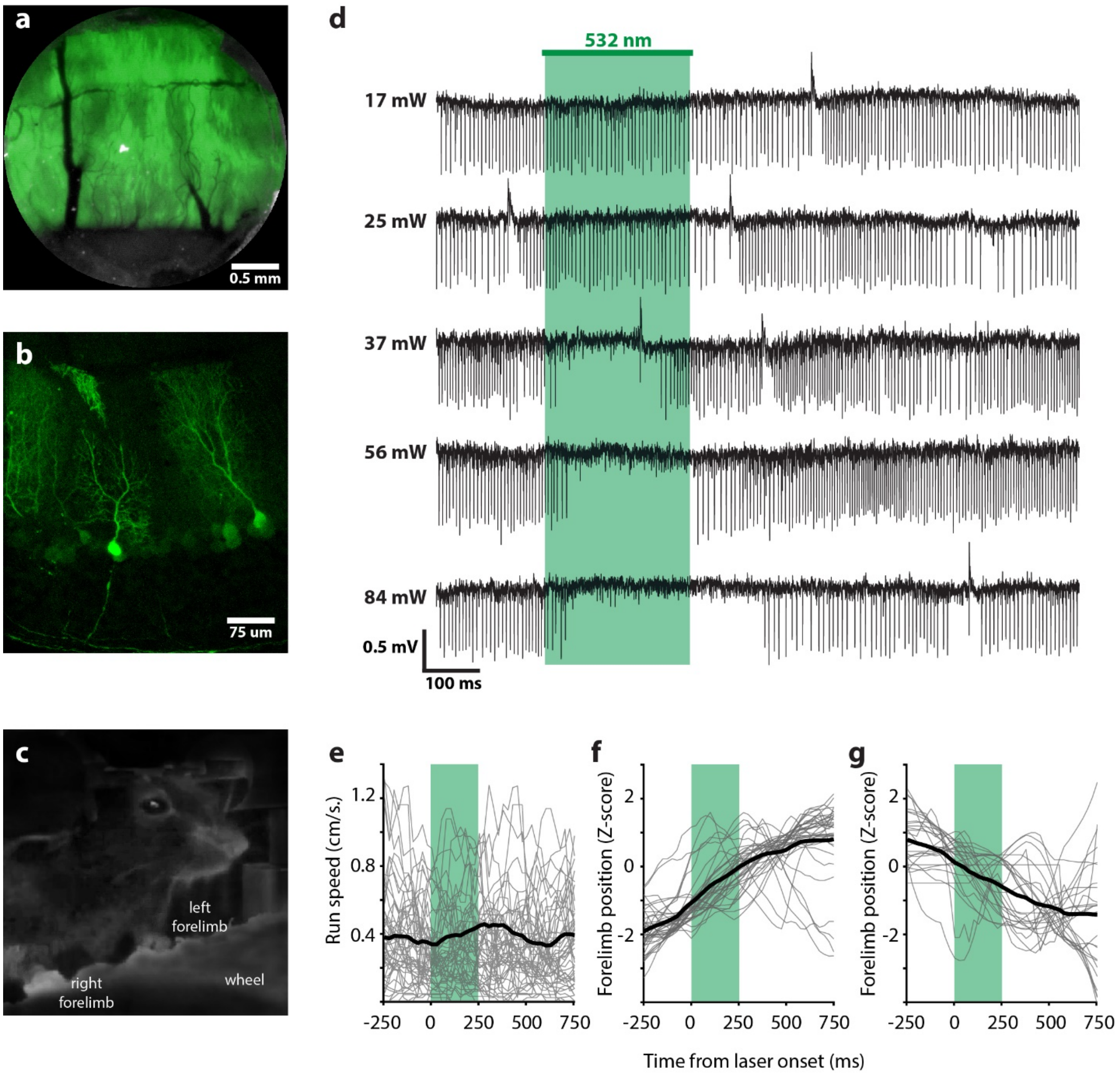


**Supplementary Figure 11**. Inactivation of Purkinje cells during light activation of ArchT specifically expressed in L7-Cre+/- mice. Related to Figure 7. (a) In vivo epifluorescence through cranial window used for stimulation of cranial window 4 weeks post-injection shows prominent expression of ArchT-GFP at Lobule VI. (b) Parasagittal section of cerebellar cortex from L7-cre +/- mouse showing Purkinje cell ArchT-GFP expression. (c) Head-fixed mouse on treadmill during stimulation. (d) Representative single unit recording from Purkinje cell responding to 250 ms light application at 1 Hz. Inactivation during light is gradually increased with increasing light intensities. Note the sustained block of spontaneous spikes after offset of light pulses at 84 mW. Treadmill speed (e), forward-moving right forelimb (f), backward-moving right forelimb (g) traces before, during (green box) and after stimulation.

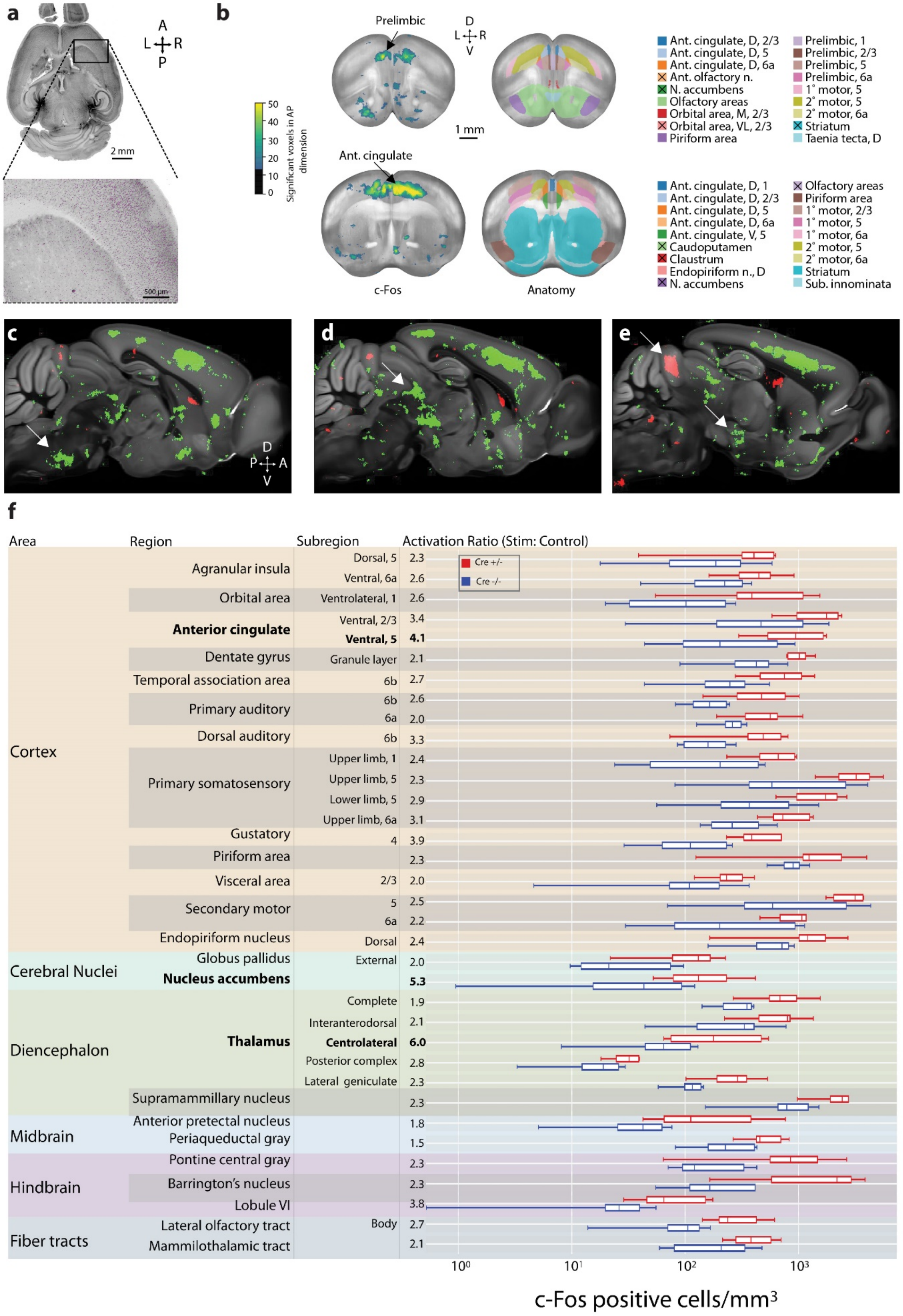
a
A
L R
P
2 mm
500 µm
b
Prelimbic
D
L R
V
1 mm
Ant. cingulate
Significant voxels in AP dimension
50
40
30
20
10
0
c-Fos
Anatomy
Ant. cingulate, D, 2/3
Ant. cingulate, D, 5
Ant. cingulate, D, 6a
Ant. olfactory n.
N. accumbens
Olfactory areas
Orbital area, M, 2/3
Orbital area, VL, 2/3
Piriform area
Prelimbic, 1
Prelimbic, 2/3
Prelimbic, 5
Prelimbic, 6a
1° motor, 5
2° motor, 5
2° motor, 6a
Striatum
Taenia tecta, D
Ant. cingulate, D, 1
Ant. cingulate, D, 2/3
Ant. cingulate, D, 5
Ant. cingulate, D, 6a
Ant. cingulate, V, 5
Caudoputamen
Claustrum
Endopiriform n., D
N. accumbens
Olfactory areas
Piriform area
1° motor, 2/3
1° motor, 5
1° motor, 6a
2° motor, 5
2° motor, 6a
Striatum
Sub. innominata
c
D
P A
V
d
e
f
Area
Region
Subregion
Activation Ratio (Stim: Control)
Cre +/-
Cre -/-
Cortex
Agranular insula
Dorsal, 5 2.3
Ventral, 6a 2.6
Orbital area
Ventrolateral, 1 2.6
Anterior cingulate
Ventral, 2/3 3.4
Ventral, 5 4.1
Dentate gyrus
Granule layer 2.1
Temporal association area
6b 2.7
Primary auditory
6b 2.6
6a 2.0
Dorsal auditory
6b 3.3
Primary somatosensory
Upper limb, 1 2.4
Upper limb, 5 2.3
Lower limb, 5 2.9
Upper limb, 6a 3.1
Gustatory
4 3.9
Piriform area
2.3
Visceral area
2/3 2.0
Secondary motor
5 2.5
6a 2.2
Endopiriform nucleus
Dorsal 2.4
Cerebral Nuclei
Globus pallidus
External 2.0
Nucleus accumbens
5.3
Diencephalon
Thalamus
Complete 1.9
Interanterodorsal 2.1
Centrolateral 6.0
Posterior complex 2.8
Lateral geniculate 2.3
Supramammillary nucleus
2.3
Midbrain
Anterior pretectal nucleus
1.8
Periaqueductal gray
1.5
Hindbrain
Pontine central gray
2.3
Barrington's nucleus
2.3
Lobule VI
3.8
Fiber tracts
Lateral olfactory tract
Body 2.7
Mammilothalamic tract
2.1
10⁰
10¹
10²
10³
c-Fos positive cells/mm³

**Supplementary Figure 12**. A brain-wide nonmotor network traced from the cerebellum. (a) ClearMap automatically quantifies c-Fos expression. Related to Figure 7. A horizontal image of a whole mouse brain with c-Fos antibody labeling (left) and overlay of c-Fos (gray) with c-Fos positive cells detected using ClearMap (purple) are shown. 132 μm maximum intensity projection. (b) Cortical areas show increased c-Fos cell counts after cerebellar optogenetic perturbation. Coronal maximum intensity projections (left) across 1 mm of tissue corresponding to Princeton mouse atlas planes 100-150 (top) and 150-200 (bottom) after 375 μm spherical voxelization. Complementary sections (right) with anatomical labels of 18 structures with the largest number of significant voxels. Structures with the largest AP span are shown when they overlap. Black X's in legend denote structures not shown due to overlap. Abbreviations: 1˚, primary; 2˚, secondary; ant, anterior; AP, anteroposterior, D, dorsal; L, lateral, M, medial; n., nucleus; SS, somatosensory; sub, substantia; V, ventral. (c-e) c-Fos p-value maps comparing brain regions activated by cerebellar optogenetic perturbation (green) vs. controls (red) reveal patterns of activation in pontine nuclei (c), midbrain (d), and superior colliculi (upper arrow) and hypothalamus (lower arrow) (e). White arrows in each panel indicate named regions of interest. Significant voxels (green or red) are shown overlaid on the Allen Brain Atlas template brain. (f) Lobule VI Purkinje cell inhibition leads to strong activity increases in nonmotor areas including the anterior cingulate, nucleus accumbens and centrolateral nucleus of thalamus. Structures listed have a Mann-Whitney p-value < 0.05. In boxplots, center line represents median; box limits, upper and lower quartiles; whiskers, 1.5 times the interquartile range.

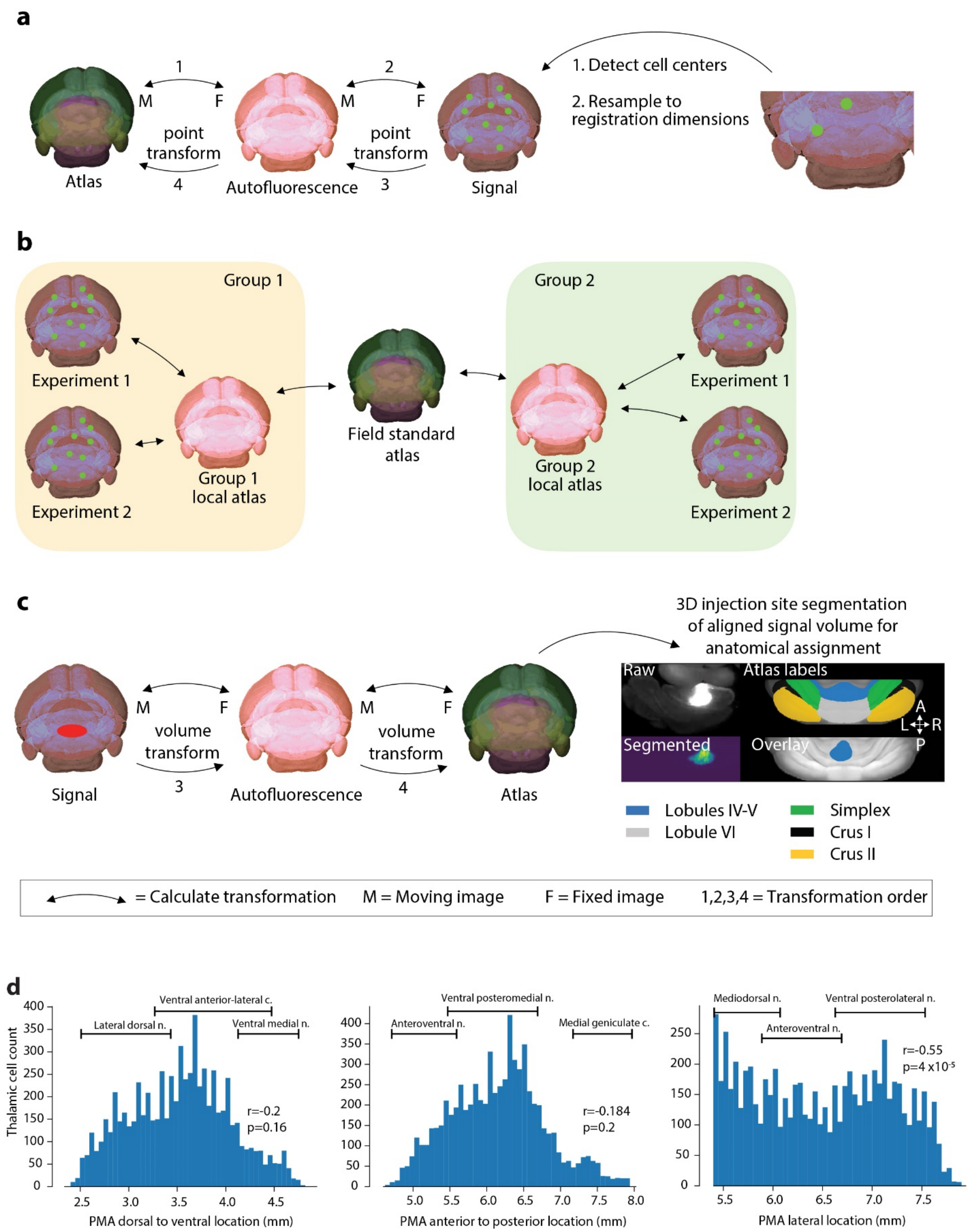


**Supplementary Figure 13**. Anatomical registration framework for automated volumetric analysis that facilitates data commutability. Related to Figures 1 and 2. (a) Cell center anatomical assignments require multiple transformations. Cell center anatomical assignment requires learning mapping between atlas and signal space. The optimal approach is determining the transformations of atlas (moving) to autofluorescence (fixed) and autofluorescence to signal space. Detected cell centers that have been resampled to registration volume dimensions can be point transformed and anatomically assigned. (b) A template solution for anatomical commutability between groups. Schematic depicting a solution of balancing considerations for project specific atlas requirements while maintaining consistency with field standards. Groups independently generate local atlases with all features required in their respective projections. Each experiment can accurately be registered with the local atlas. Each group then determines transformation between their local atlas and the field standard, allowing for anatomical commutability across groups. Line with arrows represents determining a transformation between two volumes. (c) Injection site segmentation and alignment process. Injection site anatomical assignment is most efficiently done by mapping signal space (moving) with atlas space (fixed). After the signal image transformation into atlas space, the injection site can be easily segmented and voxels anatomically assigned. F, fixed image; M, moving image. The lower half of B shows an example of segmenting a raw injection site and anatomically assigning to vermal cerebellar lobules IV/V and VI. (d) Thalamic cell count as a function of Princeton Mouse Atlas location. Cell counts as a function of location in each axis: (*left*) dorsal to ventral, (*middle*) anterior to posterior, (*right*) midline to lateral location are shown. The horizontal axis range indicates the full extent of the thalamus in PMA space. In the lateral location plot, the left boundary represents the thalamic midline. Pearson's correlation coefficients and p values were calculated using cell counts by thalamic location (n=50 bins). (scipy.stats.pearsonr). Abbreviations: c., complex; f, fixing volume; m, moving volume; n., nucleus.

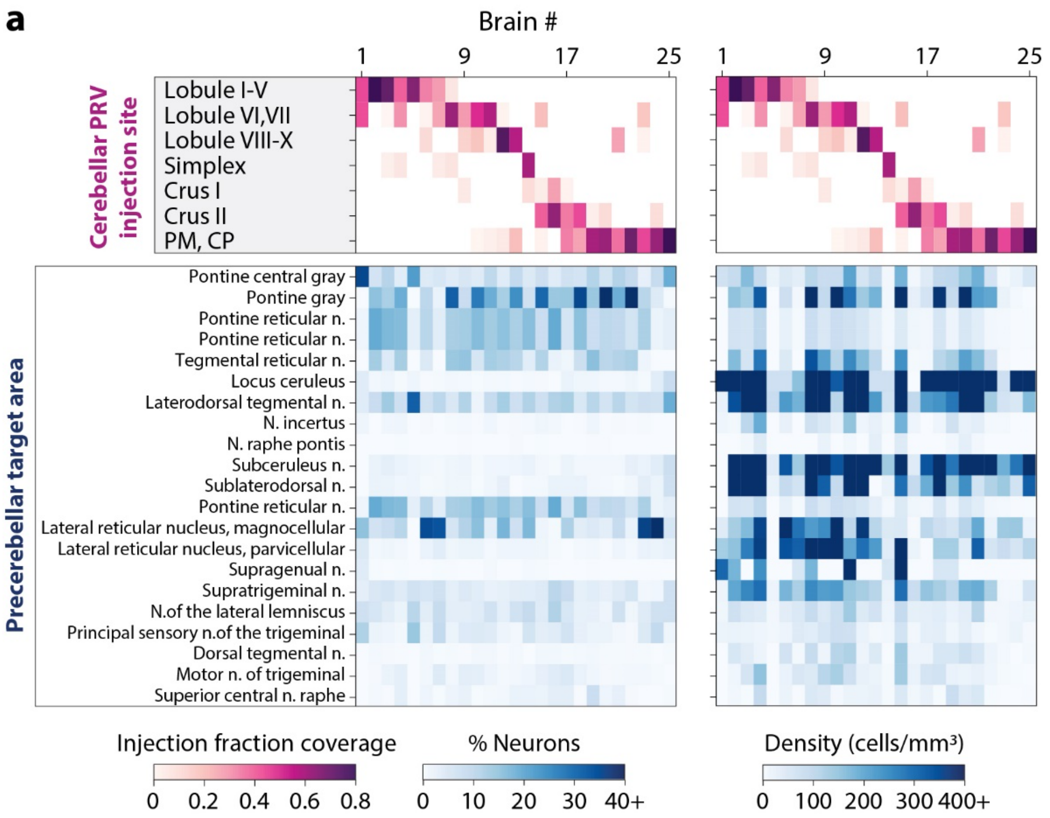
a
Brain #
1
9
17
25
Cerebellar PRV injection site
Lobule I-V
Lobule VI,VII
Lobule VIII-X
Simplex
Crus I
Crus II
PM, CP
Precerebellar target area
Pontine central gray
Pontine gray
Pontine reticular n.
Pontine reticular n.
Tegmental reticular n.
Locus ceruleus
Laterodorsal tegmental n.
N. incertus
N. raphe pontis
Subceruleus n.
Sublaterodorsal n.
Pontine reticular n.
Lateral reticular nucleus, magnocellular
Lateral reticular nucleus, parvicellular
Supragenual n.
Supratrigeminal n.
N.of the lateral lemniscus
Principal sensory n.of the trigeminal
Dorsal tegmental n.
Motor n. of trigeminal
Superior central n. raphe
Injection fraction coverage
0 0.2 0.4 0.6 0.8
% Neurons
0 10 20 30 40+
Density (cells/mm³)
0 100 200 300 400+

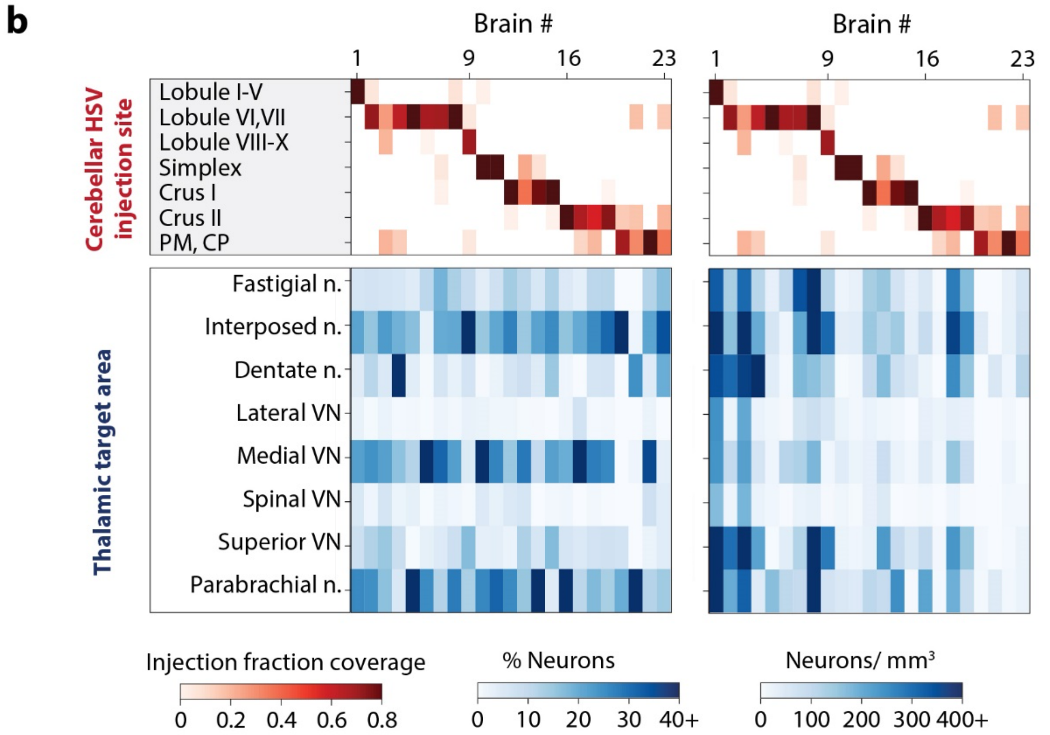
b
Brain #
1
9
16
23
Cerebellar HSV injection site
Lobule I-V
Lobule VI,VII
Lobule VIII-X
Simplex
Crus I
Crus II
PM, CP
Thalamic target area
Fastigial n.
Interposed n.
Dentate n.
Lateral VN
Medial VN
Spinal VN
Superior VN
Parabrachial n.
Injection fraction coverage
0 0.2 0.4 0.6 0.8
% Neurons
0 10 20 30 40+
Neurons/ mm³
0 100 200 300 400+

**Supplementary Figure 14**. Cerebellar monosynaptic connections. Related to Figures 2 and 6. (a) Precerebellar inputs. *Left*, fraction of neurons in each precerebellar target area for each injection site at the disynaptic PRV timepoint, 80 hours post-injection. Percentage fractions (blue) were calculated by dividing the number of neurons detected in each area by the total number of neurons detected in all precerebellar nuclei combined. Injection coverage fractions are shown in pink. *Right*, density of neurons in each precerebellar area across cerebellar injection sites. (b) Monosynaptic targets of the cerebellar cortex. *Left*, fraction of neurons in each precerebellar target area for each injection site at the disynaptic HSV-H129 timepoint, 54 hours post-injection. Percentage fractions (blue) were calculated by dividing the number of neurons detected in each area by the total number of neurons detected in all precerebellar nuclei combined. Injection coverage fractions are shown in pink. The parabrachial nucleus density may include the superior cerebellar peduncle (brachium conjunctivum), around which it wraps, and which is difficult to distinguish after tissue clearing. *Right*, density of neurons in each precerebellar area across cerebellar injection sites. Abbreviations: c., complex; n., nucleus; VN, vestibular nucleus.

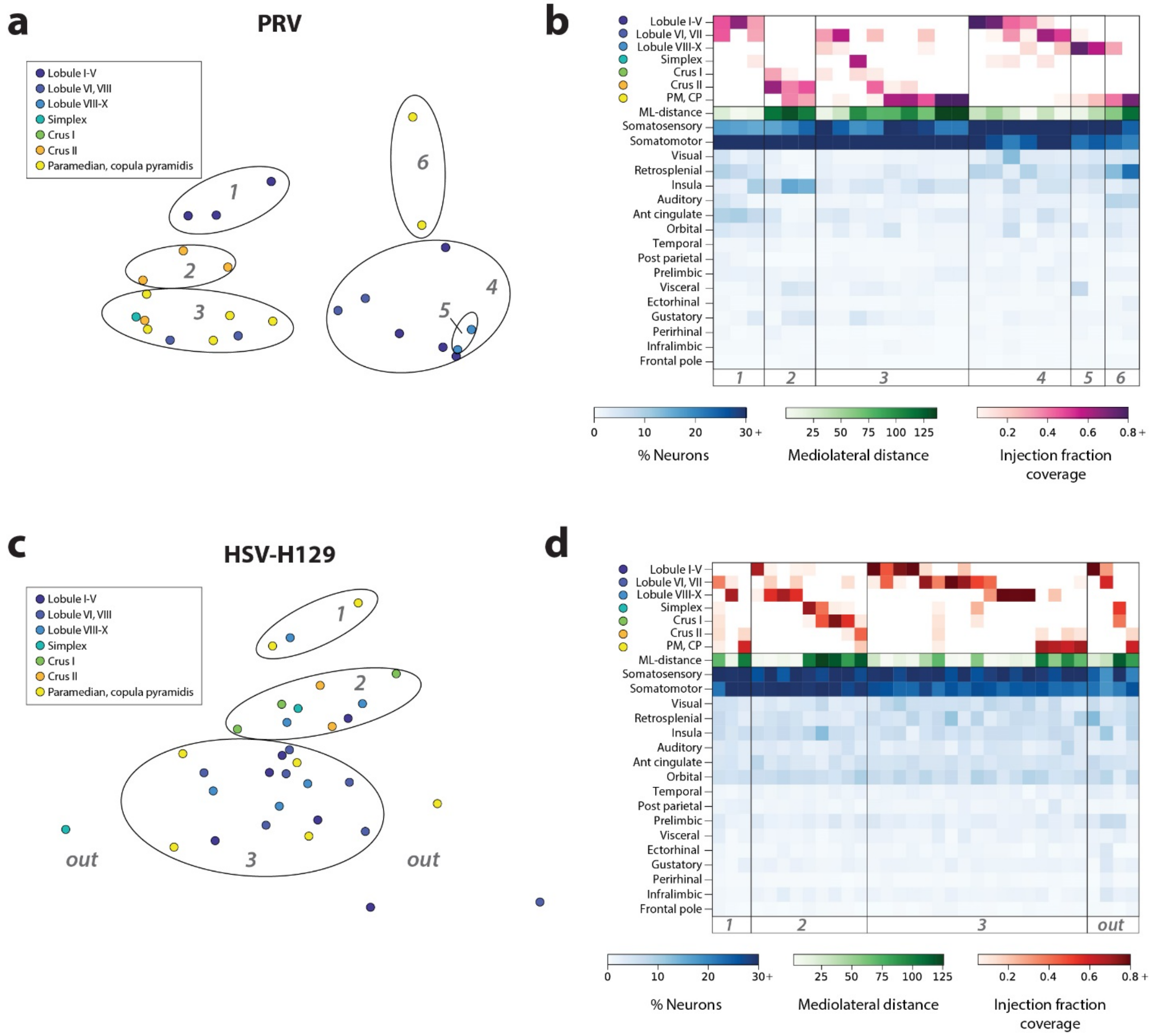


**Supplementary Figure 15**. Multidimensional scaling (MDS) of projection patterns in the neocortex generated from transsynaptic tracing. Related to Figures 4 and 6. Scatterplots were generated using as inputs the percentage of neurons in all neocortical regions. (a) PRV tracing at the disynaptic retrograde timepoint, 80 hours post-injection. The fill color indicates the lobule with the largest injection volume as determined by CTB co-injected with virus. (b) Heatmaps of the neocortical expression pattern, arranged according to groups of brains identified from MDS. The lobule volume is indicated in red and the mediolateral distance (ML-distance) is indicated in green. (c) MDS of HSV-H129 tracing at the trisynaptic anterograde timepoint, 80 hours post-injection. (d) Heatmaps of the neocortical expression pattern, arranged according to groups identified from MDS.
